# Dose-response relationship of pulmonary disorders by inhalation exposure to cross-linked water-soluble acrylic acid polymers in F344 rats

**DOI:** 10.1101/2021.12.26.474221

**Authors:** Tomoki Takeda, Shotaro Yamano, Yuko Goto, Shigeyuki Hirai, Yusuke Furukawa, Yoshinori Kikuchi, Kyohei Misumi, Masaaki Suzuki, Kenji Takanobu, Hideki Senoh, Misae Saito, Hitomi Kondo, George Daghlian, Young Kwon Hong, Yasuhiro Yoshimatsu, Masanori Hirashima, Yoichiro Kobashi, Kenzo Okamoto, Takumi Kishimoto, Yumi Umeda

## Abstract

**Background:** In Japan, six workers handling cross-linked water-soluble acrylic acid polymer (CWAAP) at a chemical plant suffered from lung diseases, including fibrosis, interstitial pneumonia, emphysema, and pneumothorax. We recently demonstrated that inhalation of CWAAP-A, one type of CWAAP, causes pulmonary disorders in rats. It is important to investigate dose-response relationships and recoverability from exposure to CWAAPs for establishing occupational health guidelines, such as setting threshold limit value for CWAAPs in the workplace.

**Methods:** Male and female F344 rats were exposed to 0.3, 1, 3, or 10 mg/m^3^ CWAAP-A for 6 hours/day, 5 days/week for 13 weeks using a whole-body inhalation exposure system. At 1 hour, 4 weeks, and 13 weeks after the last exposure the rats were euthanized and blood, bronchoalveolar lavage fluid, and all tissues including lungs and mediastinal lymph nodes were collected and subjected to biological and histopathological analyses. In a second experiment, male rats were pre-treated with clodronate liposome or polymorphonuclear leukocyte-neutralizing antibody to deplete macrophages or neutrophils, respectively, and exposed to CWAAP-A for 6 hours/day for 2 days.

**Results:** CWAAP-A exposure damaged only the alveoli. The lowest observed adverse effect concentration (LOAEC) was 1 mg/m^3^ and the no observed adverse effect concentration (NOAEC) was 0.3 mg/m^3^. Rats of both sexes were able to recover from the tissue damage caused by 13 weeks exposure to 1 mg/m^3^ CWAAP-A. In contrast, tissue damage caused by exposure to 3 and 10 mg/m^3^ was irreversible due to the development of interstitial lung lesions. There was a gender difference in the recovery from CWAAP-A induced pulmonary disorders, with females recovering less than males. Finally, acute lung effects caused by CWAAP-A were significantly reduced by depletion of alveolar macrophages.

**Conclusions:** Pulmonary damage caused by inhalation exposure to CWAAP-A was dose-dependent, specific to the lung and lymph nodes, and acute lung damage was ameliorated by depleting macrophages in the lungs. CWAAP-A had both a LOAEC and a NOAEC, and tissue damage caused by exposure to 1 mg/m^3^ CWAAP-A was reversible: recovery in female rats was less than for males. These findings indicate that concentration limits for CWAAPs in the workplace can be determined.

## Background

Cross-linked water-soluble acrylic acid polymers (CWAAPs) are used worldwide as a thickening agent to increase viscosity and sol-gel stability. CWAAPs are used in such as cosmetics and pharmaceuticals owing to their low potential for skin and eye irritation. However, recently in Japan, six workers who were handling CWAAPs at a chemical plant producing these resins suffered from lung diseases, including fibrosis, interstitial pneumonia, emphysema, and pneumothorax [1]. Notably, five of these workers had a work history of only about two years. Based on the results obtained from a clinical research project on this case [2, 3], in April 2019 the Ministry of Health, Labor and Welfare certified that five of the workers handling CWAAPs had incurred occupational injuries [4]. Consequently, to protect the health of workers who are still working in environments where CWAAPs are handled, it is extremely important to characterize the respiratory diseases caused by exposure to CWAAPs and to examine the mechanisms by which CWAAP-exposure causes these diseases.

Little is known about lung disorders caused by inhalation exposure to CWAAPs, Therefore, using a whole-body systemic inhalation system, we recently examined the effects of inhalation exposure to CWAAP in the rat, from the acute to the chronic phase [5]. The results obtained revealed: (1) inhalation exposure to CWAAP causes alveolar injury in the acute phase and continuous exposure caused regenerative changes in the alveoli; (2) during the recovery phase, some of the alveolar lesions were repaired, while others progressed to alveolitis with fibrous thickening; and (3) cell types activated by TGFβ signaling may be involved in the pathogenesis of CWAAP. However, in the above study, we adopted an irregular inhalation exposure protocol of four hours per day, once a week at high concentrations, in order to simulate exposure at the time of the industrial accident. Moreover, to ensure a sufficient number of animals per CWAAP exposure concentration, only two concentrations were used. Thus, while our previous study yielded important information, further studies are required to properly assess the effect of CWAAP exposure in experimental animals.

It is widely accepted that a reliable toxicity test should be conducted with reference to testing guidelines adopted by the Organization for Economic Co-operation and Development (OECD). The OECD guidelines for the testing of chemicals 413 (OECD TG 413) requires an exposure protocol of 6 hours per day, 5 days per week, for 90 days at more than 3 concentrations, using both male and female rodents [6].

Therefore, to obtain data that would contribute to occupational health, such as setting acceptable limits for CWAAP exposure in the workplace, in the present study we conducted a 90-day inhalation exposure test of CWAAP with reference to OECD TG 413. In addition, recovery from CWAAP inhalation exposure was investigated.

Our previous research had demonstrated that inhalation of CWAAP-A causes alveolar injury characterized by alveolar collapse with a high degree of neutrophilic infiltration in the acute phase [5]. In addition, it is generally known that macrophages are involved in pulmonary disorders caused by inorganic dust [7]. Consequently, the contribution of macrophages to the adverse effects of CWAAP-A, an organic dust, is of great interest. Therefore, in addition to the 90-day subchronic inhalation study we also conducted a second experiment to investigate whether the removal of alveolar macrophage or neutrophils would affect CWAAP-induced acute lung injury. We used clodronate liposomes for macrophage depletion [8] and polymorphonuclear leukocytes (PMN)-neutralizing antibodies for neutrophil depletion [9].

## Results

### Stability of aerosol generation and mass concentration and particle size distribution of CWAAP-A in the inhalation chamber

The mass concentrations of CWAAP-A aerosol in the inhalation chamber are shown in Table 1. Each CWAAP-A concentration was essentially equal to the target concentration over the 13-week exposure period (Fig. S1B). The mass median aerodynamic diameters (MMAD) and geometric standard deviation (GSD) of the CWAAP-A aerosol were within 0.8-0.9 μm and 2.6-2.7, respectively, and were similar for all CWAAP-exposed groups (Table 1 and Fig. S1C). Morphological observations by scanning electron microscope (SEM) confirmed that the CWAAP-A particles generated in the chamber did not appear to be immediately highly aggregated or humidified (Fig. S1D). These data indicate that the CWAAP-A aerosols were generated stably during the 13-week exposure period.

**Table 1.** The characterization of CWAAP-A generated in the inhalation chamber.

| Target concentration (mg/m<sup>3</sup>) | 0 | 0.3 | 1 | 3 | 10 |
| --- | --- | --- | --- | --- | --- |
| Temprature (°C) | 23.6 ± 0.5 | 23.5 ± 0.6 | 23.7 ± 0.7 | 23.7 ± 0.7 | 23.8 ± 0.6 |
| Humidity (%) | 46.6 ± 1.4 | 44.8 ± 1.0 | 44.5 ± 2.3 | 44.6 ± 1.0 | 45.0 ± 0.8 |
| Chamber air flow (L/min) | 411.9 ± 1.3 | 417.5 ± 2.8 | 416.1 ± 3.1 | 413.0 ± 2.9 | 416.5 ± 2.9 |
| Gravimetric mass concentration (mg/m<sup>3</sup>) | - | 0.31 ± 0.04 | 1.02 ± 0.06 | 3.05 ± 0.17 | 9.96 ± 0.52 |
| Calibrated mass concentration (mg/m<sup>3</sup>) | - | 0.31 ± 0.02 | 1.05 ± 0.05 | 3.07 ± 0.13 | 10.19 ± 0.32 |
| 1 week: MMAD µm (GSD) | - | 0.8 (2.7) | 0.9 (2.6) | 0.9 (2.6) | 0.8 (2.7) |
| 6 week: MMAD µm (GSD) | - | 0.8 (2.7) | 0.8 (2.6) | 0.8 (2.7) | 0.8 (2.7) |
| 13 week: MMAD µm (GSD) | - | 0.8 (2.6) | 0.8 (2.6) | 0.8 (2.6) | 0.8 (2.7) |

### Cytology and biochemistry of plasma, and organ weight

In all CWAAP-exposed rats, neither mortality nor respiratory clinical signs were observed throughout the study (Fig. 1A, B), and there was no change in final body weight (Fig. 1C, D). Differential white blood cell analysis showed an increase in the percentage of neutrophils and a corresponding significant decrease in the percentage of lymphocytes in the blood immediately after the 13 week exposure period in a CWAAP-A concentration dependent manner (Extended file 1). In the 10 mg/m^3^ group, these changes were still significant after the 13-week recovery period, while in the 3 mg/m^3^ exposure groups these changes were no longer significant and in the 1 mg/m^3^ and 0.3 mg/m^3^ exposure groups these changes were completely resolved. Exposure to CWAAP-A significantly decreased total cholesterol and phospholipid (PL) in the plasma immediately after the 13 week exposure period in a CWAAP-A concentration dependent manner, however, these changes were mostly resolved after the 13 week recovery period (Extended file 2). Other changes in the blood/plasma of the CWAAP-A exposed rats did not have a correlation with the exposure concentration or were very modest, suggesting low toxicological significance.

**Figure 1.**
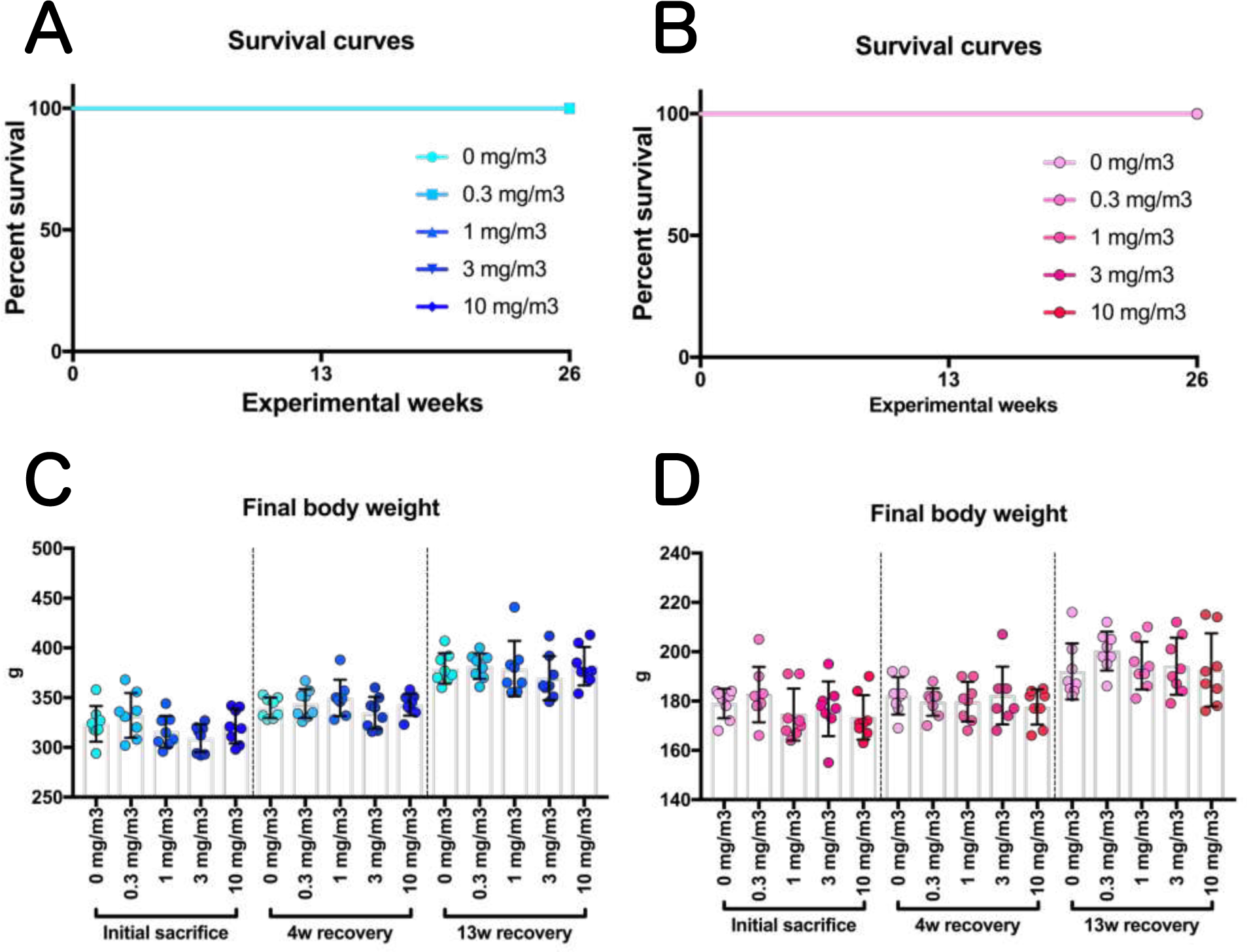
Survival curves and final body weights of rats inhaled cross-linked water-soluble acrylic acid polymer (CWAAP-A) (0.3, 1, 3 or 10 mg/m^3^, 6 hours/day, 5 days/week, 13 weeks). Final body weight of male (C) and female (D) rats were measured at each sacrifice. Abbreviation: w, week.

CWAAP-A concentration-dependent increases in lung and mediastinal lymph node weights were observed in both males and females immediately after the final exposure, and the increased weights lessened with the length of the recovery period (Figs. 2 and 3). However, even after a 13-week recovery period, significant increases in the lung weights of male rats exposed to CWAAP-A concentrations of 1 mg/m^3^ and above and female rats exposed to CWAAP-A concentrations of 3 mg/m^3^ and 10 mg/m^3^ were observed (Fig. 2), and mediastinal lymph node weights were still significantly increased in the male and female 10 mg/m^3^ groups (Fig. 3). Although CWAAP-A also caused significant increases or decreases in several other organ weights, including adrenals and ovaries, most of the changes were slight (approximately 10%) or did not correlate with exposure concentration (Extended file 3) and no gross changes were observed, suggesting low toxicological significance.

**Figure 2.**
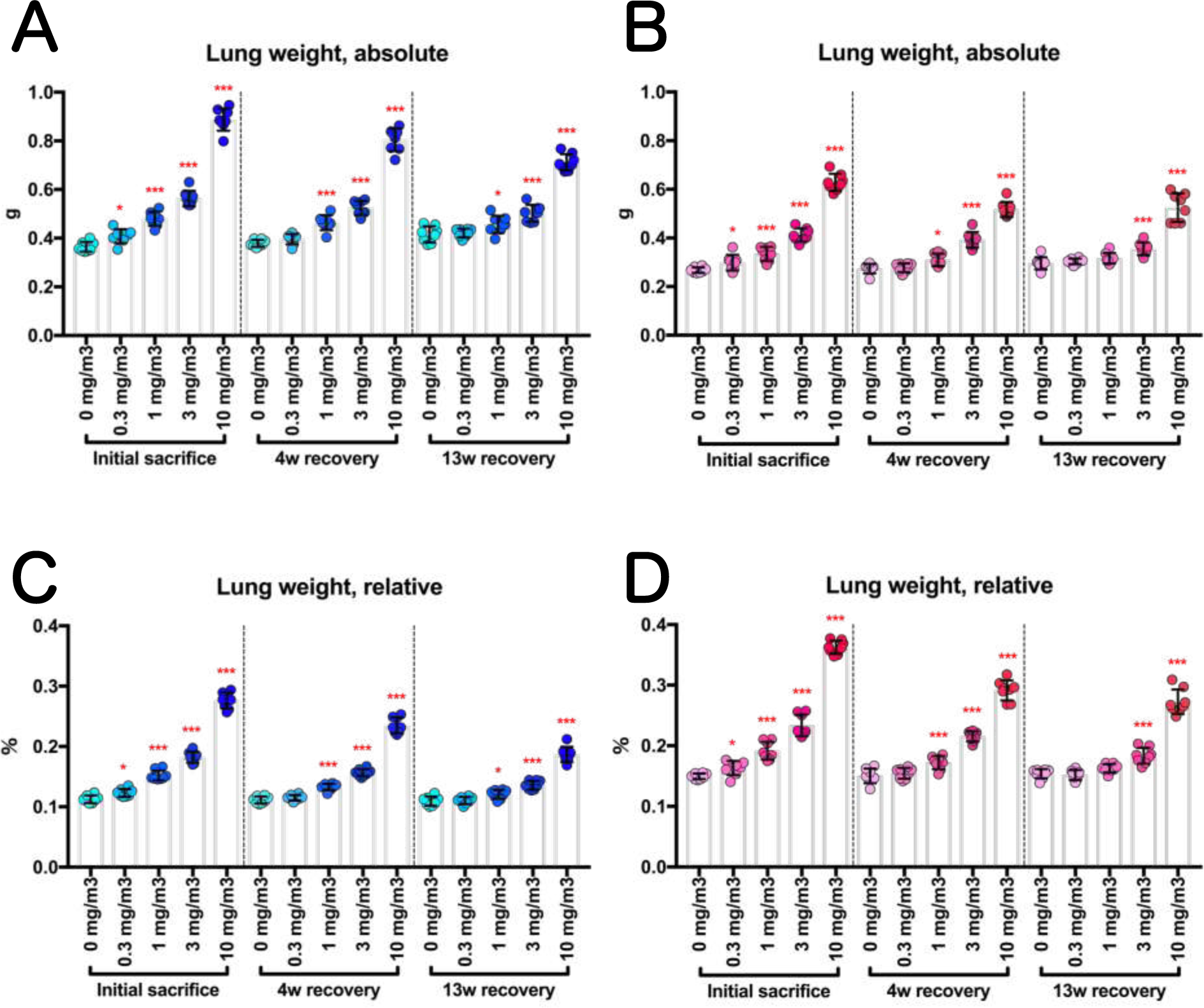
Dose-dependent increase in lung weight of rats following inhalation exposure to CWAAP-A (0.3, 1, 3 or 10 mg/m^3^, 6 hours/day, 5 days/week, 13 weeks). Absolute lung weights in male (A) and female (B) rats were measured at each sacrifice, and the relative weights of males (C) and females (D) were calculated as a percentage of body weight. William’s multiple comparison test compared with age-matched control (0 mg/m^3^) groups: \**p*<0.025 and \*\*\**p*<0.0005.

**Figure 3.**
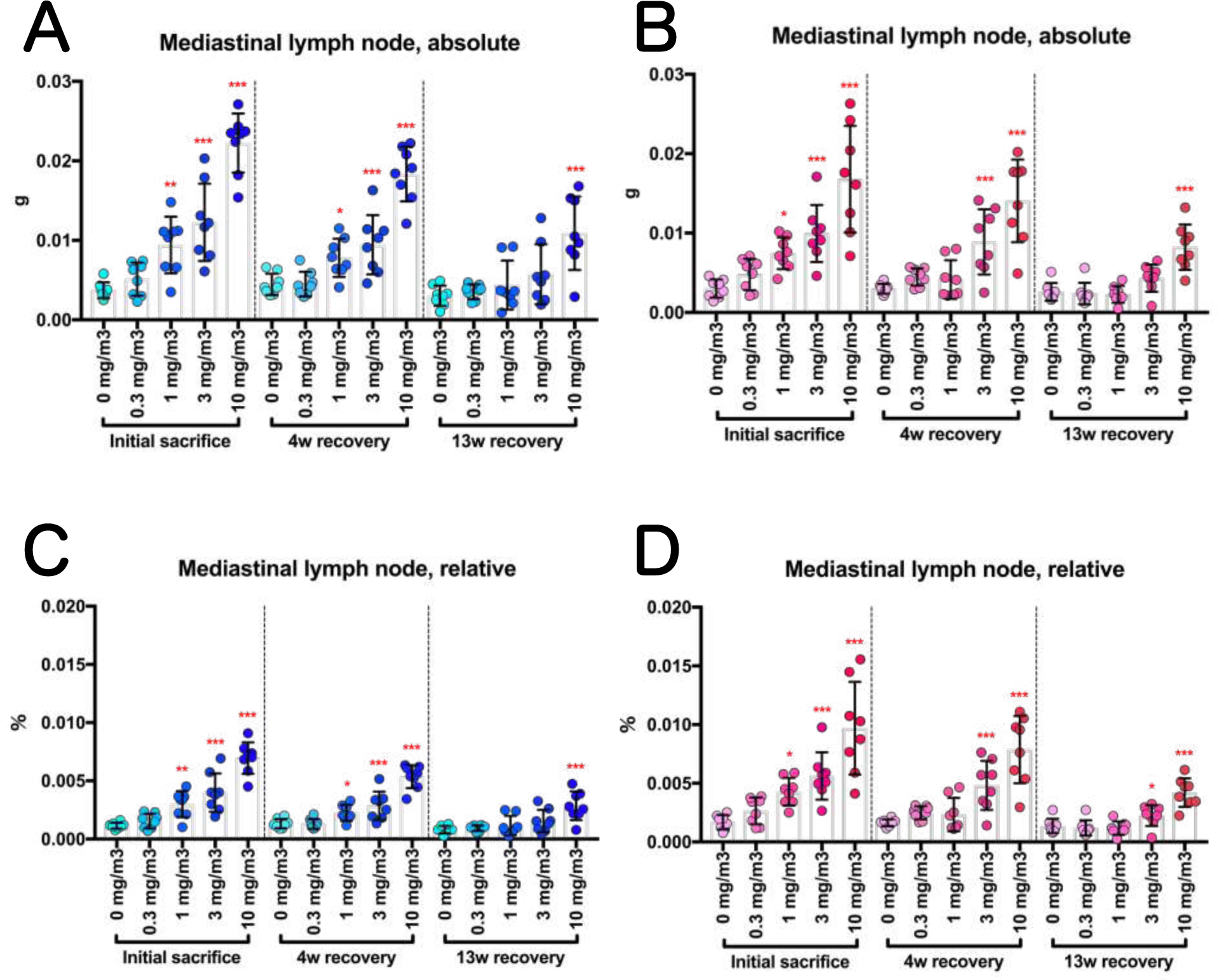
Change in mediastinal lymph node weight of rats after inhalation of CWAAP-A (0.3, 1, 3 or 10 mg/m^3^, 6 hours/day, 5 days/week, 13 weeks). Absolute weights of mediastinal lymph nodes in male (A) and female (B) rats were measured at each sacrifice. In C and D, their relative weights to body weight are shown as males (C) and females (D), respectively. William’s multiple comparison test compared with age-matched control (0 mg/m^3^) groups: \**p*<0.025, \**p*<0.005 and \*\*\**p*<0.0005.

### Macroscopic images of lung and mediastinal lymph node

Representative macroscopic images of the lungs and mediastinal lymph nodes are shown in Figs. 4 and 5 and Fig. S2. There were color changes in the lungs of the rats exposed to CWAAP-A. Compared to the pure salmon-pink color of the control group, the CWAAP-exposed lungs, especially in the 10 mg/m^3^ group, had lost their reddish color immediately after the final exposure and appeared ochre-colored, with diffuse edematous changes with marked swelling (Fig. 4). These changes had not been restored to normal after a 4-week recovery period (Fig. 4). After the 13-week recovery period, the enlargement and edema-like changes in the lungs tended to be restored to normal. In addition, enlargement of a white zone in the hilar region of the heart surface of the left lung was observed (hotspot area), which was consistent with our recent report [5].Observation of the surface (Fig. 5) and cross-section (Fig. S2) of the left lung after fixation found a large number of white spots in the 10 mg/m^3^ group immediately after exposure, and these spots were still present after the 13-week recovery period (Fig. S2). There were fewer white spots in the lungs of rats exposed to lower concentrations of CWAAP-A. Dose-dependent changes were also confirmed in mediastinal lymph nodes. Enlargement of mediastinal lymph nodes was observed after CWAAP-A exposure, and restoration to normal occurred in a recovery period dependent manner (Fig. 5). Thus, similarly to lung and mediastinal lymph node weights, the changes noted above in the lungs and mediastinal lymph nodes and the recovery to normal showed exposure concentration dependence and recovery period dependence.

**Figure 4.**
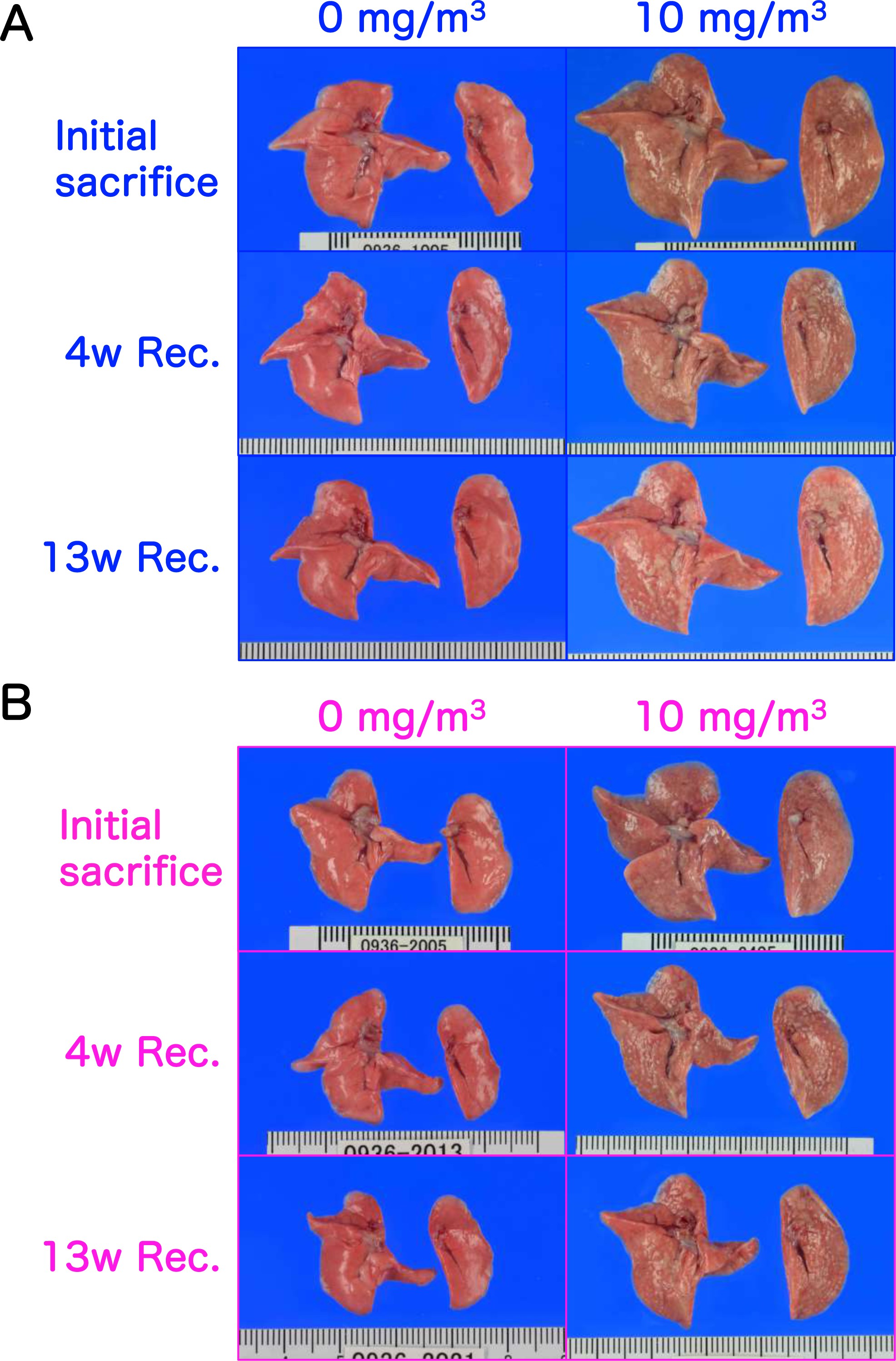
Representative macroscopic photographs of male (A) and female (B) rat lungs after 13-week inhalation exposure to CWAAP-A (10 mg/m^3^, 6 hours/day, 5 days/week). Abbreviations: Expo, exposure; Rec, recovery; and w: week.

**Figure 5.**
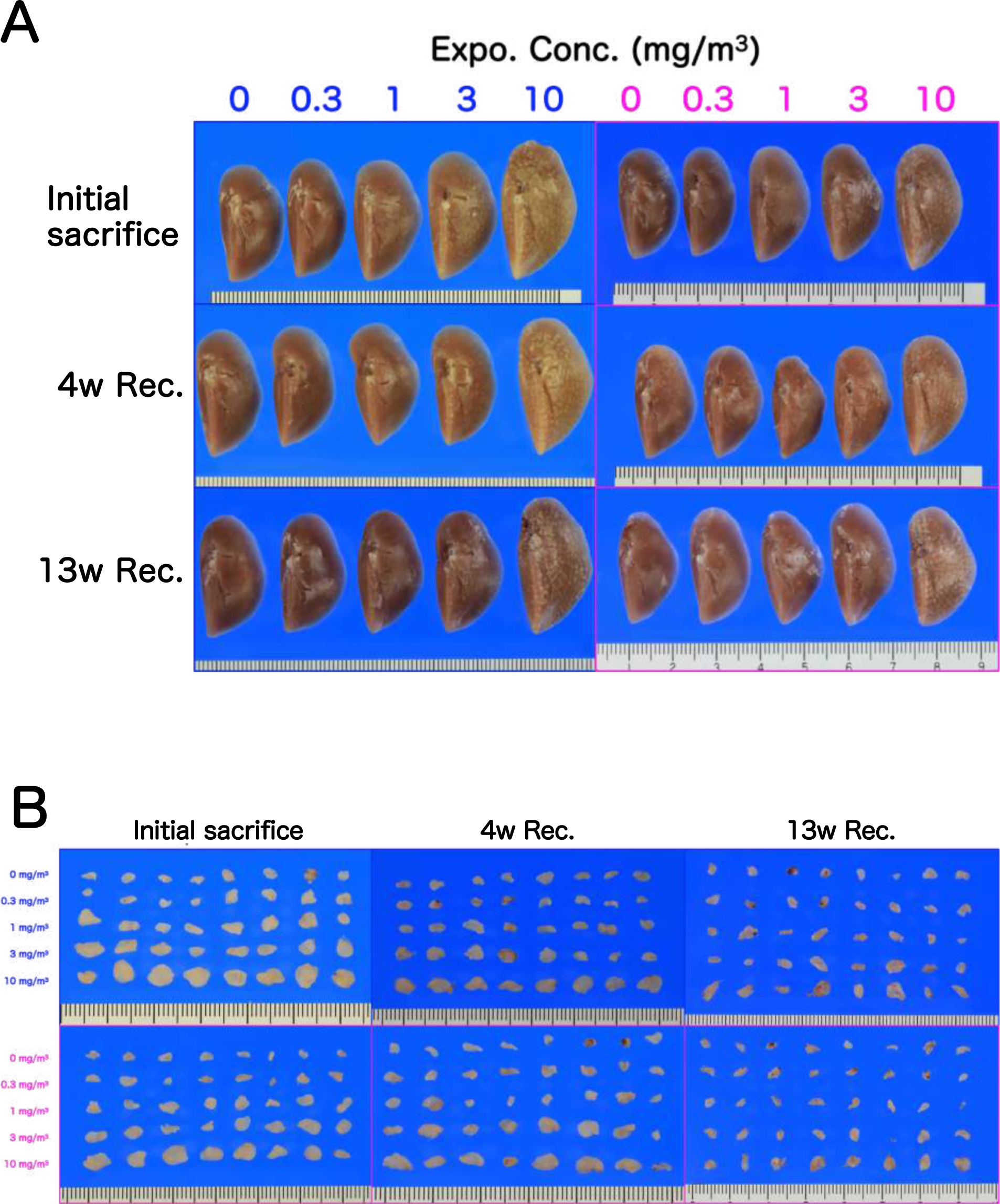
Macroscopic photographs of lungs (A) and mediastinal lymph nodes (B) in CWAAP-A-exposed rats after fixation. (A) Photographs of males on the left and females on the right. (B) Photographs of males on the top and females on the bottom.

### Histopathological examination for lung and mediastinal lymph node

Histopathological data for the lung and mediastinal lymph nodes immediately after the last exposure are shown in Table 2. CWAAP-A exposure induced various inflammation/injury and associated regeneration-related findings in the pulmonary alveolar region. Granulomatous changes, commonly observed with inhalation exposure to dust, were observed in all rats exposed to CWAAP-A at concentrations of 1 mg/m^3^ and above, and lesion intensity was concentration-dependent (Table 2). In addition, significant accumulation of lipoproteinous material, which may represent a mild form of pulmonary alveolar proteinosis, was observed in male rats exposed to 10 mg/m^3^ and female rats exposed to 3 mg/m^3^ and 10 mg/m^3^ CWAAP-A (Table 2). Accumulation of lipoproteinous material was observed principally around the hilar region of the heart surface, a hotspot (Fig. 6). Importantly, exposure to 1 mg/m^3^ or higher concentrations of CWAAP-A caused “multifocal lesions” in the lungs, consisting mainly of enlarged/proliferating alveolar epithelium and infiltration of neutrophils and macrophages and inflammation of the air spaces. In the mediastinal lymph nodes, significant lymphoid hyperplasia was seen in males exposed to 3 and 10 mg/m^3^ CWAAP-A and in females exposed to 10 mg/m^3^ (Table 2). Taken together, all the lesions observed were caused by exposure to 1 mg/m^3^ or more CWAAP-A, and no pathological toxicity was observed in male or female rats exposed to 0.3 mg/m^3^ CWAAP-A.

**Figure 6.**
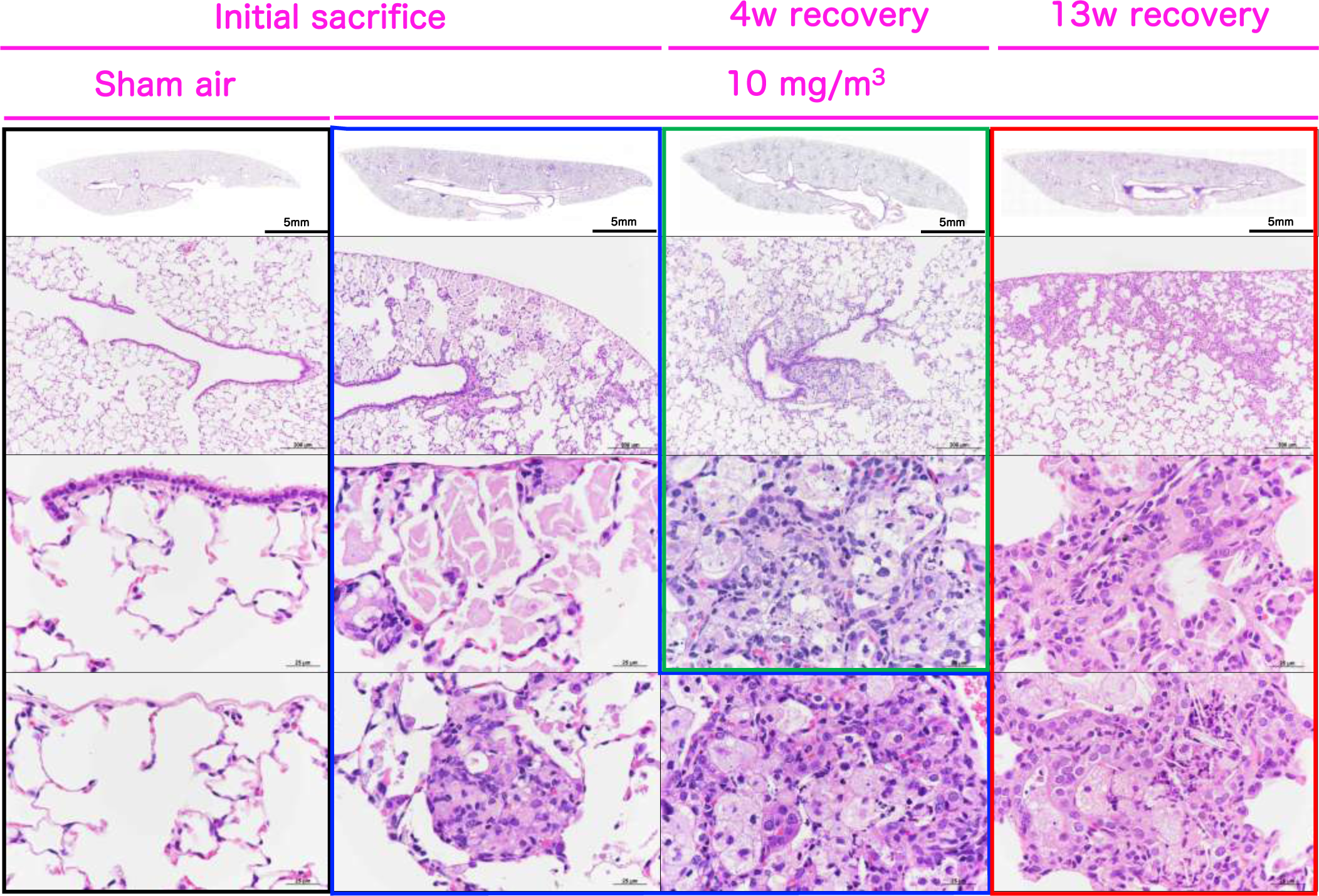
Representative microscopic photographs of female rat lungs after inhalation exposure to CWAAP-A (10 mg/m^3^). The control (0 mg/m^3^) and CWAAP-A-exposed lungs were stained with hematoxylin and eosin (HE), and their typical lesions are shown for each recovery period.

**Table 2.** Incidence and integrity of the histopathological findings of the lung and mediastinal lymph nodes immediately after 90-day inhalation exposure to CWAAP-A.

|  |  | Male |  |  |  |  | Female |  |  |  |  |
| --- | --- | --- | --- | --- | --- | --- | --- | --- | --- | --- | --- |
| Exposure concentration (mg/m<sup>3</sup>) |  | 0 | 0.3 | 1 | 3 | 10 | 0 | 0.3 | 1 | 3 | 10 |
| No. of Animals Examined |  | 8 | 8 | 8 | 8 | 8 | 8 | 8 | 8 | 8 | 8 |
| Histopathological findings |  |  |  |  |  |  |  |  |  |  |  |
| Mediastinal lymph node |  |  |  |  |  |  |  |  |  |  |  |
|  | Lymphoid hyperplasia | 0 | 0 | 1<br><1> | 4 *<br><1> | 6 **<br><1.3> | 0 | 1<br><1> | 1<br><1> | 1<br><1> | 4 *<br><1.3> |
| Lung |  |  |  |  |  |  |  |  |  |  |  |
|  | Deposition of test compound (AB-pos granules) | 0 | 8 ***<br><1> | 8 ***<br><1> | 8 ***<br><1> | 8 ***<br><2> | 0 | 8 ***<br><1> | 8 ***<br><1> | 8 ***<br><2> | 8 ***<br><2> |
|  | Granulomatous change, alveolar | 0 | 0 | 8 ***<br><1> | 8 ***<br><1.1> | 8 ***<br><1.9> | 0 | 0 | 8 ***<br><1> | 8 ***<br><1> | 8 ***<br><1.6> |
|  | Multifocal lesions, alveolar |  |  |  |  |  |  |  |  |  |  |
|  | Hypertrophy/Proliferation of alveolar epithelium | 0 | 0 | 3<br><1> | 8 ***<br><1.1> | 8 ***<br><2> | 0 | 0 | 6 **<br><1> | 8 ***<br><1> | 8 ***<br><2> |
|  | Inflammation, air space | 0 | 0 | 6 **<br><1> | 8 ***<br><1.1> | 8 ***<br><2> | 0 | 0 | 6 **<br><1> | 8 ***<br><1> | 8 ***<br><2> |
|  | Cholesterol creft, air space | 0 | 0 | 0 | 0 | 0 | 0 | 0 | 0 | 0 | 0 |
|  | Alveolitis | 0 | 0 | 0 | 0 | 0 | 0 | 0 | 0 | 0 | 0 |
|  | Fibrous thickening, interstitial | 0 | 0 | 0 | 0 | 0 | 0 | 0 | 0 | 0 | 0 |
|  | Accumulation of Lipoproteinous material, air space | 0 | 0 | 0 | 2<br><1> | 8 ***<br><1.9> | 0 | 0 | 0 | 6 **<br><1> | 8 ***<br><1.9> |

Note: Values indicate number of animals bearing lesions. The value in angle bracket indicate the average of severity grade index of the lesion. The average of severity grade is calculated with a following equation. $\Sigma(\text{grade} \times \text{number of animals with grade}) / \text{number of affected animals}$ Grade: 1, slight; 2, moderate; 3, marked; 4, severe. Significant difference: \*, p<0.05; \*\*, p<0.01; \*\*\*, p<0.001 by Chi square test compared with each cont.

The 13-week recovery period resulted in complete resolution of tissue damage caused by exposure to 1 mg/m^3^ CWAAP-A for 13 weeks in both male and female rats (Tables 2 and 4). The only abnormality in the lungs of these animals was granulation tissue encapsulating CWAAP-A. In contrast, rats exposed to 3 mg/m^3^ and 10 mg/m^3^ developed cholesterol cleft, alveolitis which is a pathological finding of interstitial pneumonia, and fibrous thickening of the interstitium. These lesions were not present in any of the rats immediately after the end of the inhalation exposure period (Table 2), but had begun to develop at the 4 week recovery period (Table 3), and were all significantly increased (except for cholesterol cleft in the 3 mg/m^3^ male group) at the 13 week recovery period (Table 4). These findings indicate that lung disease developed in the rats exposed to 3 mg/m^3^ and 10 mg/m^3^ CWAAP-A.

**Table 3.** Incidence and integrity of the histopathological findings of the lung and mediastinal lymph nodes 4 weeks after 90-day inhalation exposure to CWAAP-A.

|  |  | Male |  |  |  |  | Female |  |  |  |  |
| --- | --- | --- | --- | --- | --- | --- | --- | --- | --- | --- | --- |
| Exposure concentration (mg/m<sup>3</sup>) |  | 0 | 0.3 | 1 | 3 | 10 | 0 | 0.3 | 1 | 3 | 10 |
| No. of Animals Examined |  | 8 | 8 | 8 | 8 | 8 | 8 | 8 | 8 | 8 | 8 |
| Histopathological findings |  |  |  |  |  |  |  |  |  |  |  |
| Mediastinal lymph node |  |  |  |  |  |  |  |  |  |  |  |
|  | Lymphoid hyperplasia | 0 | 0 | 1 | 1 | 2 | 0 | 0 | 0 | 0 | 1 |
|  |  |  |  | <1> | <1> | <1> |  |  |  |  | <1> |
| Lung |  |  |  |  |  |  |  |  |  |  |  |
|  | Deposition of test compound (AB-pos granules) | 0 | 8 *** | 8 *** | 8 *** | 8 *** | 0 | 8 *** | 8 *** | 8 *** | 8 *** |
|  |  |  | <1> | <1> | <1> | <2> |  | <1> | <1> | <1> | <2> |
|  | Granulomatous change, alveolar | 0 | 0 | 8 *** | 8 *** | 8 *** | 0 | 0 | 8 *** | 8 *** | 8 *** |
|  |  |  |  | <1> | <1> | <1.9> |  |  | <1> | <1> | <2> |
|  | Multifocal lesions, alveolar |  |  |  |  |  |  |  |  |  |  |
|  | Hypertrophy/Proliferation of alveolar epithelium | 0 | 0 | 1 | 8 *** | 8 *** | 0 | 0 | 2 | 8 *** | 8 *** |
|  |  |  |  | <1> | <1> | <2> |  |  | <1> | <1> | <2> |
|  | Inflammation, air space | 0 | 0 | 1 | 8 *** | 8 *** | 0 | 0 | 2 | 8 *** | 8 *** |
|  |  |  |  | <1> | <1> | <2> |  |  | <1> | <1> | <2> |
|  | Cholesterol creft, air space | 0 | 0 | 0 | 0 | 8 *** | 0 | 0 | 0 | 2 | 8 *** |
|  |  |  |  |  |  | <1> |  |  |  | <1> | <1> |
|  | Alveolitis | 0 | 0 | 0 | 0 | 3 | 0 | 0 | 0 | 0 | 3 |
|  |  |  |  |  |  | <1> |  |  |  |  | <1> |
|  | Fibrous thickening, interstitial | 0 | 0 | 0 | 0 | 3 | 0 | 0 | 0 | 0 | 3 |
|  |  |  |  |  |  | <1> |  |  |  |  | <1.3> |
|  | Accumulation of Lipoproteinous material, air space | 0 | 0 | 0 | 0 | 6 ** | 0 | 0 | 0 | 1 | 8 *** |
|  |  |  |  |  |  | <1> |  |  |  | <1> | <1.1> |

Note: Values indicate number of animals bearing lesions. The value in angle bracket indicate the average of severity grade index of the lesion. The average of severity grade is calculated with a following equation. $\Sigma(\text{grade} \times \text{number of animals with grade}) / \text{number of affected animals}$ Grade: 1, slight; 2, moderate; 3, marked; 4, severe. Significant difference: \*, p<0.05; \*\*, p<0.01; \*\*\*, p<0.001 by Chi square test compared with each cont.

**Table 4.** Incidence and integrity of the histopathological findings of the lung and mediastinal lymph nodes 13 weeks after 90-day inhalation exposure to CWAAP-A.

|  |  | Male |  |  |  |  | Female |  |  |  |  |
| --- | --- | --- | --- | --- | --- | --- | --- | --- | --- | --- | --- |
| Exposure concentration (mg/m<sup>3</sup>) |  | 0 | 0.3 | 1 | 3 | 10 | 0 | 0.3 | 1 | 3 | 10 |
| No. of Animals Examined |  | 8 | 8 | 8 | 8 | 8 | 8 | 8 | 8 | 8 | 8 |
| Histopathological findings |  |  |  |  |  |  |  |  |  |  |  |
| Mediastinal lymph node |  |  |  |  |  |  |  |  |  |  |  |
|  | Lymphoid hyperplasia | 0 | 0 | 0 | 0 | 0 | 0 | 0 | 0 | 0 | 0 |
| Lung |  |  |  |  |  |  |  |  |  |  |  |
|  | Deposition of test compound (AB-pos granules) | 0 | 3 | 8 *** | 8 *** | 8 *** | 0 | 3 | 8 *** | 8 *** | 8 *** |
|  |  |  | <1> | <1> | <1> | <2> |  | <1> | <1> | <1> | <2> |
|  | Granulomatous change, alveolar | 0 | 0 | 8 *** | 8 *** | 8 *** | 0 | 0 | 8 *** | 8 *** | 8 *** |
|  |  |  |  | <1> | <1> | <1> |  |  | <1> | <1> | <1.3> |
|  | Multifocal lesion, alveolar |  |  |  |  |  |  |  |  |  |  |
|  | Hypertrophy/Proliferation of alveolar epithelium | 0 | 0 | 0 | 6 ** | 8 *** | 0 | 0 | 0 | 7 *** | 8 *** |
|  |  |  |  |  | <1> | <1.8> |  |  |  | <1.1> | <1.8> |
|  | Inflammation, air space | 0 | 0 | 0 | 8 *** | 8 *** | 0 | 0 | 0 | 8 *** | 8 *** |
|  |  |  |  |  | <1> | <1> |  |  |  | <1.1> | <1> |
|  | Cholesterol creft, air space | 0 | 0 | 0 | 1 | 8 *** | 0 | 0 | 0 | 4 * | 8 *** |
|  |  |  |  |  | <1> | <1> |  |  |  | <1> | <1.1> |
|  | Alveolitis | 0 | 0 | 0 | 6 ** | 8 *** | 0 | 0 | 0 | 6 ** | 8 *** |
|  |  |  |  |  | <1> | <2> |  |  |  | <1.2> | <1.8> |
|  | Fibrous thickening, interstitial | 0 | 0 | 0 | 6 ** | 8 *** | 0 | 0 | 0 | 6 ** | 8 *** |
|  |  |  |  |  | <1> | <1> |  |  |  | <1> | <1.5> |
|  | Accumulation of Lipoproteinous material, air space | 0 | 0 | 0 | 0 | 8 *** | 0 | 0 | 0 | 0 | 7 *** |
|  |  |  |  |  |  | <1> |  |  |  |  | <1.4> |

Note: Values indicate number of animals bearing lesions. The value in angle bracket indicate the average of severity grade index of the lesion. The average of severity grade is calculated with a following equation. I/(grade X number of animals with grade)/number of affected animals. Grade: 1, slight; 2, moderate; 3, marked; 4, severe. Significant difference: \*, p<0.05; \*\*, p<0.01; \*\*\*, p<0.001 by Chi square test compared with each cont.

### Involvement of the lymphatic vessels in CWAAP-induced pulmonary disorders

The main routes for removing foreign bodies aspirated into the lung are by retrograde movement of the mucus comprising the bronchial mucosa driven by the ciliated cells lining the lung passages, i.e., the mucociliary escalator, and drainage via lymphatic vessels [10–12]. We investigated the relationship between lymphatic vessels and localization of multifocal lesions in the rat lung. First, we visualized lymphatic vessels by green fluorescent protein (GFP) staining in the lungs of Prospero homeobox protein-1(Prox1)-EGFP transgenic rats [13] (Fig. S3). We then examined the co-localization of EGFP with commercially available lymphatic markers, including vascular endothelial growth factor receptor 3 (VEGFR3), lymphatic vessel endothelial hyaluronan receptor 1 (Lyve-1) and podoplanin (RT1-40) (Fig. S4). VEGFR3 showed the best co-localization with GFP (Fig. S4). We then examined the localization of lymphatic vessels in lungs exposed to CWAAP-A, we stained lung sections with α-smooth muscle actin (αSMA), a marker for arterial and venous smooth muscle, and VEGFR3 (Fig. 7). Both immediately after the end of the exposure period and after the 13 week recovery period, lungs exposed to 10 mg/m^3^ CWAAP-A had dilated collecting lymphatic vessels both in the hilar region and bronchovascular bundle interstitium, and an increase in the number of capillary lymphatic vessels (Fig. 7B and 7C). These results indicate that the CWAAP-A exposure affected the lymphatic vasculature.

**Figure 7.**
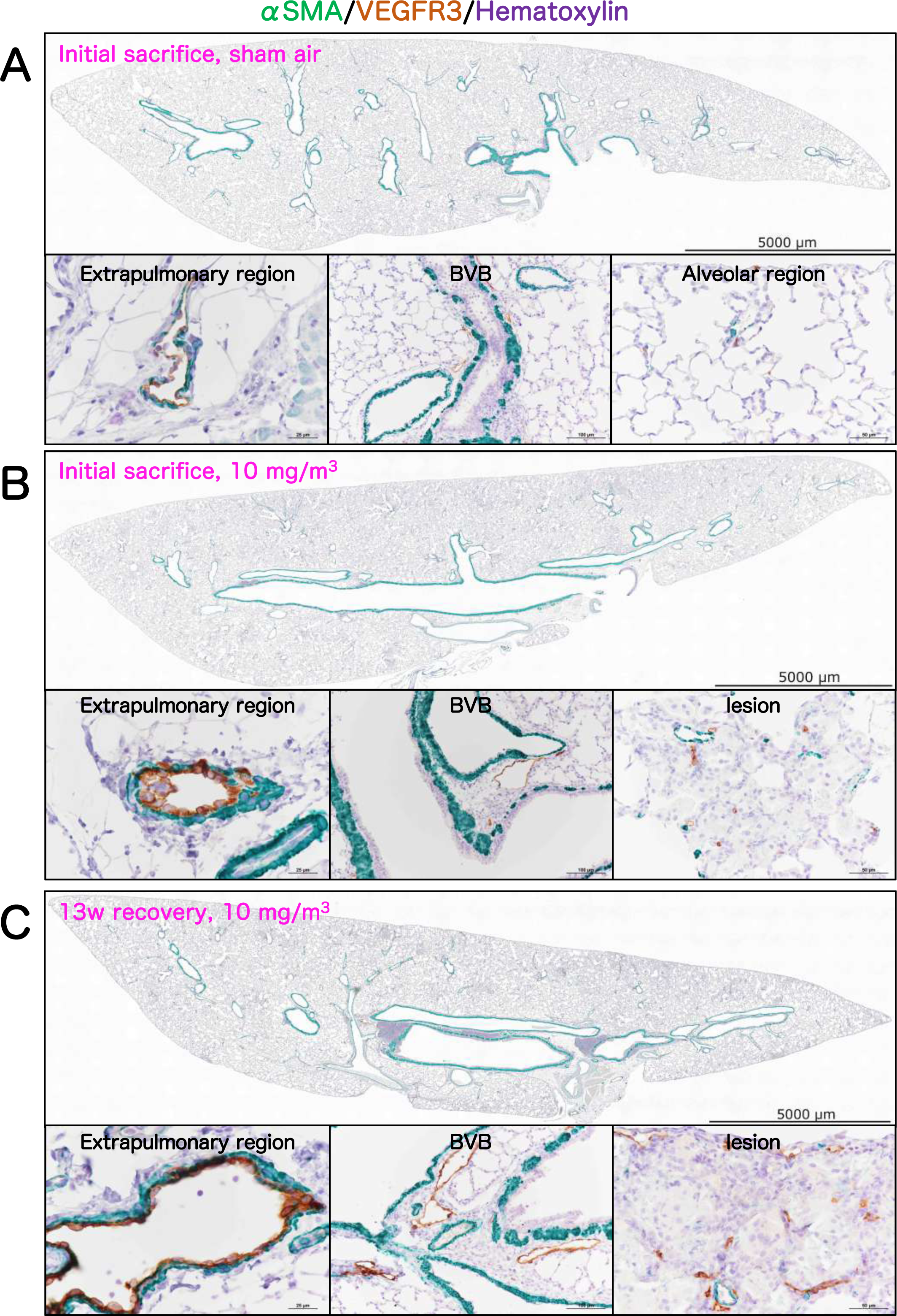
Double staining of rat lung for α-smooth muscle actin (α SMA; green), and vascular endothelial growth factor receptor 3 (VEGFR3; a lymphatic marker, brown in apical membrane), with counterstaining of hematoxylin. Representative loupe photograph and their magnified images of extrapulmonary region, bronchovascular bundle (BVB), alveolar region and CWAAP-A-induced lesion are shown for each control (A), immediately after 13-week exposure (B) and during the 13 week recovery period (C).

### Histopathological findings in other organs

Histopathological findings in the nasal cavity, trachea, liver, stomach, pancreas, kidney, prostate, pituitary gland, thyroid, mammary gland, uterus, brain, eye, harder gland, and bone marrow are shown in Extended file 4. Inhalation exposure to CWAAP-A did not affect these organs, indicating that the effects of CWAAP-A exposure are limited to the alveolar region of the lungs and the mediastinal lymph nodes.

### CWAAP-A deposition in the lung

As reported previously, our modified alcian blue staining method stains CWAAPs [5]. The results of alcian blue staining of the lung sections from a rat sacrificed immediately after the end of the exposure period are shown in Fig. 8. In accordance with our previous report, blue-stained structures were specifically observed in lesions in the alveolar region and normal periapical tissues only in the CWAAP-A exposed rats. The incidences and grading of CWAAP-A deposition are shown in Tables 2-4. Immediately after the exposure period and after the 4 and 13 week recovery periods, blue particles were observed in the lungs of all male and female exposed groups. Notably, in the 0.3 mg/m^3^ groups, after the 13 week recovery period, blue particles were found in only 3 of the 8 animals examined, indicating some clearance of CWAAP-A from the lungs of the rats exposed to 0.3 mg/m^3^ CWAAP-A.

**Figure 8.**
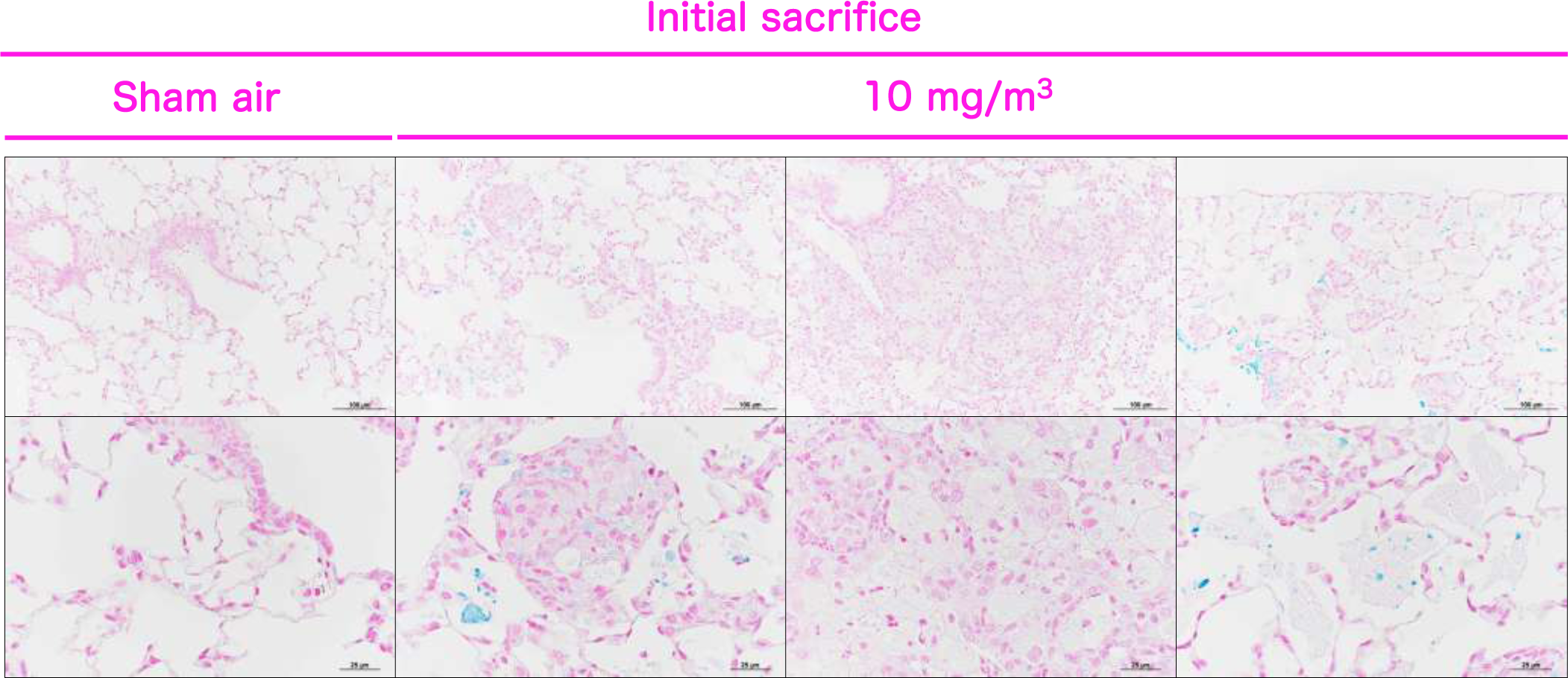
Representative images of alcian blue staining in the normal lung and the lesions of the rat lungs after inhalation exposure to CWAAP-A (10 mg/m^3^).

### Measurement of cytological and biochemical markers, including fibrotic changes

In the sham air group of both sexes, normal macrophages with fine vacuoles were observed in the bronchoalveolar lavage fluid (BALF) (Fig. 9). However, in the BALF of the 10 mg/m^3^ group a large number of neutrophils and enlarged macrophages phagocytosing CWAAP-A were observed immediately after the end of the exposure period and after the 4 and 13 week recovery periods (Fig. 9): CWAAP-A deposits can also be seen in Figure 9. Cell analysis of the BALF found that CWAAP-A exposure increased total cell number in a concentration-dependent manner, and immediately after the end of the exposure period this increase was significant in rats exposed to 1 mg/m^3^ and higher concentrations of CWAAP-A (Fig. 10A and B). Total cell numbers in the BALF decreased after the 4 week and 13 week recovery periods, and only the 10 mg/m^3^ CWAAP-A exposed groups had significantly increased cell numbers in the BALF after the 13 week recovery period. Neutrophil, alveolar macrophage, and lymphocyte counts in the BALF immediately after the end of the exposure period and after the 4 week and 13 week recovery periods are shown in Fig. 10C-H, neutrophils were the major contributor to the effect that CWAAP-A exposure had on total cell number in the BALF. Notably, after the 13-week recovery period, the alveolar macrophage count in males had returned to normal in all exposed groups, while in females the alveolar macrophage count was statistically higher in the 10 mg/m^3^ group compared to the controls, suggesting a difference between males and females.

**Figure 9.**
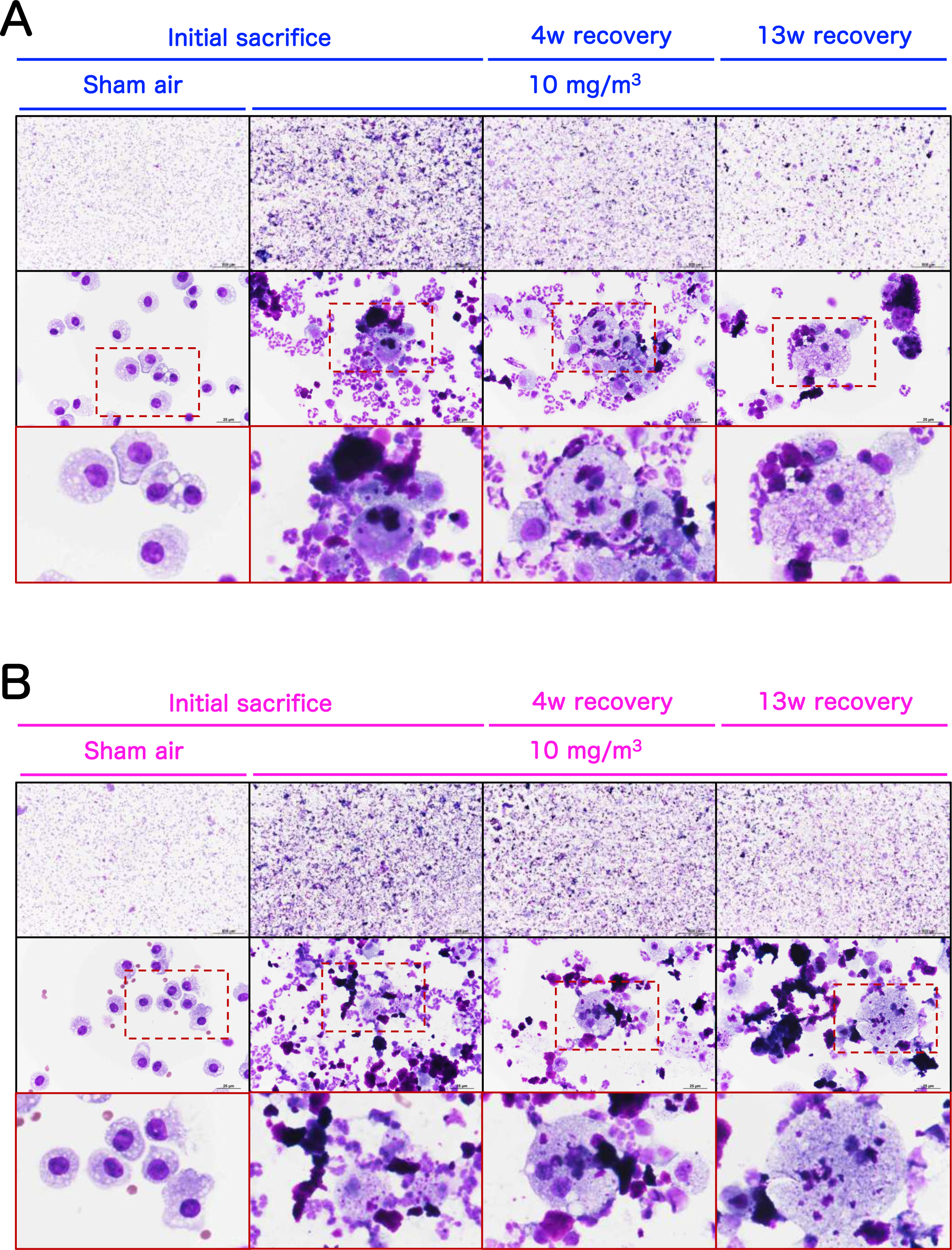
Representative images of the bronchoalveolar lavage fluid (BALF) cytospin cytology. The BALF samples obtained from males (A) and females (B) exposed to 10 mg/m^3^ CWAAP-A were stained with May-Grunwald-Giemsa.

**Figure 10.**
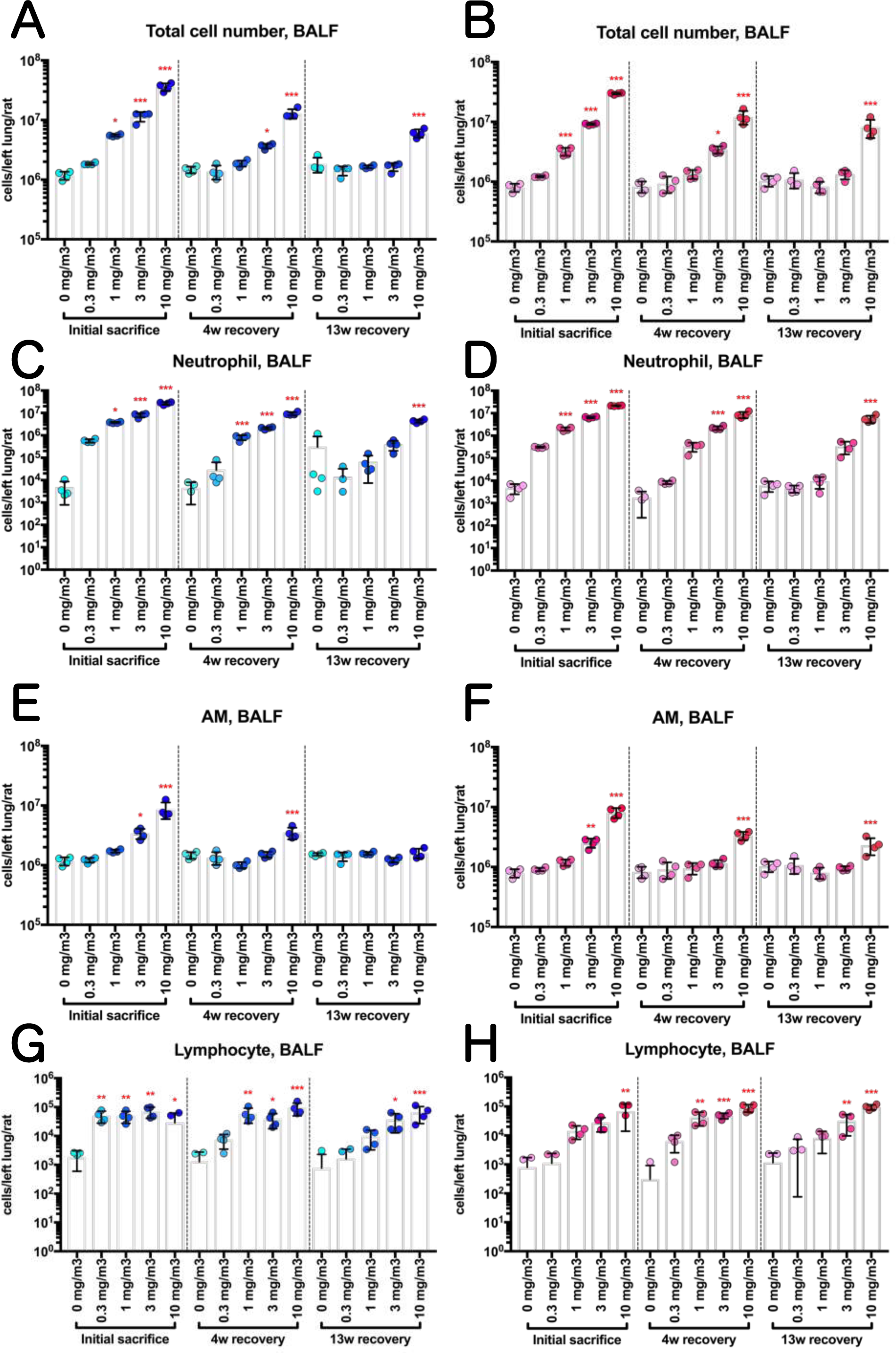
Effect of inhalation exposure to CWAAP-A on cell number in BALF. The number of total cells (A, B), neutrophils (C, D), alveolar macrophages (AM) (E, F) and lymphocytes (G, H) were counted using an automated hematology analyzer, and are shown by sex (males: A, C, E and G; females: B, D, F and H). Statistical significance was analyzed using William’s multiple comparison test compared with age-matched control (0 mg/m^3^) groups: \**p*<0.025, \*\**p*<0.005 and \*\*\**p*<0.0005.

CWAAP-A exposure increased lung lesion markers in the BALF and plasma. In both male and female groups, the activities of lactate dehydrogenase (LDH), alkaline phosphatase (ALP), and γ-glutamyl transpeptidase (γ-GTP) and the levels of PL in the BALF were all increased in an exposure concentration-dependent manner (Fig. 11A-H). ALP activity, γ-GTP activity, and PL levels had returned to normal after the 13 week recovery period in the male 0.3, 1, and 3 mg/m^3^ exposure groups, and LDH activity had returned to normal in the 0.3 and 1 mg/m^3^ groups. In female rats, LDH activity, ALP activity, and PL levels had returned to normal after the 13 week recovery period in the 0.3, 1, and 3 mg/m^3^ exposure groups, and γ-GTP activity had returned to normal in the 0.3 and 1 mg/m^3^ groups. In both males and females exposed to 10 mg/m^3^ CWAAP-A, the activities of LDH, ALP, and γ-GTP and the PL levels remained significantly elevated after the 13-week recovery period. In agreement with these lung lesion markers, the plasma and BALF levels of surfactant protein D (SP-D), a marker for interstitial pneumonia, were increased by CWAAP exposure in an exposure concentration-dependent manner (Fig. 12A-D). After the 13-week recovery period, SP-D levels in the BALF of both males and females had returned to normal in the 0.3, 1, and 3 mg/m^3^ exposure groups, and SP-D levels in the plasma had returned to normal in the male 0.3, 1, and 3 mg/m^3^ exposure groups and in the female 0.3 and 1 mg/m^3^ exposure groups. In both males and females exposed to 10 mg/m^3^ CWAAP-A, the SP-D levels remained significantly elevated after the 13 week recovery period in both the BALF and plasma.

**Figure 11.**
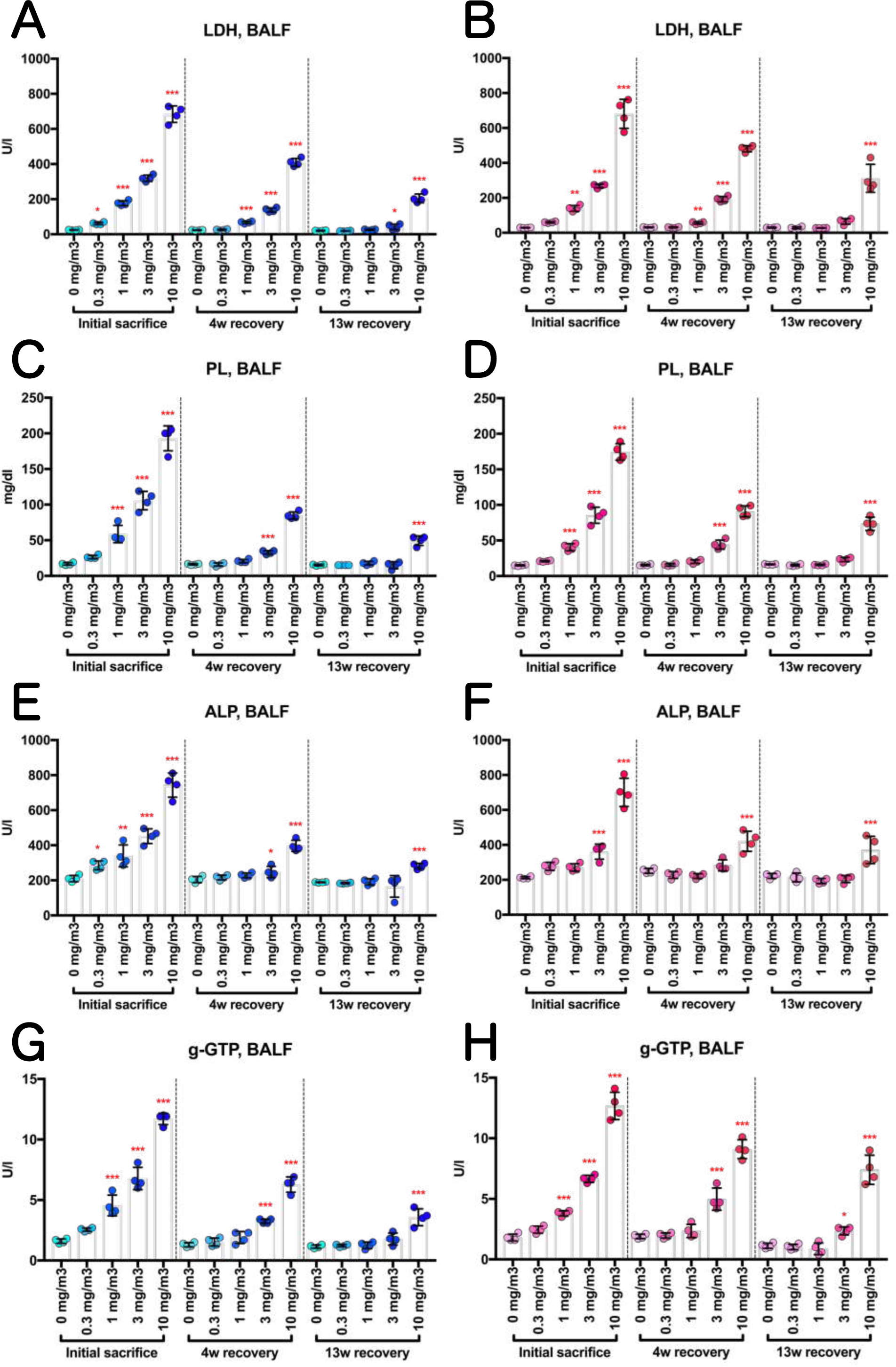
Dose-dependent induction of biochemical markers in BALF obtained from the lungs of phospholipid (PL) concentration (C, D), alkaline phosphatase (ALP) activity (E, F), γ-glutamyl transpeptidase (γ-GTP) activity (G, H) were measured using an automatic analyzer, and are shown by sex (males: A, C, E and G; females: B, D, F and H). Statistical significance was analyzed using William’s multiple comparison test compared with age-matched control (0 mg/m^3^) groups: \**p*<0.025, \*\**p*<0.005 and \*\*\**p*<0.0005.

**Figure 12.**
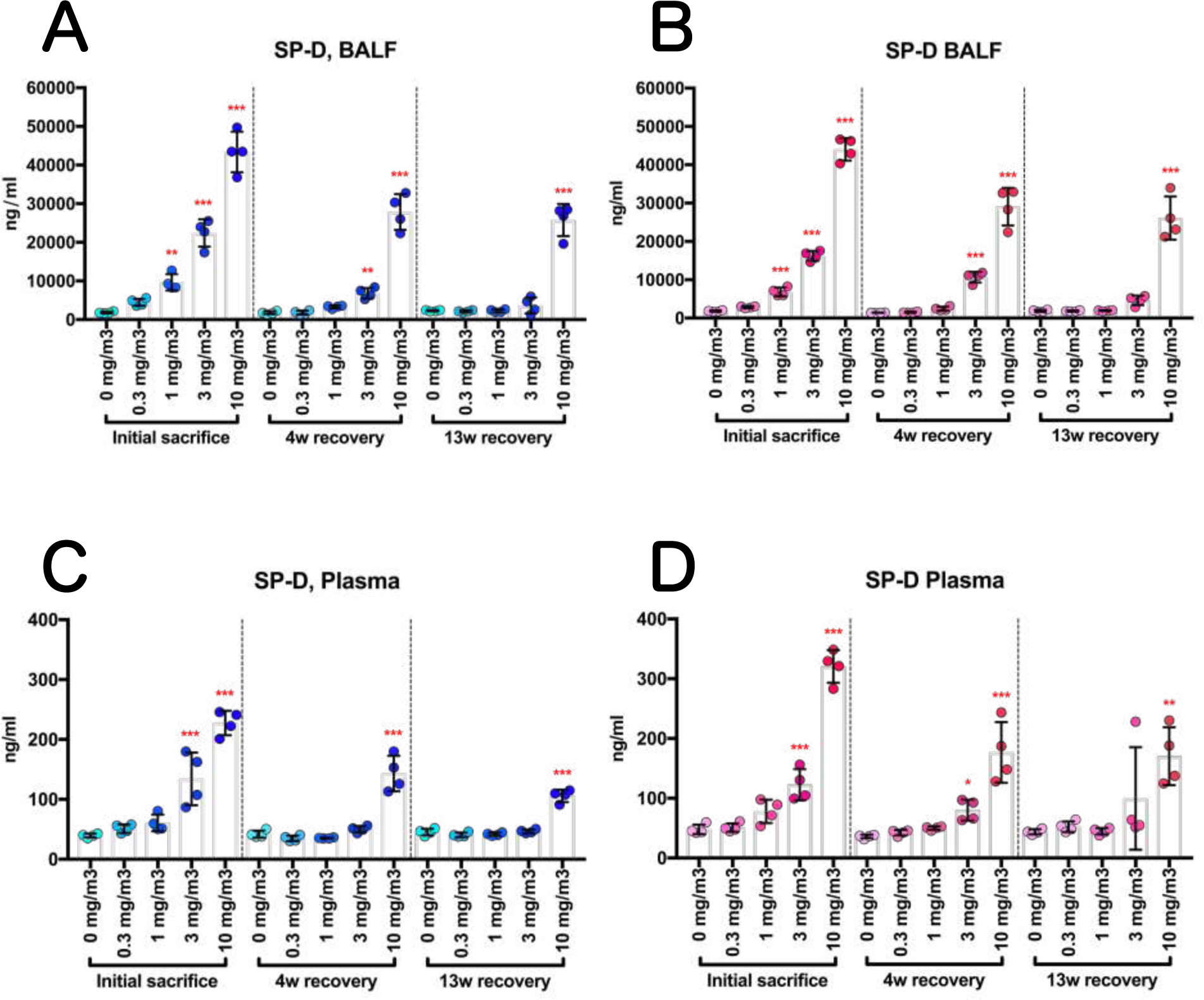
The increase in the level of surfactant protein-D (SP-D) of rats by inhalation exposure to CWAAP-A for 13 weeks. SP-D level in BALF (A, B) and plasma (C, D) were measured by an enzyme immunoassay, and are shown by sex (males: A and C; females: B and D). Statistical significance was analyzed using William’s multiple comparison test compared with age-matched control (0 mg/m^3^) groups: \**p*<0.025, \*\**p*<0.005 and \*\*\**p*<0.0005.

To quantify the fibrotic changes in the lungs, we measured the levels of transforming growth factor TGFβ1 and TGFβ2, the major signaling molecules for fibrosis [14, 15], in the BALF and the levels of hydroxyproline, a major component of collagen fibers, in the lung. Similarly to the markers of lung lesions discussed above, in both males and females the levels of TGFβ1 and TGFβ2 in the BALF were increased in an exposure concentration-dependent manner (Fig. 13A-D), and the rats recovered from these increases over time until TGFβ1 was significantly increased only in the 10 mg/m^3^ CWAAP-A exposed groups after the 13 week recovery period and TGFβ2 was significantly increased only in the 10 mg/m3 CWAAP-A exposed groups after the 4 week and 13 week recovery periods.

**Figure 13.**
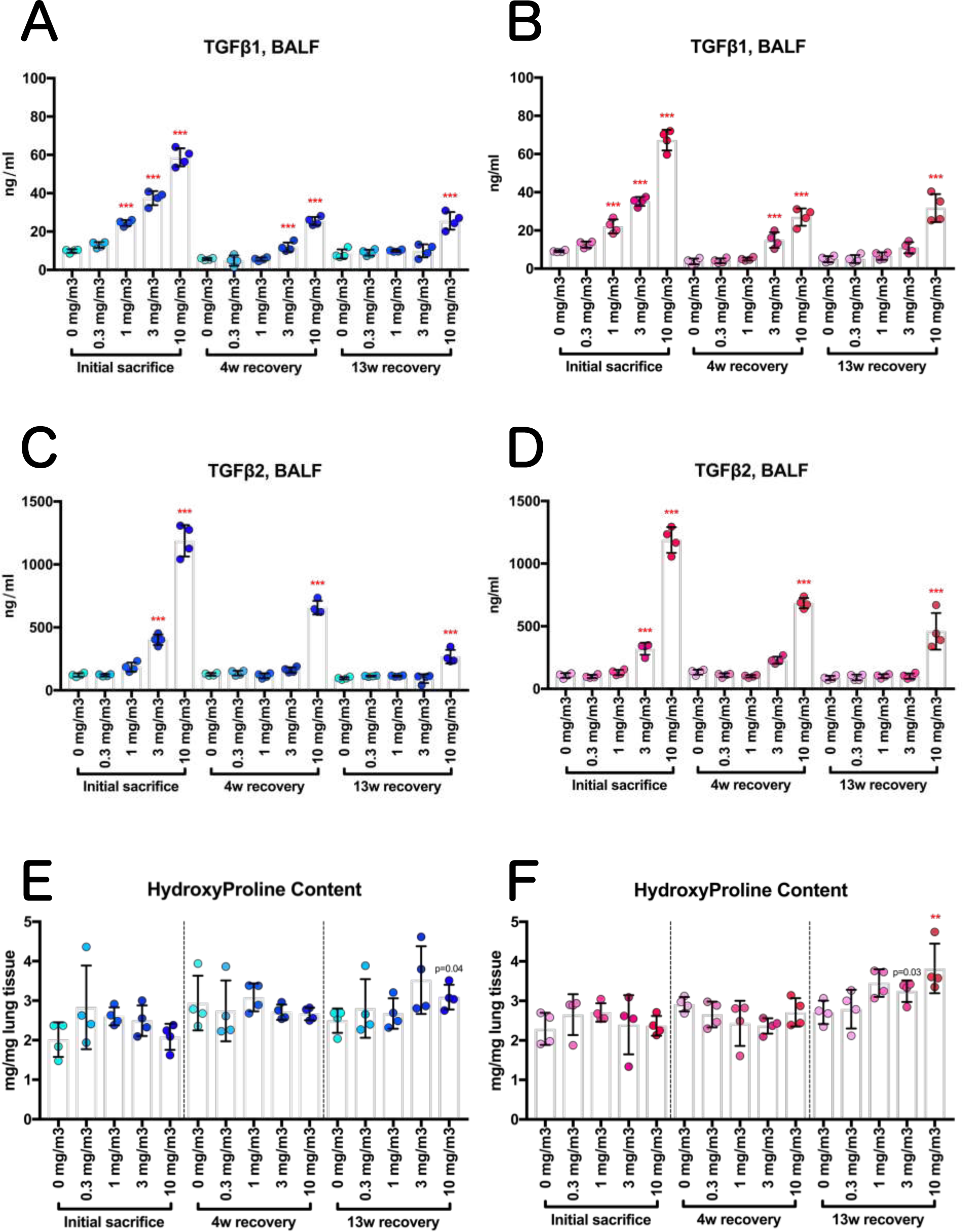
Induction of lung levels of transforming growth factor (TGF) β and hydroxyproline, a major component of collagen fibers, in rats treated with CWAAP-A by inhalation for 13 weeks. The concentrations of TGFβ1 and TGFβ2 in BALF were measured by enzyme immunoassay, and represented in panels A and C for males and in panels B and D for females. Hydroxyproline content in the lungs of males (E) and females (F) were determined using a commercially available colorimetric kit. William’s multiple comparison test compared with age-matched control (0 mg/m^3^) groups: \**p*<0.025, \*\**p*<0.005 and \*\*\**p*<0.0005.

Increased hydroxyproline content in the lung is a result of the fibrotic process and consequently did not follow the same pattern as TGFβ1 and TGFβ2. Hydroxyproline content was not significantly increased immediately after the end of the exposure period or after the 4 week recovery period. Hydroxyproline content did have a tendency to be increased in the higher exposure concentration groups after the 13-week recovery period and in the female rats exposed to 10 mg/m^3^ CWAAP-A hydroxyproline content was significantly increased (Fig. 13E and F).

These results indicate that systemic inhalation exposure to CWAAP at concentrations ranging from 0.3 to 10 mg/m^3^ for 6 hours per day, 5 days per week, for 13 weeks, resulted in toxicity in the lungs in a concentration-dependent manner. Furthermore, in both the male and female 3 mg/m^3^ and 10 mg/m^3^ exposure groups, after the 13-week recovery period, partial progression of lung pathology was observed.

### CWAAP-induced acute injury in rat lungs was attenuated by depletion of alveolar macrophages

Similarly to our recent study [5], inhalation exposure to CWAAP-A caused alveolar injury with infiltration of inflammatory cells into rat alveoli during the acute phase (Table 2, and Fig. 10). Therefore, we investigated the effect of elimination alveolar macrophages or neutrophils on CWAAP-A induced lung injury. Clodronate liposome and PMN-neutralizing antibody were used to deplete macrophages and neutrophils, respectively. The experimental protocol is shown in Figure S6: two days prior to CWAAP-A exposure rats were treated with clodronate liposome or control liposome; the next day rats were treated with PMN-neutralizing antibody or normal rabbit serum; rats were then exposed to 10 mg/m^3^ CWAAP-A for 6 hours on days 2 and 3. The next day after the final exposure, BALF and plasma were collected and analyzed. In rats without CWAAP-A exposure, intratracheal administration of clodronate liposome disrupted alveolar macrophages and caused an increase in neutrophils in the BALF, while PMN antibody-treatment had little effect (Fig. 14A). In rats exposed to CWAAP-A, administration of clodronate liposome suppressed both macrophages and neutrophils in the BALF, while PMN antibody-treatment had less of an effect (Fig. 14B). Cell count analysis confirmed these observations. Clodronate liposome treatment significantly decreased the number of alveolar macrophages in the BALF of both unexposed rats and in rats exposed to CWAAP. Clodronate liposome treatment also increased neutrophils in the BALF of unexposed rats and decreased neutrophils in the BALF of CWAAP-A exposed rats. PMN antibody administration caused a decrease in white blood cell count along with a dramatic decrease in neutrophil count in the blood. PMN antibody administration also caused a decrease in monocyte, lymphocyte, and eosinophil counts in the blood (Fig. 15 and Fig. S5). These results indicate that treatment with PMN-neutralizing antibody was effective in decreasing total neutrophil count, but had little effect on neutrophils migrating from the intravascular space to the alveolar air space in this experimental system. These results also mirror the results obtained from the 13 week exposure to 10 mg/m^3^ CWAAP-A: neutrophils move into the alveolar air space in response to exposure to CWAAP-A. Finally, to evaluate the effect of macrophage depletion on CWAAP-A induced acute lung inflammation, plasma SP-D levels, a marker for interstitial pneumonia, were measured. The results showed that CWAAP-A exposure induced a significant increase in plasma SP-D levels and pretreatment with clodronate liposome significantly reduced this increase (Fig. 15G). Thus, CWAAP-A induced pulmonary disorders in rats were attenuated by depletion of alveolar macrophages. On the other hand, clodronate liposome-treated rats that were not exposed to CWAAP-A and clodronate liposome-treated rats that were exposed to CWAAP-A had significantly higher LDH activity and SP-D levels in the BALF compared to rats their respective untreated controls (Fig. S5A and S5B), indicating that these markers, which are both elevated by CWAAP-A exposure, are also elevated by clodronate-induced macrophage damage.

**Figure 14.**
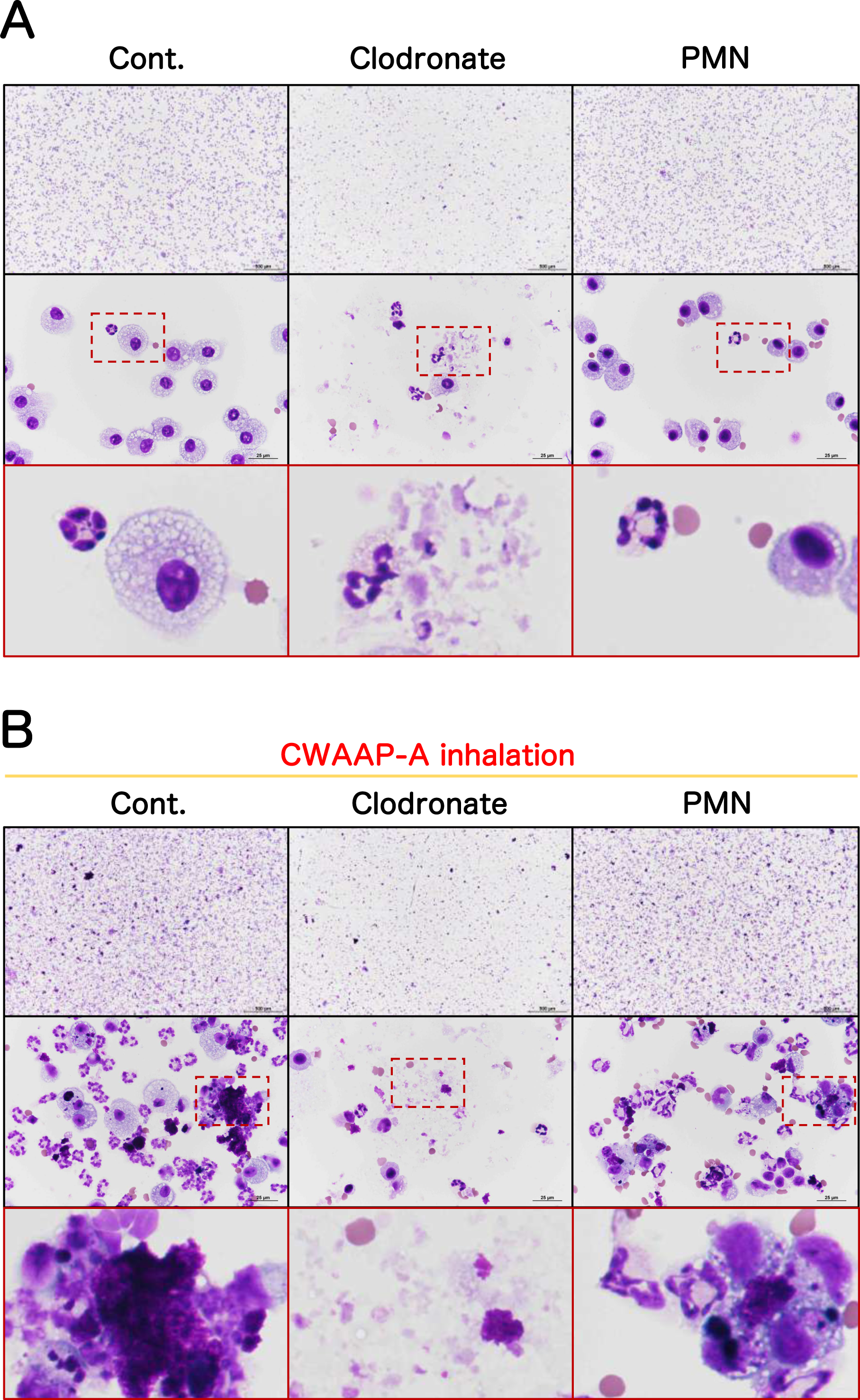
The effect of pretreatment of the CWAAP-A-exposed rats with clodronate liposomes (to deplete macrophages) or polymorphonuclear leukocytes (PMN)-neutralizing antibodies (to deplete neutrophils) on BALF cells. Male rats were given a single intratracheal dose of clodronate liposomes (1 ml/kg), or a single intravenous injection of PMN antibodies (0.5 ml/kg). They were then exposed to 10 mg/m^3^ of CWAAP-A for 6 hours and sacrificed the next day for analysis. The results of BALF cytospin cytology without and with CWAAP-A exposure are shown in A and B, respectively.

**Figure 15.**
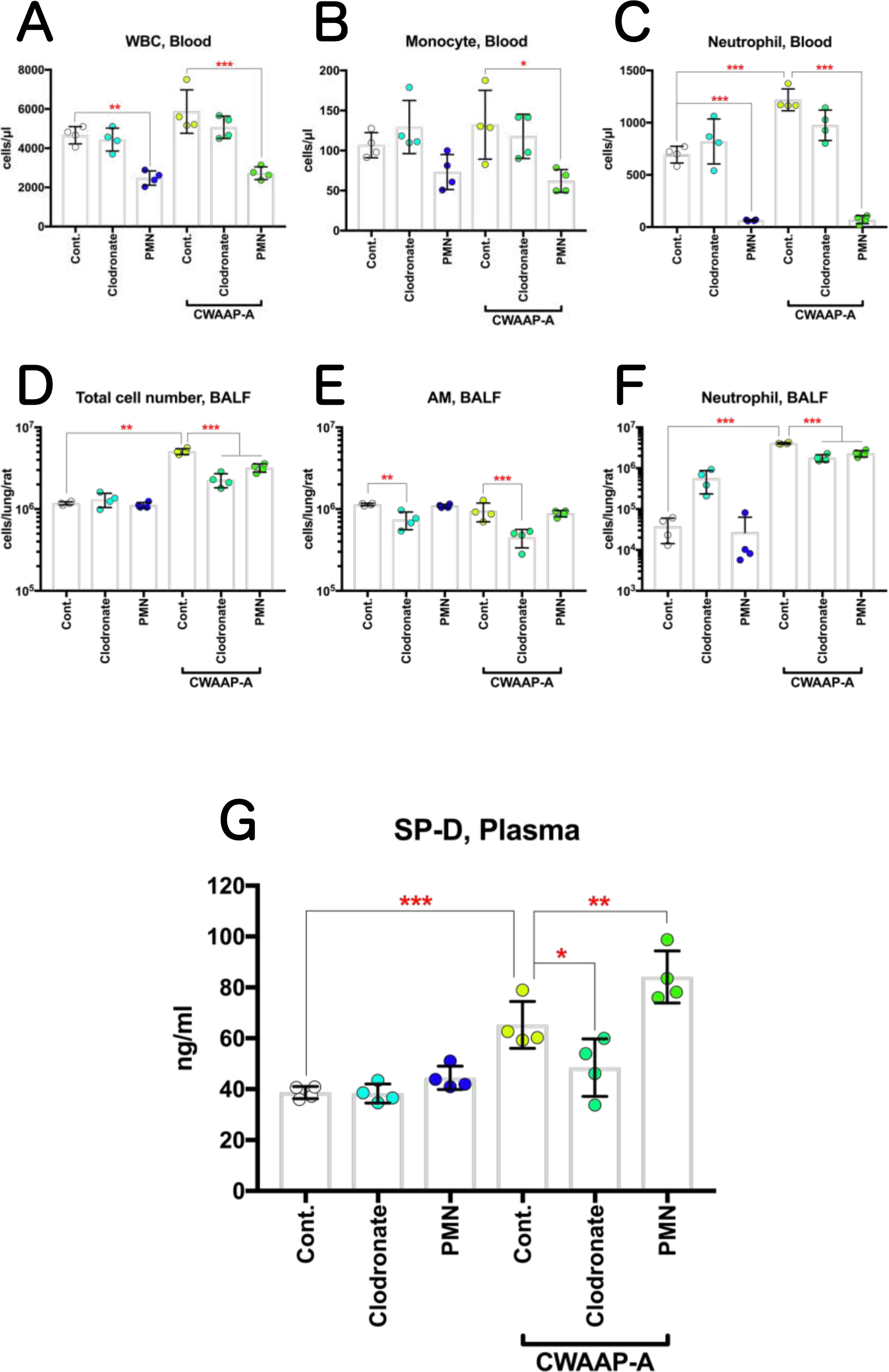
Recovery of the pulmonary acute toxicity of CWAAP-A by pretreatment of the CWAAP-A-exposed rats with clodronate liposome rather than PMN-neutralizing antibody. Male rats were given a single intratracheal dose of clodronate liposomes (1 ml/kg), or a single intravenous injection of PMN antibodies (0.5 ml/kg). They were then exposed to 10 mg/m^3^ of CWAAP-A for 6 hours, and their plasma and BALF were collected the next day. The number of white blood cells (WBC) (A), monocytes (B) and neutrophils (C) in plasma, and the number of total cells (D), alveolar macrophage (AM) (E) and neutrophils (F) in BALF were measured by an automated hematology analyzer. The plasma concentration of SP-D was determined by an enzyme immunoassay (G). Tukey’s multiple comparison test: \**p*<0.05, \*\**p*<0.01, and \*\*\**p*<0.001, pairs indicated.

**Figure 16.**
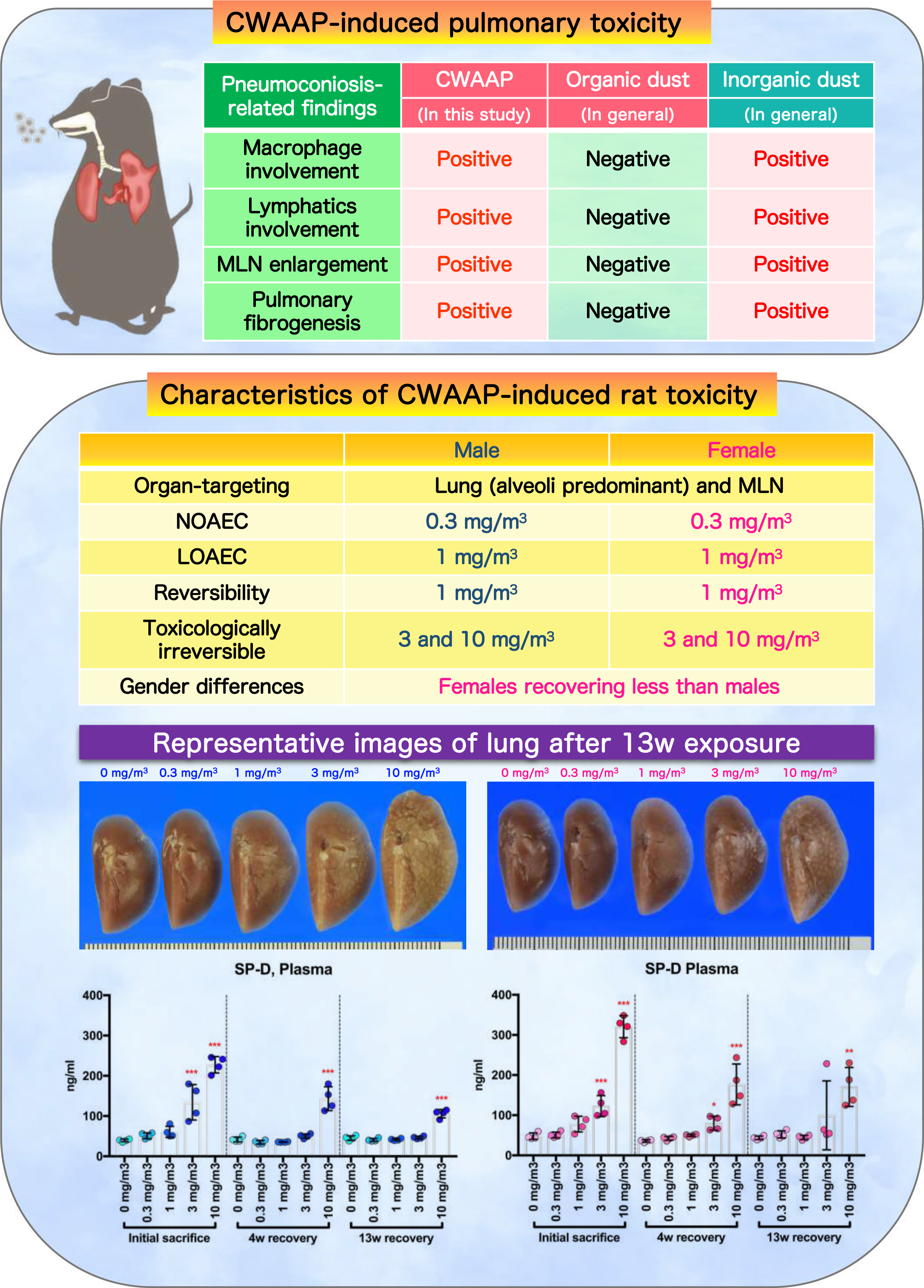
Graphical abstract in this study.

## Discussion

We have previously reported the clinical pathology of occupational lung diseases caused by inhalation of CWAAP in a chemical plant in Japan [16] and our initial study of the effects in rat of exposure to CWAAP. In the rat study we used two exposure methods: whole body exposure and intratracheal instillation: rats were exposed by whole body inhalation to CWAAP-A concentrations of 15 and 40 mg/m^3^ for 4 hours once a week for 2 months or once to CWAAP-A concentration of 40 and 100 mg/m^3^ for 4 hours, or rats were administered CWAAP-A or CWAAP-B by intratracheal instillation at 1 mg/kg once every other week for 2 months [5]. We found that CWAAP caused alveolar lesions and that in the 40 mg/m^3^ group exposed CWAAP-A for 2 months these lesions developed into alveolitis with fibrous thickening of the alveolar septum. Intratracheal instillation caused qualitatively similar pulmonary pathology as rats exposed to CWAAP-A by inhalation. Based on these findings, and to obtain additional reliable systemic toxicity data for occupational health and disease, the present study was conducted in compliance with OECD TG 413. Male and female rats were exposed to 0, 0.3, 1, 3 and 10 mg/m^3^ CWAAP-A for 6 hours per day, 5 days per week, for 13 weeks. Evaluation of 40 organs in the rats demonstrated that damage caused by inhalation exposure to CWAAP-A was limited to alveolar lesions in the lungs with no effects in the upper respiratory tract, including the nasal cavity and pharynx. Alveolar lesions developed in both sexes in a dose-dependent manner. Histopathological diagnosis of the lungs on the day after the final exposure showed that granulomatous changes, which are inflammatory reaction, were observed in all rats exposed to 1, 3 and 10 mg/m^3^ CWAAP-A. The lowest observed adverse effect concentration (LOAEC) for CWAAP-A in this study was 1 mg/m^3^ for both sexes, and 0.3 mg/m^3^ was the no observed adverse effect concentration (NOAEC).

This study also used recovery periods of 4 and 13 weeks to examine the recoverability from the histopathological and clinicopathological changes caused by CWAAP-A inhalation. We found that statistically significant changes in many clinicopathological parameters disappeared after a 4-week recovery period in the 1 mg/m^3^ group and after a 13-week recovery period in the 3 mg/m^3^ group (Extended file 5). However, while proliferative lesions in alveolar epithelium and inflammatory changes in alveolar air spaces were completely restored in the 1 mg/m^3^ group after a 13-week recovery period, these lesions were not resolved in the 3 mg/m^3^ group. Thus, rats recovered from lung disorders caused by inhalation of 1 mg/m^3^ CWAAP-A for 13 weeks, but exposure to 3 mg/m^3^ and 10 mg/m^3^ CWAAP-A caused inflammatory and fibrotic changes that were not resolved even after a 13-week recovery period. Consequently, these lesions are considered to be toxicological “point of no return” or tipping points associated with adverse outcomes [5, 17]. Thus, 3 mg/m^3^ and 10 mg/m^3^ were the exposure concentrations identified in this study that induced irreversible and progressive lesions.

The present study using male and female rats also provided new insight into the sex differences in CWAAP-induced lung disorders. In the groups exposed to 10 mg/m^3^ CWAAP-A there was no obvious difference in pulmonary lesions immediately after the end of the exposure period. However, after the 13-week recovery period pathological findings, such as fibrous changes and lipoproteinous material deposition, tended to be more sever in females than in males. In agreement with this, restoration to normal plasma SP-D levels was also slower in females. These results suggest that there is a gender difference in lesion recovery and progression, with females having weaker recovery and more progressive disease than males.

Epidemiology studies have demonstrated that organic dusts cause respiratory diseases, such as byssinosis caused by cotton dust and farmer’s lung disease, a hypersensitivity pneumonitis caused by spores of hyperthermia actinomycetes [18–20]. In addition, there are case reports in which chlorides of acrylic acid, the smallest unit of CWAAP, and acrylic esters and derivatives, have caused allergic contact dermatitis, asthma, and respiratory hypersensitivity [21–23]. However, no epidemiological studies have linked inhalation of organic dust to the development of interstitial lung disease with fibrosis. Our clinicopathological and rodent studies [5, 16] showed that none of the CWAAP-exposed lungs showed bronchitis with eosinophilic involvement, which are common in allergic diseases [24–26], and no increase in eosinophils was observed in the BALF or blood of CWAAP-exposed rats. Furthermore, in the industrial accident at the chemical plant handling CWAAP, pneumoconiosis rather than occupational asthma was found to occur, and interstitial pneumonia with fibrous thickening around the alveoli was observed in rodents exposed to CWAAP. These results indicate that inhalation exposure to CWAAP is a novel risk factor for causing pneumoconiosis in both humans and rodents.

Our recent study found that exposure to CWAAP resulted in enlargement of the mediastinal lymph nodes in rats [5]. Similarly, in the current study, CWAAP inhalation resulted in the dose-dependent enlargement of the mediastinal lymph nodes in rats. In addition, we previously found that whole-body inhalation exposure of rodents to indium tin oxide [27–29] and multi-walled carbon nanotube (MWCNT) [30, 31] also caused enlargement of mediastinal lymph nodes, as well as lung diseases. In clinical practice, it has been reported that enlarged mediastinal lymph nodes are observed secondary to interstitial lung diseases such as idiopathic pulmonary fibrosis and pneumoconiosis [32, 33]. These data are in agreement with the conclusion stated above that exposure to CWAAP is a risk factor for pneumoconiosis.

The number of capillary lymphatic vessels in hotspot areas and multifocal lesions in CWAAP-A exposed lungs tended to increase compared to normal alveolar areas, and the collective lymphatic vessels in the bronchovascular bundle and hilar regions of the lung tended to dilate. This indicates that lymphangiogenesis occurred in the inflammatory lesions, and the increased collection of interstitial fluid due to inflammation may have caused the subsequent dilation of the collecting lymphatics. Baluk *et al.* demonstrated that the widespread lymphangiogenesis observed in bleomycin-induced murine pulmonary fibrosis ameliorated pulmonary fibrosis after lung injury [34]. Thus, the change in lymphatics seen in our study appear to have been in due to a repair response to CWAAP-A induced damage. In summary, lymphatic vessel changes may be a possible secondary etiology arising from lung lesions caused by CWAAP.

The experiment using clodronate liposomes to deplete macrophages demonstrated that the increased level of plasma SP-D resulting from CWAAP-A exposure was suppressed by decreasing alveolar macrophages prior to CWAAP-A inhalation. Plasma SP-D levels is a marker of various lung diseases including interstitial pneumonia marker: disease induced modifications of SP-D facilitate its leakage from the lung [35]. Thus, the findings that macrophage depletion decreased the leakage of SP-D out of the lung and into the plasma of CWAAP-A exposed rats indicate that alveolar macrophages contribute to CWAAP-A induced acute lung injury.

Clodronate liposome pretreatment increased neutrophils in the BALF of unexposed rats, as has been observed previously [36]. This effect is hypothesized to be due noninflammatory phagocytosis by alveolar macrophages of foreign bodies that are inspired during normal respiration, thereby cloaking them from resident neutrophils and preventing superfluous neutrophil recruitment into the healthy lung [37]. In CWAAP-A exposed lungs, clodronate liposome pretreatment resulted in a moderate but significant reduction of the increase in neutrophil count in the BALF caused by CWAAP-A exposure. Cytokine-induced neutrophil chemoattractant-1 (CINC-1) in rats is a key player in neutrophil chemoattractant to sites of inflammation and is a counterpart of human growth-related oncogene belonging to the interleukin-8 family [38]. Previous reports have shown that in a rat lung injury model induced by lipopolysaccharide (LPS) and characterized by neutrophil infiltration, alveolar macrophages responded to LPS and produced CINC-1 within 6 hours after administration of LPS [39–41]. These reports are consistent with our results that macrophage depletion attenuated increased neutrophil numbers in the lung.

PMN antibody pretreatment dramatically reduced the number of neutrophils found in the blood. However, the decrease in neutrophils in the BALF while significant was moderate. PMN antibody and clodronate liposome pretreated rats had similar numbers of neutrophils in the BALF; however, in contrast to clodronate liposome pretreated rats, PMN pretreatment did not reduce the levels of SP-D in the plasma, but rather the levels of SP-D in the plasma of PMN antibody treated rats exposed to CWAAP-A were increased. This suggests that while it is likely that neutrophils in the lung contributed to CWAAP-A induced acute injury, other macrophage associated factors also contributed to lung injury.

It is notable that while PMN antibody pretreatment reduced the number of neutrophils in the blood by more than 90%, there was no significant reduction of neutrophils in the lungs of unexposed rats and only a mild decrease in neutrophils in the lungs of CWAAP-A exposed rats. This suggests that the increase in neutrophil counts in the CWAAP-A exposed lung in rats pretreated with PMN antibodies depend more on proliferation of tissue resident neutrophil in the lung [42] than on infiltration of neutrophils from outside the lung. To define the role of neutrophils in CWAAP-A induced acute lung damage more definitively, future studies will need to be carried out, including improved neutrophil removal. A recent study reports the development of a double antibody-based depletion strategy [43], and this can be tested in future work.

## Conclusions

Inhalation exposure to CWAAP-A had harmful effects only in the alveoli. In this study, the LOAEC for pulmonary disorders in male and female F344 rats was 1 mg/m^3^ and the NOAEC was 0.3 mg/m^3^. In addition, rats of both sexes were able to recover from the tissue damage caused by 13 weeks exposure to 1 mg/m^3^ CWAAP-A, however, tissue damage caused by exposure to 3 and 10 mg/m^3^ were irreversible due to the development of interstitial lung lesions. There was a sex difference in the recovery of lung lesions, with females showing less recovery than males. Comparison of CWAAP-A, MWCNT-7, and titanium dioxide revealed that LDH activity in rat BALF after inhalation exposure to CWAAP-A (this study) was higher than that of MWCNT-7 and titanium dioxide [30, 44] (Table S1). These results suggest that inhalation of CWAAP-A is more harmful than inhalation of MWCNT-7 and titanium dioxide: both MWCNT-7 and titanium dioxide are classified as group 2B carcinogens by IARC. Finally, acute lung lesions caused by CWAAP-A exposure were significantly reduced by depletion of macrophages in the lungs.

## Methods

### Materials

CWAAP was purchased from a company which produces acrylic acid polymers. This product is the same as the one we used in our previous study (defined as CWAAP-A) [5]. A list of all primary antibodies used in these studies is summarized in Table S2. The donkey anti-mouse IgG conjugated Alexa Fluor 594 (ab150112), anti-goat IgG conjugated Alexa Fluor 594 (ab150136), anti-rabbit IgG conjugated Alexa Fluor 594 (ab150064) and goat anti-chicken IgY conjugated Alexa Fluor 488 (ab150169) were purchased from Abcam plc (Cambridge, UK). The VECTASHIELD Mounting Medium with DAPI (H-1200) was purchased from Vector laboratories (Burlingame, CA). The other reagents were of the highest grade commercially available.

### Animals

Male and female F344 rats at 6-7 weeks old were purchased from Charles River Laboratories Japan, Inc. (Yokohama, Japan). The rats were housed in an air-conditioned room under a 12 hours light/12 hours dark (8:00-20:00, light cycle) photoperiod, and fed a general diet (CRF-1, Oriental Yeast Co. Ltd., Tokyo, Japan) and tap water *ad libitum*. After approximately 1-2 weeks of quarantine and acclimation, they were exposed to CWAAP-A. All animal experiments were approved by the Animal Experiment Committee of the Japan Bioassay Research Center.

### Generation of CWAAP-A aerosol

The generation of CWAAP-A aerosol into the inhalation chamber was performed using our established method (cyclone sieve method) [31, 45] with some modifications to optimize it for CWAAP-A aerosol generation. Briefly, CWAAP-A was fed into a dust feeder (DF-3, Shibata Scientific Technology, Ltd., Soka, Japan) to generate CWAAP-A aerosol, and then introduced into a particle generator (custom-made by Seishin Enterprise Co., Ltd., Saitama, Japan) to separate the aerosol and feed it into the inhalation chamber. The concentration of the CWAAP-A aerosol in the chamber was measured and monitored by an optical particle controller (OPC; OPC-AP-600, Shibata Scientific Technology), and the operation of the dust feeder was adjusted by feedback control based on upper and lower limit signals to maintain a steady state.

The gravimetric mass concentration of CWAAP aerosol in the chamber was measured every two weeks during the exposure period. Aerosols collected on a fluoropolymer binder glass fiber filter (TX40HI20-WW, φ55 mm, Tokyo Dylec, Corp., Tokyo, Japan) were weighed for each target concentration at 1, 3, and 5 hours after the start of the exposure. Using the mass per particle (K-value) calculated using the measured mass results (mg/m^3^) and the particle concentration data (particles/m^3^) obtained from the OPC, the particle concentration for each group during the exposure period was converted to a mass concentration (shown as calibrated mass concentration in Table 1). The particle size distribution and morphology of the CWAAP particles were measured at the first, sixth, and last week of exposure. Particle size distribution was measured using a micro-orifice uniform deposit cascade impactor (MOUDI-II, MSP Corp., Shoreview, MN). The MMAD and GSD were calculated by cumulative frequency distribution graphs with logarithmic probability (Fig. S1C and Table 1). CWAAP-A particles in the inhalation chamber were collected on a 0.2 μm polycarbonate filter (φ47 mm, Whatman plc, Little Chalfont, UK), and observed using SEM (SU8000, Hitachi High-Tech, Tokyo, Japan).

### 13-week inhalation study

This experiment was conducted with reference to the OECD Guideline for Testing of Chemicals (TG 413) [6]. The sodium salt of cross-linked acrylic acid polymers has been reported to cause collagen deposition in the lungs after inhalation of 10 mg/m^3^ for 13 weeks [46]. Also, at the site where the incident occurred, the 8h-Time-Weighted-Average for personal exposure concentrations of inhalable CWAAP dust was 0.4 to 7.6 mg/m^3^ [16]. Based on these reports and the OECD TG 413, target concentrations for CWAAP aerosols were set at 0.3, 1, 3 and 10 mg/m^3^, and the exposure schedule was 6 hours per day, 5 days per week, for 13 weeks (Fig. S6A). Rats were divided into 5 exposure groups of 48 rats (24 males and 24 females) in each group, and 8 males and 8 females of each group were euthanized at 1 hour, 4 weeks, and 13 weeks after the last exposure. Rats were transferred to individual stainless steel cages and exposed to 0, 0.3, 1, 3, or 10 mg/m^3^ CWAAP-A for 6 hours without food or water. All inhalation chambers were maintained at an air exchange rate of 7-9 times/hour, a temperature of 21-25°C, and a humidity of 30-70% during the study period (see Table 1). After exposure, the rats were returned to the stainless steel bedding cages and kept in groups with free access to food and water. During the study period, body weight and food consumption of the rats were measured once a week. At 1 hour, 4 weeks, and 13 weeks after the last exposure, the blood of the rats was collected under isoflurane anesthesia and rats were euthanized by exsanguination. BALF was collected from four males and four females in each group as described below. The lungs from which BALF was not collected were used for hydroxyproline assay. For histopathological analysis, all tissues were collected from the remaining four males and four females in each group, and fixed in 10% neutral phosphate buffered formalin solution.

### Macrophage or neutrophil depletion

Male F344 rats (8 weeks old) were divided into 6 groups of 4 rats each. Alveolar macrophages and neutrophils were depleted using clodronate liposome [8] and PMN-neutralizing antibody [9], respectively. Rats (8 weeks old) were administered a single intratracheal dose of clodronate liposome or control liposome (LIPOSOMA, Inc., Amsterdam, The Netherlands) at a dose of 1 ml/kg. The next day, PMN-neutralizing antibody (Accurate Chemical Scientific Corp., Carle Place, NY) or normal rabbit serum (Cedarlane Laboratories, Inc., Burlington, Canada) was administered intravenously at a dose of 0.5 ml/kg (Fig. S6B). The PMN antibody and normal rabbit serum were previously diluted 5-fold in PBS and administered at a volume of 2.5 ml per kg of body weight. The day after administration of PMN antibody, inhalation exposure to 10 mg/m^3^ CWAAP-A was begun. Rats were placed in the inhalation chamber and exposed to CWAAP at a concentration of 10 mg/m^3^ for 6 hour/day for 2 days. Next day after the last exposure, animals were euthanized by exsanguination under isoflurane anesthesia, and BALF was collected for analyses (see below for details).

### BALF collection and analysis

For the 13-week inhalation study, the right bronchus was tied with a thread, and the left lung was lavaged. 4∼7ml of saline was injected into the lung through the trachea, in and out twice, and collected as BALF. In the macrophage or neutrophil depletion experiment, 6.2 ml of saline was injected into all lungs through the trachea, in and out twice, and collected as BALF. The total cell numbers in the BALF were counted using an automatic cell analyzer (Sysmex Corp., Automated Hematology Analyzer, XN-2000V). Thin-layer cell populations on glass slides were prepared using Cytospin 4 (Thermo Fisher Scientific, Inc., Waltham, MA). After May-Grunwald-Giemsa staining, differential white blood cell count was conducted by visual observation. BALF was centrifuged at 1,960 rpm (800 × *g*) for 10 min at 4°C, and the activity of LDH, ALP and γ-GTP and the phospholipid level in the supernatant was measured using an automatic analyzer (Hitachi 7080, Hitachi, High-Tech Corp., Tokyo, Japan).

### Hematological and blood chemistry tests

For hematological examination, blood samples were analyzed with an automated hematology analyzer (Sysmex). For biochemical tests, the blood was centrifuged at 3,000 rpm (2,110 × *g*) for 20 minutes, and the supernatant was analyzed with an automated analyzer (Hitachi High-Tech).

### Enzyme immunoassay

SP-D concentrations in BALF supernatants and plasma were determined using Rat/Mouse SP-D kit YAMASA EIA (YAMASA Corp., Choshi, Japan). In this assay, BALF was diluted 500- or 1,000-fold, and plasma was diluted 25-fold with the assay diluent included in the kit. BALF concentrations of TGFβ1 and TGFβ2 were measured by Human/Mouse/Rat/Porcine/Canine TGF-beta 1 Quantikine ELISA Kit (R&D Systems) and Mouse/Rat/Canine/Porcine TGF-beta 2 Quantikine ELISA Kit (R&D Systems). For the TGFβ2 assay, BALF was diluted 3-fold with the sample diluent included in the kit. The absorbance at 450 nm was measured using a microplate reader (Spark^®^; Tecan Group, Ltd., Männedorf, Switzerland or SpectraMax; Molecular Devices, LLC., San Jose, CA). For TGFβ1 and TGFβ2 assays, the absorbance at 570 nm was also measured and subtracted as a background signal.

### Hydroxyproline assay

The lung content of hydroxyproline, a major component of collagen fibers, was determined using a hydroxyproline assay kit (Perchlorate-Free) (BioVision, Inc., San Francisco, CA). Small pieces of lung (approximately 30 mg) were homogenized in 10 volumes of distilled water using a portable power homogenizer (ASONE Corp., Osaka, Japan). The resulting homogenates were mixed with an equal volume of 10 M NaOH, and incubated at 120°C for 1 hour. After neutralization by addition of 10 M HCl, the mixture was centrifuged at 10,000 × *g* for 5 minutes. 10 μl of the supernatant was transferred to a 96-well plate and dried at 65°C in a hybridization incubator (TAITEC, Corp., Koshigaya, Japan). The hydroxyproline signal (absorbance at 560 nm) was measured using a microplate reader (Spark^®^; Tecan Group, Ltd., Männedorf, Switzerland or SpectraMax; Molecular Devices, LLC., San Jose, CA) according to the kit instructions.

### Histopathological analysis

Serial tissue sections were cut from paraffin-embedded lung specimens, and the first section (2-μm thick) was stained with H&E for histological examination and the remaining sections were used for immunohistochemcal analysis. The histopathological findings for lung and mediastinal lymph node in this study were determined after multifaceted discussions between certified pathologists from the Japanese Society of Toxicologic Pathology and certified medical pathologists from the Japanese Society of Pathology, based on the definitions of terms adopted by International Harmonization of Nomenclature and Diagnostic Criteria for Lesions in Rats and Mice (INHAND)[47]. Pathological diagnosis was performed blindly by three pathologists and summarized as a result of the discussion.

### Alcian blue staining

Details of the method have been described previously [5]. Briefly, after deparaffinization and rinsing, the slides were incubated in 0.1M HCl solution for 3 minutes. Then, they were incubated in alcian blue staining solution (alcian blue 8GX, C.I.74240, Merck-Millipore, Burlington, MA) for 10 minutes at room temperature. The slides were then lightly passed through 0.1N HCl solution and washed under running water, followed by contrast staining with Kernechtrot (NUCLEAR FAST RED, C.I.60760, Merck-Millipore) for 5 minutes. After rinsing, the samples were dehydrated, permeabilized, and sealed.

### Immunohistological multiple staining analyses

Details of the multiple staining method have been described previously [5]. Briefly, lung tissue sections were deparaffinized with xylene and hydrated through a graded ethanol series. For immunohistochemical staining, sections were incubated with 0.3% hydrogen peroxide for 10 minutes to block endogenous peroxidase activity, then incubated with 10% normal serum at room temperature (RT) for 10 minutes to block background staining, and then incubated for 2 hours at RT with the primary antibodies (VEGFR3 or GFP). After washing with PBS, the sections were incubated with histofine simple stain kit Rat MAX-PO(G) (414331F, Nichirei, Tokyo, Japan) or goat anti-chicken IgY conjugated biotin (ab6876, abcam plc, Cambridge, UK) for 30 minutrs at RT. After further washing with PBS, sections were incubated with DAB EqV Peroxidase Substrate Kit, ImmPACT (SK-4103, Vector laboratories) for 2-5 minutes at RT for colorization. Importantly, after washing with dH_2_O after color detection, the sections were treated with citrate buffer at 98°C for 30 minutes before reacting with the next primary antibody to denature the antibodies bound on the sections used. The second antibody was then reacted in the same way as for the initial staining procedure. Histogreen chromogen (AYS-E109, Cosmo Bio, Tokyo, Japan) was used for the second coloration, followed by hematoxylin staining for 30-45 seconds as a contrast stain. The sections were dehydrated and sealed. For immunofluorescence staining, all primary antibodies used were made into a cocktail for the staining step and used simultaneously. The sections were then incubated with secondary antibodies conjugated with Alexa Fluor 594. Sections were thoroughly washed and incubated with secondary antibody conjugated with Alexa Fluor 488. After the fluorescence-labeled secondary antibody reactions, the sections were shielded with DAPI-containing encapsulant and used for imaging. The sections were observed under an optical microscope ECLIPSE Ni (Nikon Corp., Tokyo, Japan) or BZ-X810 (Keyence, Osaka, Japan).

### Statistical analysis

Except in the case of incidence and integrity of histopathological lesions, the data comparisons among multiple groups were performed by one-way analysis of variance with a post-hoc test of William’s test or Tukey’s multiple comparison test using Pharmaco Basic Ver. 16.1 (Scientist Co., Tokyo, Japan) or GraphPad Prism 5 (GraphPad Software, San Diego, CA), respectively. The incidences and integrity of lesions were analyzed by chi-square test using GraphPad Prism 5 (GraphPad Software, San Diego, CA). Statistical significance was set at *p* < 0.025 for William’s test and *p* < 0.05 in the other cases.

## Abbreviations

BALF: Bronchoalveolar lavage fluid
CINC-1: Cytokine-induced neutrophil chemoattractant-1
CWAAP: Cross-linked water-soluble acrylic acid polymer
DAPI: 4’,6-diamino-2-phenylindole
EGFP: Enhanced green fluorescence protein
GSD: geometric standard deviation
HE: Hematoxylin and eosin
LDH: Lactate dehydrogenase
LOAEC: Lowest Observed Adverse Effect Concentration
LPS: Lipopolysaccharide
Lyve-1: Lymphatic vessel endothelial hyaluronan receptor 1
MHLW: Ministry of Health, Labour and Welfare
MMAD: Mass median aerodynamic diameter
MWCNT: multi-walled carbon nanotube
NOAEC: No Observed Adverse Effect Concentration
PMN: Polymorphonuclear leukocytes
Prox1: Prospero homeobox protein-1
SEM: Scanning electron microscope
SP-D: Surfactant protein D
TGF: Transforming growth factor
VEGFR3: Vascular endothelial growth factor receptor 3

## Declarations

### Ethics approval and consent to participate

All animals were treated humanely and all procedures were performed in compliance with the Animal Experiment Committee of the Japan Bioassay Research Center.

### Consent for publication

All authors gave their consent for publication of this manuscript.

### Availability of data and materials

The datasets used and analyzed during the current study are available from the corresponding authors on reasonable request.

## Acknowledgments

We would like to express our gratitude to the Collaborative Research External Evaluation Committee of the Japan Organization of Occupational Health and Safety and Hisao Yamaguchi, Manager, Masako Kiguchi, Director, Hirohide Ohnishi, Director and Dr. Shigeki Koda, Acting Director of the National Institute for Occupational Safety and Health, for their valuable advice in the planning and implementation of this study. In addition, we wish to thank Dr. David B. Alexander of Nanotoxicology project, Nagoya City University Graduate School of Medicine for his insightful comments and English editing. We are deeply grateful to Drs. George Daghlian and Young-Kwon Hong of the Division of Plastic and Reconstructive Surgery, Department of Surgery, Norris Comprehensive Cancer Center, Keck School of Medicine, University of Southern California for providing us with the properly prepared lungs of Prox1-EGFP rats. We are also deeply grateful to Drs. Yasuhiro Yoshimatsu and Masanori Hirashima of the Division of Pharmacology, Niigata University Graduate School of Medical and Dental Sciences for helpful discussions and comments on the lymphatics of the lungs.

## Author information

Tomoki Takeda and Shotaro Yamano contributed equally to this work.

Japan Bioassay Research Center, Japan Organization of Occupational Health and Safety, Kanagawa, Japan

Tomoki Takeda, Shotaro Yamano, Yuko Goto, Shigeyuki Hirai, Yusuke Furukawa, Yoshinori Kikuchi, Kyohei Misumi, Masaaki Suzuki, Kenji Takanobu, Hideki Seno, Misae Saito, Hitomi Kondo, Yumi Umeda

Department of Surgery, Norris Comprehensive Cancer Center, Keck School of Medicine, University of Southern California, Los Angeles, CA, USA George Daghlian, Young Kwon Hong

Division of Pharmacology, Niigata University Graduate School of Medical and Dental Sciences, Niigata, Japan Yasuhiro Yoshimatsu, Masanori Hirashima

Department of Pathology, Tenri Hospital, Nara, Japan

Yoichiro Kobashi

Department of Pathology, Hokkaido Chuo Rosai Hospital, Japan Organization of Occupational Health and Safety, Hokkaido, Japan

Kenzo Okamoto

Director of Research and Training Center for Asbestos-Related Diseases, Okayama, Japan

Takumi Kishimoto

## Contributions

T.T. and S.Y. performed the experiments and analyzed the data. Y.G., S.H., Y.F., Y.K., K.M. and M.S. assisted with animal experiments. K.T., H.S., Y.U., and S.Y. performed histopathological diagnoses. S.Y., Y.U., Y.K., and K.O. conducted a pathological conference. M.S. and H.K. performed BALF sampling and dissection. G.D., Y.K.H., Y.Y. and M.H. performed the analysis of the normal pulmonary lymphatic vessel. Y.K. K.O. and T.K. analyzed and interpreted the data. T.T. and S.Y. conceived, designed, and directed the study and interpreted the data. T.T., S.Y., and Y.U. drafted and revised the manuscript. All authors approved the manuscript as submitted.

### Corresponding authors

Correspondence to Tomoki Takeda or Shotaro Yamano

### Supplementary Information

**Additional file 1: Figure S1.**
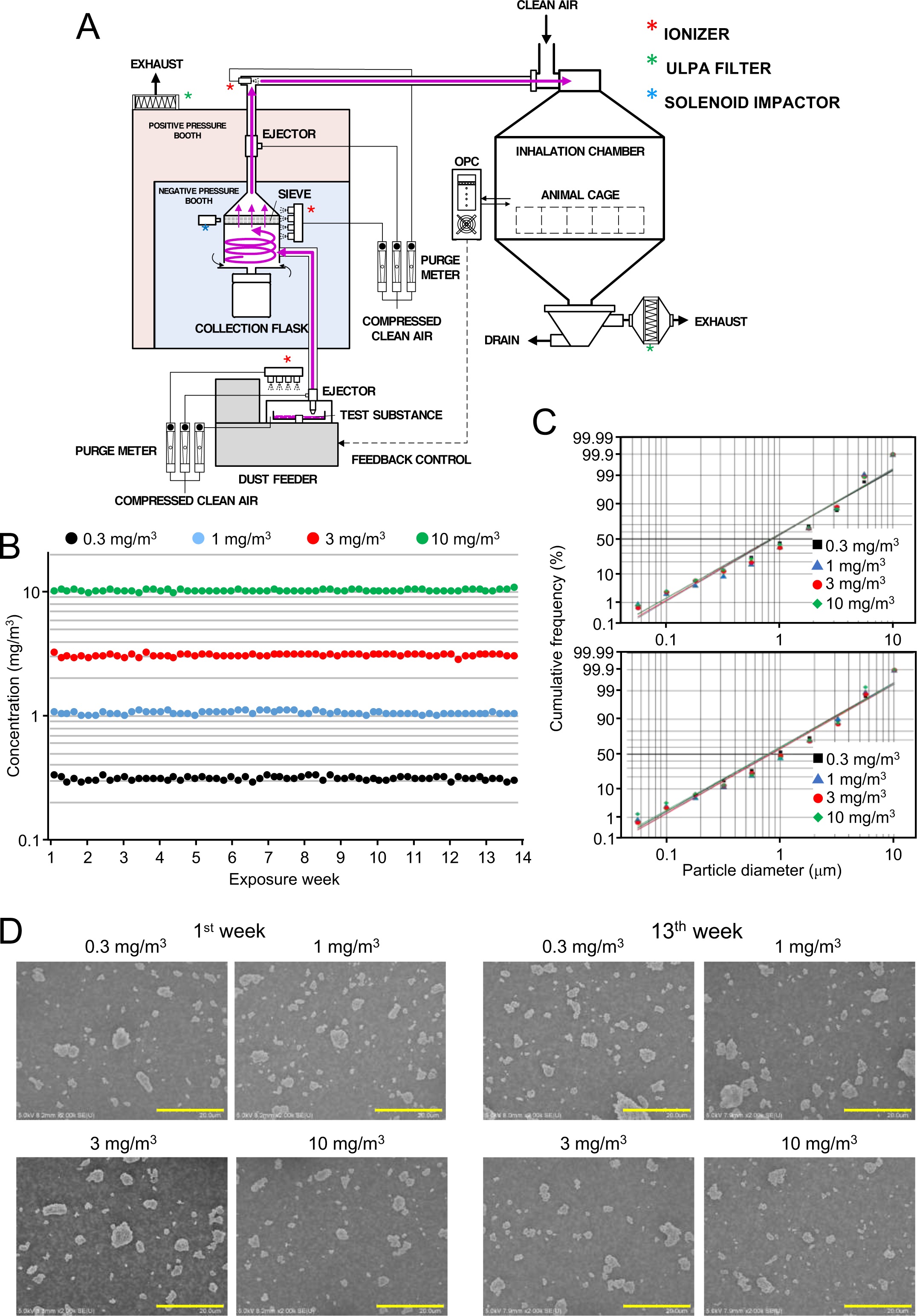
The whole body inhalation exposure system (A), the averaged CWAAP-A concentration in the chamber per each exposure day (B), cumulative frequency distribution graphs with logarithmic probability (C) and representative scanning electron microscope (SEM) images of the CWAAP-A particles in the chambers (D). Scale bar: 20 μm (panel D).

**Additional file 2: Figure S2.**
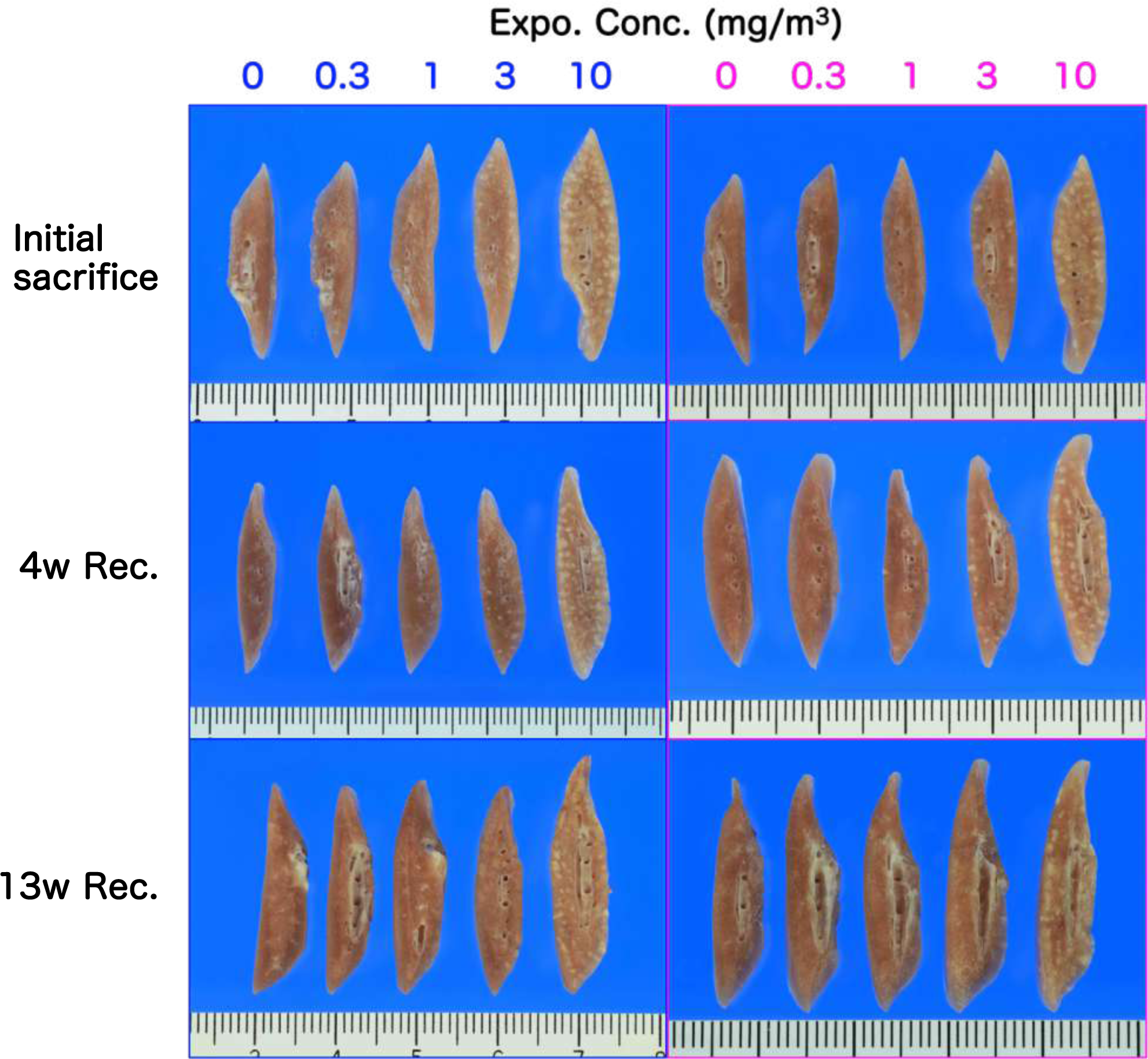
Representative macroscopic photographs of cross-sections of rat lungs. The left and right sides represent males and females, respectively. Abbreviations: conc, concentration; expo, exposure; and rec, recovery.

**Additional file 3: Figure S3.**
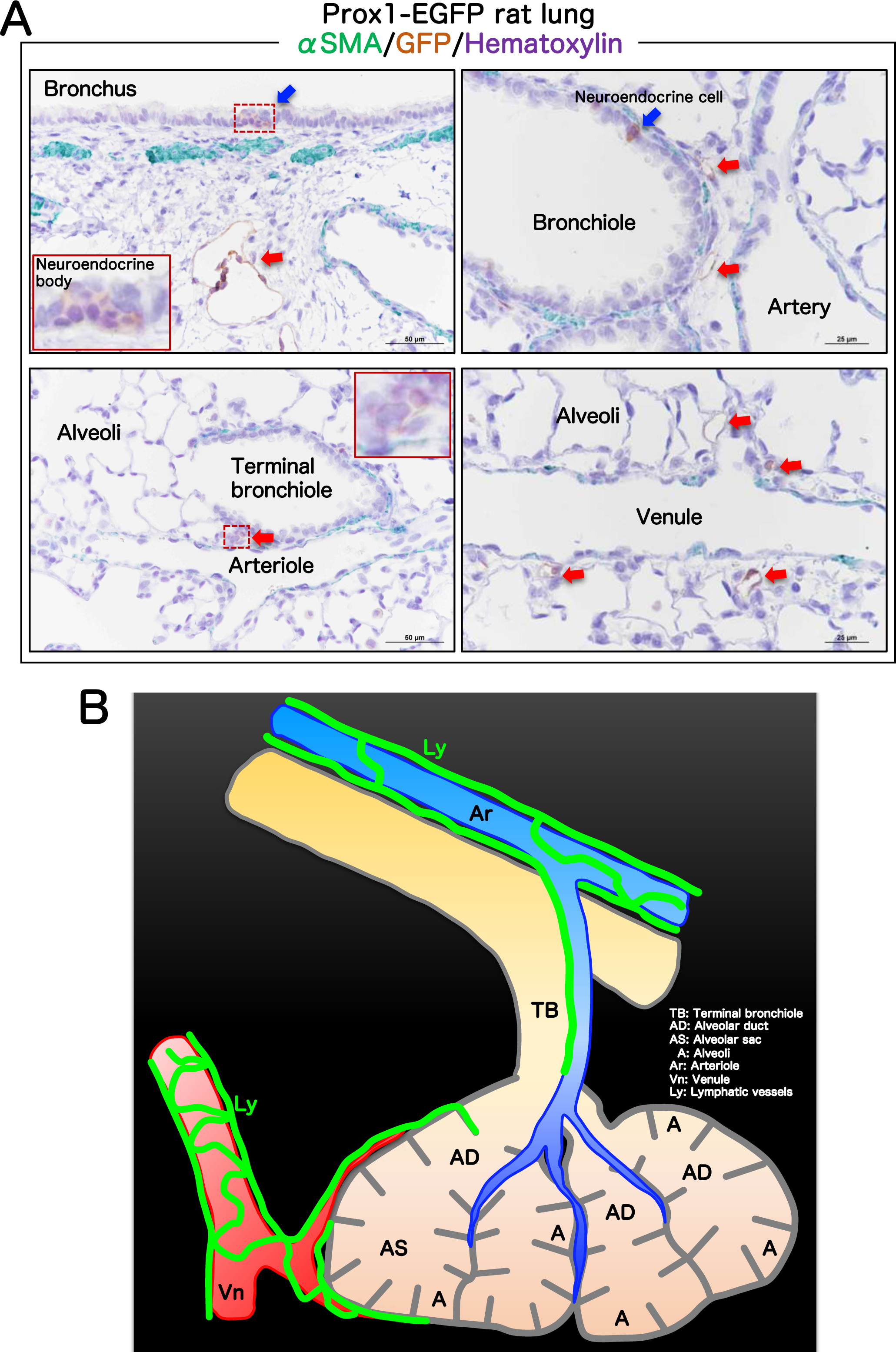
Visualization of normal lymphatic vessels running using the lungs of Prospero homeobox protein-1 (Prox1)-enhanced green fluorescent protein (EGFP) transgenic rats [13]. (A) Lung sections from Prox1-EGFP transgenic rats were double-stained with GFP and α-smooth muscle actin (αSMA) and counterstained with hematoxylin. Lymphatic vessels (red arrows) run around the veins and arteries. Blue arrows represent neuroendocrine cells. (B) A graphical image of lymphatic vessels running (green lines) through the lungs.

**Additional file 4: Figure S4.**
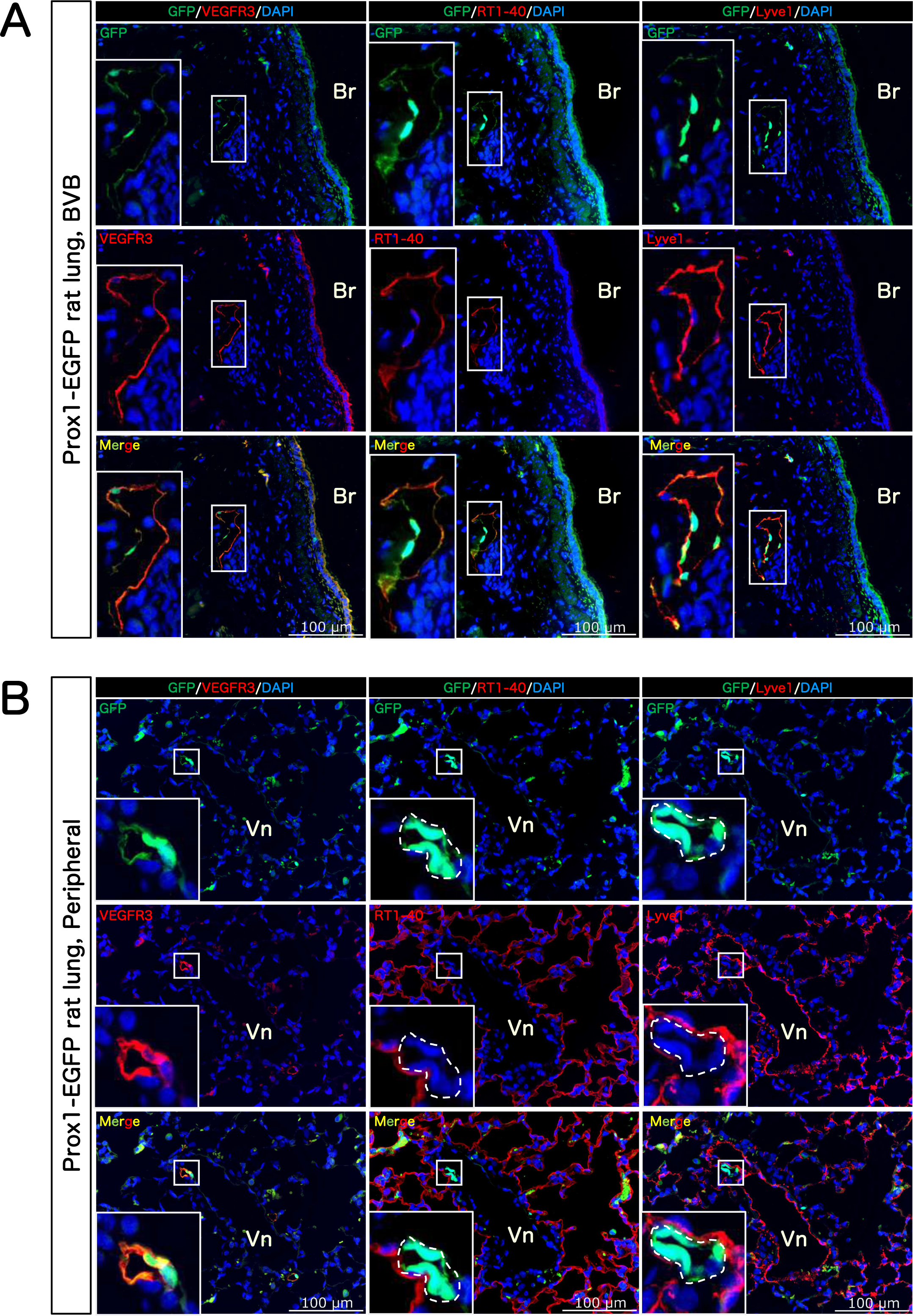
Co-localization of vascular endothelial growth factor receptor 3 (VEGFR3) with lymphatic vessels in rat lungs. Lung sections from Prox1-EGFP transgenic rats were stained with three commercially available lymphatic markers, VEGFR3, lymphatic vessel endothelial hyaluronan receptor 1 (Lyve-1) and podoplanin (RT1-40), under co-staining of GFP and 4’,6-diamidino-2-phenylindole (DAPI; a nucleus marker). Only VEGFR3 co-localized almost completely with GFP in rat lungs including peripheral regions. Abbreviations: Br, Bronchus and Vn, Venule.

**Additional file 5: Figure S5.**
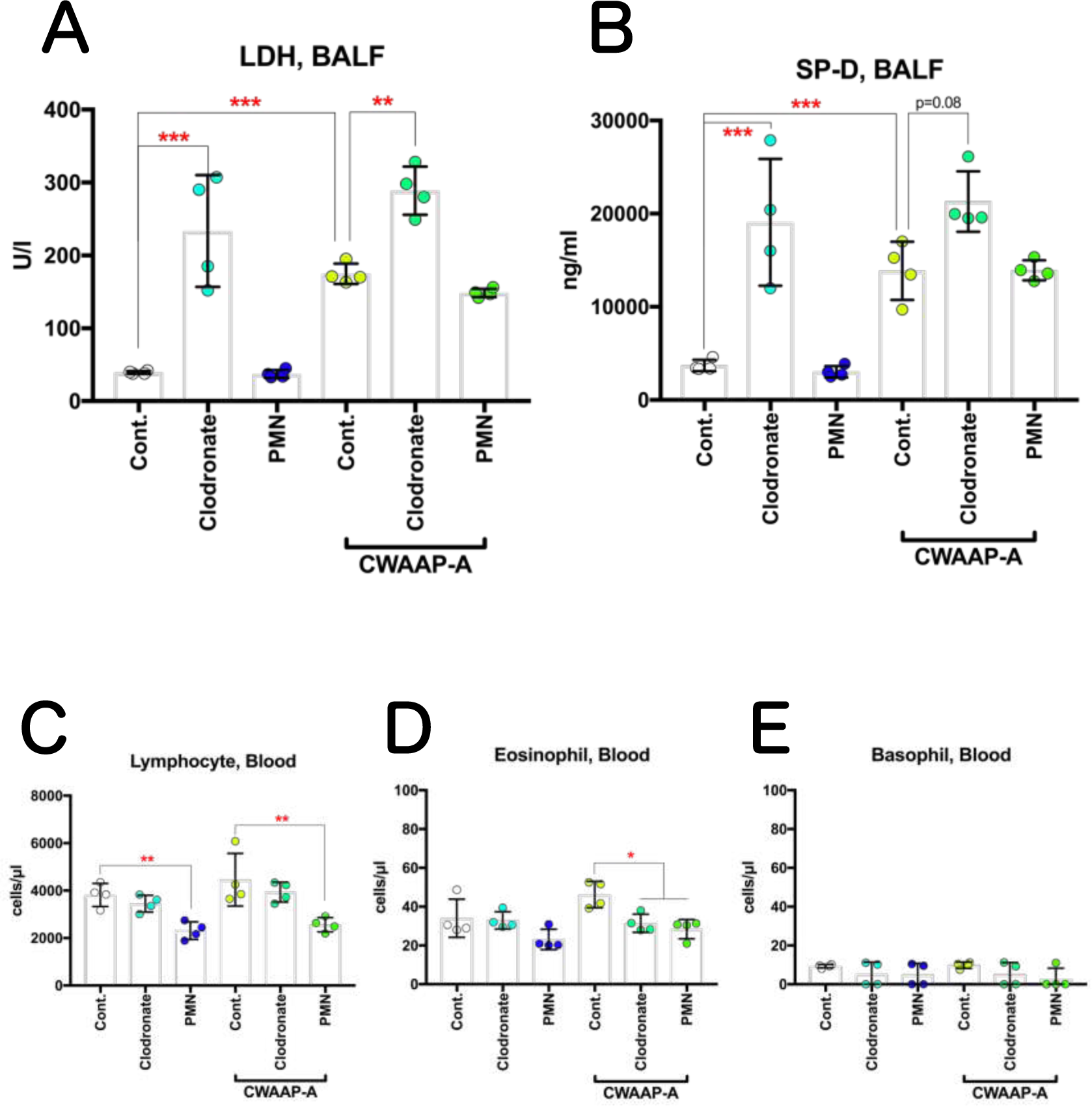
The effect of pretreatment of the CWAAP-A-exposed rats with clodronate liposomes (to (to deplete neutrophils) on BALF markers and cell populations. The lactate dehydrogenase (LDH) activity (A) and surfactant protein-D (SP-D) in BALF, and the number of lymphocytes (C), eosinophils (D) and basophils € in plasma were shown. Tukey’s multiple comparison test: \**p*<0.05, \*\**p*<0.01, and \*\*\**p*<0.001, pairs indicated.

**Additional file 6: Figure S6.**
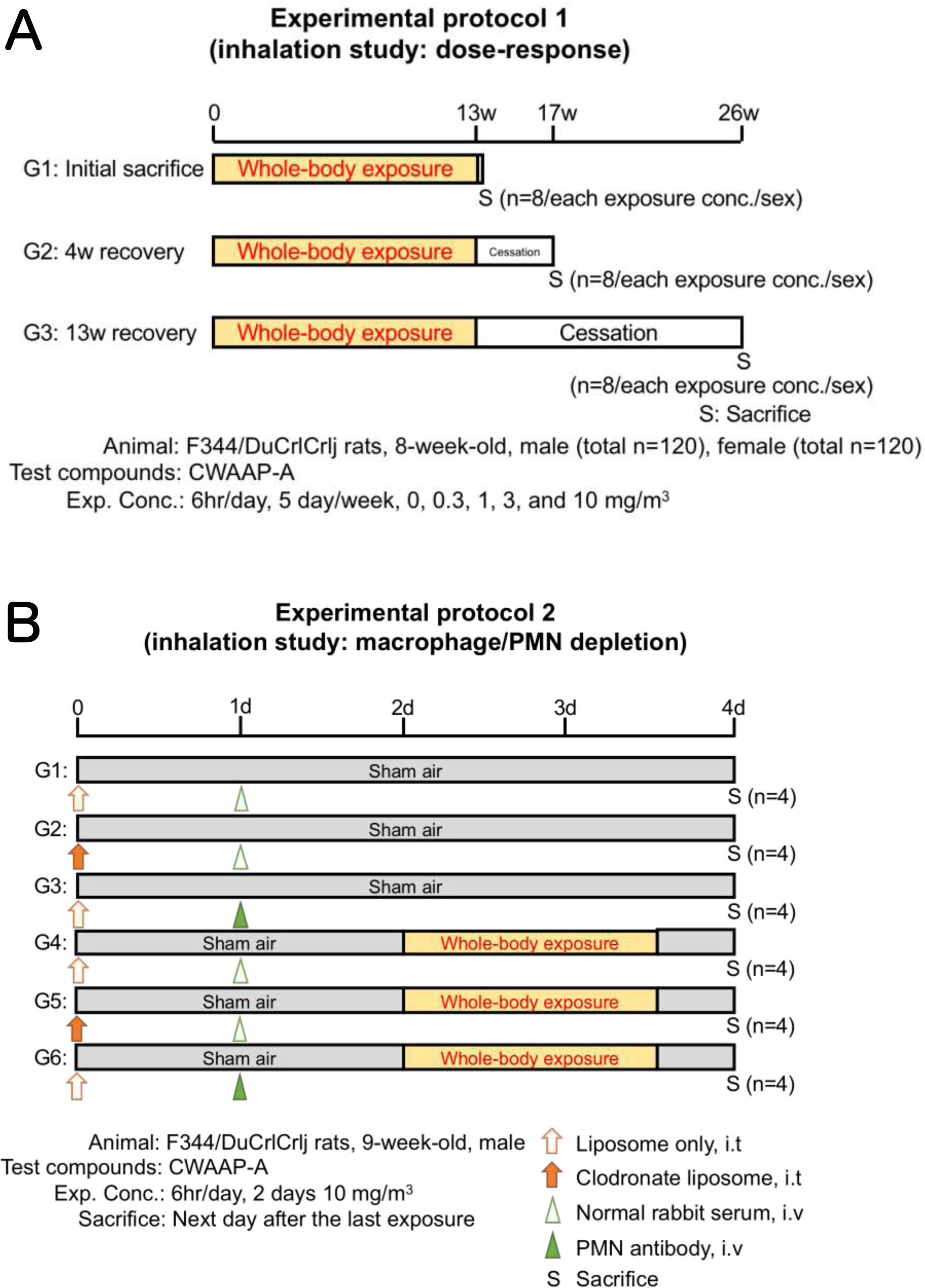
Design of animal experimental protocols in this research. 13-week inhalation exposure study (A), and macrophages or neutrophils depletion study (B).

**Additional file 7: Table S1.** The summary of the effect of inhalation exposure to CWAAP-A, multi-walled carbon nanotube (MWCNT)-7, and titanium dioxide (TiO_2_) on the LDH activity Abbreviation: NP, nanoparticles.

|  |  | chamber condition |  | LDH in BALF |  |
| --- | --- | --- | --- | --- | --- |
| Test compounds | 13-week study | MMAD<br>(GSD)<br>($\mu\text{m}$) | Exposure<br>concentration<br>($\text{mg}/\text{m}^3$) | average ± SD<br>(U/l) | Fold<br>change |
| CWAAP-A (in-house, in this study) |  |  |  |  |  |
| | male F344/DuCrI CrIj rat | | 0 | 25 ± 1.2 | - |
| | | 0.8 (2.7) | 0.3 | 62 ± 6.3 | 2.5 |
| | | 0.8 (2.6) | 1 | 177 ± 13.6 | 7.1 |
| | | 0.8 (2.6) | 3 | 319 ± 17.6 | 12.8 |
| | | 0.8 (2.7) | 10 | 684 ± 46.9 | 27.4 |
| | female F344/DuCrI CrIj rat | | 0 | 29 ± 1.0 | - |
| | | 0.8 (2.7) | 0.3 | 60 ± 4.1 | 2.0 |
| | | 0.8 (2.6) | 1 | 140 ± 16.6 | 4.8 |
| | | 0.8 (2.6) | 3 | 268 ± 10.4 | 9.2 |
| | | 0.8 (2.7) | 10 | 681 ± 83.0 | 23.3 |
| MWCNT (in-house, *Kasai et al.*) |  |  |  |  |  |
| | male F344/DuCrI CrIj rat | | 0 | 35 ± 2.6 | - |
| | | 1.5 (2.8) | 0.2 | 53 ± 4.2 | 1.5 |
| | | 1.5 (2.7) | 1 | 83 ± 6.5 | 2.4 |
| | | 1.5 (2.6) | 5 | 167 ± 4.6 | 4.8 |
| | female F344/DuCrI CrIj rat | | 0 | 37 ± 3.1 | - |
| | | 1.5 (2.8) | 0.2 | 51 ± 6.6 | 1.4 |
| | | 1.5 (2.7) | 1 | 81 ± 8.1 | 2.2 |
| | | 1.5 (2.6) | 5 | 165 ± 9.1 | 4.5 |
| TiO<sub>2</sub> NP, (in-house, Unpublished report, MHLW, Japan) |  |  |  |  |  |
| | male F344/DuCrI CrIj rat | | 0 | 24 ± 4.7 | - |
| | | 0.9 (2.1) | 6.3 | 28 ± 3.8 | 1.2 |
| | | 0.9 (2.1) | 12.5 | 30 ± 2.3 | 1.3 |
| | | 0.9 (2.1) | 25 | 33 ± 1.4 | 1.4 |
| | | 1.0 (2.1) | 50 | 60 ± 2.7 | 2.5 |
| | female F344/DuCrI CrIj rat | | 0 | 30 ± 2.8 | - |
| | | 0.9 (2.1) | 6.3 | 36 ± 2.4 | 1.2 |
| | | 0.9 (2.1) | 12.5 | 38 ± 4.7 | 1.2 |
| | | 0.9 (2.1) | 25 | 42 ± 2.1 | 1.4 |
| | | 1.0 (2.1) | 50 | 108 ± 16.2 | 3.6 |
| TiO<sub>2</sub>, ultrafine (*Bermudez et al.*) |  |  |  |  |  |
| | female CDF(F344)/CrIBR rat | | 0 | 24 ± 2.0 | - |
| | | 1.4 (2.6) | 0.5 | 26 ± 2.0 | 1.1 |
| | | 1.4 (2.6) | 2 | 29 ± 7.0 | 1.2 |
| | | 1.4 (2.6) | 10 | 122 ± 18.0 | 5.1 |

**Additional file 8: Table S2.** List of primary antibodies used in this study.

| 1st Antibodies | product company | ID | host | clonarity | dilution |
| --- | --- | --- | --- | --- | --- |
| αSMA | abcam | ab5694 | rabbit | poly | 1:250 |
| GFP | abcam | ab13970 | chicken | poly | 1:500 |
| VEGFR3 | R&D | AF743 | goat | poly | 1:100 |
| RT1-40 | terracebiotech | TB-11ART1-40 | mouse | mono | 1:100 |
| Lyve1 | ORIGENE | DP3513P | rabbit | poly | 1:100 |

**Additional file 9: Extended file 1.**
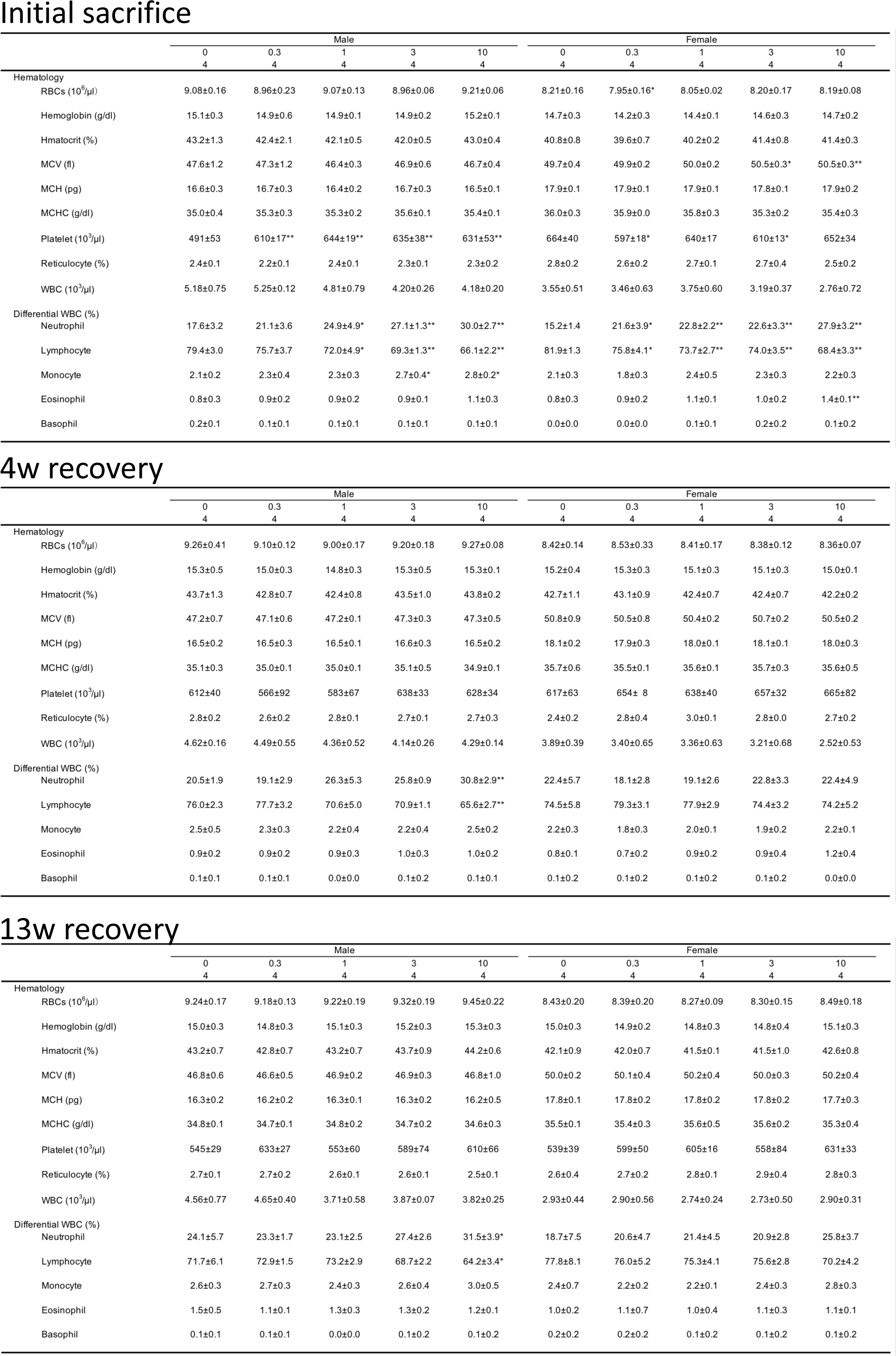
All blood-hematologic data observed in 13-week inhalation exposure study.

Extended file 1 Initial sacrifice
|  | Male |  |  |  |  | Female |  |  |  |  |
| --- | --- | --- | --- | --- | --- | --- | --- | --- | --- | --- |
|  | 0<br>4 | 0.3<br>4 | 1<br>4 | 3<br>4 | 10<br>4 | 0<br>4 | 0.3<br>4 | 1<br>4 | 3<br>4 | 10<br>4 |
| Hematology |  |  |  |  |  |  |  |  |  |  |
| RBCs (10<sup>6</sup>/μl) | 9.08±0.16 | 8.96±0.23 | 9.07±0.13 | 8.96±0.06 | 9.21±0.06 | 8.21±0.16 | 7.95±0.16* | 8.05±0.02 | 8.20±0.17 | 8.19±0.08 |
| Hemoglobin (g/dl) | 15.1±0.3 | 14.9±0.6 | 14.9±0.1 | 14.9±0.2 | 15.2±0.1 | 14.7±0.3 | 14.2±0.3 | 14.4±0.1 | 14.6±0.3 | 14.7±0.2 |
| Hmatocrit (%) | 43.2±1.3 | 42.4±2.1 | 42.1±0.5 | 42.0±0.5 | 43.0±0.4 | 40.8±0.8 | 39.6±0.7 | 40.2±0.2 | 41.4±0.8 | 41.4±0.3 |
| MCV (fl) | 47.6±1.2 | 47.3±1.2 | 46.4±0.3 | 46.9±0.6 | 46.7±0.4 | 49.7±0.4 | 49.9±0.2 | 50.0±0.2 | 50.5±0.3* | 50.5±0.3** |
| MCH (pg) | 16.6±0.3 | 16.7±0.3 | 16.4±0.2 | 16.7±0.3 | 16.5±0.1 | 17.9±0.1 | 17.9±0.1 | 17.9±0.1 | 17.8±0.1 | 17.9±0.2 |
| MCHC (g/dl) | 35.0±0.4 | 35.3±0.3 | 35.3±0.2 | 35.6±0.1 | 35.4±0.1 | 36.0±0.3 | 35.9±0.0 | 35.8±0.3 | 35.3±0.2 | 35.4±0.3 |
| Platelet (10<sup>3</sup>/μl) | 491±53 | 610±17** | 644±19** | 635±38** | 631±53** | 664±40 | 597±18* | 640±17 | 610±13* | 652±34 |
| Reticulocyte (%) | 2.4±0.1 | 2.2±0.1 | 2.4±0.1 | 2.3±0.1 | 2.3±0.2 | 2.8±0.2 | 2.6±0.2 | 2.7±0.1 | 2.7±0.4 | 2.5±0.2 |
| WBC (10<sup>3</sup>/μl) | 5.18±0.75 | 5.25±0.12 | 4.81±0.79 | 4.20±0.26 | 4.18±0.20 | 3.55±0.51 | 3.46±0.63 | 3.75±0.60 | 3.19±0.37 | 2.76±0.72 |
| Differential WBC (%) |  |  |  |  |  |  |  |  |  |  |
| Neutrophil | 17.6±3.2 | 21.1±3.6 | 24.9±4.9* | 27.1±1.3** | 30.0±2.7** | 15.2±1.4 | 21.6±3.9* | 22.8±2.2** | 22.6±3.3** | 27.9±3.2** |
| Lymphocyte | 79.4±3.0 | 75.7±3.7 | 72.0±4.9* | 69.3±1.3** | 66.1±2.2** | 81.9±1.3 | 75.8±4.1* | 73.7±2.7** | 74.0±3.5** | 68.4±3.3** |
| Monocyte | 2.1±0.2 | 2.3±0.4 | 2.3±0.3 | 2.7±0.4* | 2.8±0.2* | 2.1±0.3 | 1.8±0.3 | 2.4±0.5 | 2.3±0.3 | 2.2±0.3 |
| Eosinophil | 0.8±0.3 | 0.9±0.2 | 0.9±0.2 | 0.9±0.1 | 1.1±0.3 | 0.8±0.3 | 0.9±0.2 | 1.1±0.1 | 1.0±0.2 | 1.4±0.1** |
| Basophil | 0.2±0.1 | 0.1±0.1 | 0.1±0.1 | 0.1±0.1 | 0.1±0.1 | 0.0±0.0 | 0.0±0.0 | 0.1±0.1 | 0.2±0.2 | 0.1±0.2 |

4w recovery
|  | Male |  |  |  |  | Female |  |  |  |  |
| --- | --- | --- | --- | --- | --- | --- | --- | --- | --- | --- |
|  | 0<br>4 | 0.3<br>4 | 1<br>4 | 3<br>4 | 10<br>4 | 0<br>4 | 0.3<br>4 | 1<br>4 | 3<br>4 | 10<br>4 |
| Hematology |  |  |  |  |  |  |  |  |  |  |
| RBCs (10<sup>6</sup>/μl) | 9.26±0.41 | 9.10±0.12 | 9.00±0.17 | 9.20±0.18 | 9.27±0.08 | 8.42±0.14 | 8.53±0.33 | 8.41±0.17 | 8.38±0.12 | 8.36±0.07 |
| Hemoglobin (g/dl) | 15.3±0.5 | 15.0±0.3 | 14.8±0.3 | 15.3±0.5 | 15.3±0.1 | 15.2±0.4 | 15.3±0.3 | 15.1±0.3 | 15.1±0.3 | 15.0±0.1 |
| Hmatocrit (%) | 43.7±1.3 | 42.8±0.7 | 42.4±0.8 | 43.5±1.0 | 43.8±0.2 | 42.7±1.1 | 43.1±0.9 | 42.4±0.7 | 42.4±0.7 | 42.2±0.2 |
| MCV (fl) | 47.2±0.7 | 47.1±0.6 | 47.2±0.1 | 47.3±0.3 | 47.3±0.5 | 50.8±0.9 | 50.5±0.8 | 50.4±0.2 | 50.7±0.2 | 50.5±0.2 |
| MCH (pg) | 16.5±0.2 | 16.5±0.3 | 16.5±0.1 | 16.6±0.3 | 16.5±0.2 | 18.1±0.2 | 17.9±0.3 | 18.0±0.1 | 18.1±0.1 | 18.0±0.3 |
| MCHC (g/dl) | 35.1±0.3 | 35.0±0.1 | 35.0±0.1 | 35.1±0.5 | 34.9±0.1 | 35.7±0.6 | 35.5±0.1 | 35.6±0.1 | 35.7±0.3 | 35.6±0.5 |
| Platelet (10<sup>3</sup>/μl) | 612±40 | 566±92 | 583±67 | 638±33 | 628±34 | 617±63 | 654± 8 | 638±40 | 657±32 | 665±82 |
| Reticulocyte (%) | 2.8±0.2 | 2.6±0.2 | 2.8±0.1 | 2.7±0.1 | 2.7±0.3 | 2.4±0.2 | 2.8±0.4 | 3.0±0.1 | 2.8±0.0 | 2.7±0.2 |
| WBC (10<sup>3</sup>/μl) | 4.62±0.16 | 4.49±0.55 | 4.36±0.52 | 4.14±0.26 | 4.29±0.14 | 3.89±0.39 | 3.40±0.65 | 3.36±0.63 | 3.21±0.68 | 2.52±0.53 |
| Differential WBC (%) |  |  |  |  |  |  |  |  |  |  |
| Neutrophil | 20.5±1.9 | 19.1±2.9 | 26.3±5.3 | 25.8±0.9 | 30.8±2.9** | 22.4±5.7 | 18.1±2.8 | 19.1±2.6 | 22.8±3.3 | 22.4±4.9 |
| Lymphocyte | 76.0±2.3 | 77.7±3.2 | 70.6±5.0 | 70.9±1.1 | 65.6±2.7** | 74.5±5.8 | 79.3±3.1 | 77.9±2.9 | 74.4±3.2 | 74.2±5.2 |
| Monocyte | 2.5±0.5 | 2.3±0.3 | 2.2±0.4 | 2.2±0.4 | 2.5±0.2 | 2.2±0.3 | 1.8±0.3 | 2.0±0.1 | 1.9±0.2 | 2.2±0.1 |
| Eosinophil | 0.9±0.2 | 0.9±0.2 | 0.9±0.3 | 1.0±0.3 | 1.0±0.2 | 0.8±0.1 | 0.7±0.2 | 0.9±0.2 | 0.9±0.4 | 1.2±0.4 |
| Basophil | 0.1±0.1 | 0.1±0.1 | 0.0±0.0 | 0.1±0.2 | 0.1±0.1 | 0.1±0.2 | 0.1±0.2 | 0.1±0.2 | 0.1±0.2 | 0.0±0.0 |

13w recovery
|  | Male |  |  |  |  | Female |  |  |  |  |
| --- | --- | --- | --- | --- | --- | --- | --- | --- | --- | --- |
|  | 0<br>4 | 0.3<br>4 | 1<br>4 | 3<br>4 | 10<br>4 | 0<br>4 | 0.3<br>4 | 1<br>4 | 3<br>4 | 10<br>4 |
| Hematology |  |  |  |  |  |  |  |  |  |  |
| RBCs (10<sup>6</sup>/μl) | 9.24±0.17 | 9.18±0.13 | 9.22±0.19 | 9.32±0.19 | 9.45±0.22 | 8.43±0.20 | 8.39±0.20 | 8.27±0.09 | 8.30±0.15 | 8.49±0.18 |
| Hemoglobin (g/dl) | 15.0±0.3 | 14.8±0.3 | 15.1±0.3 | 15.2±0.3 | 15.3±0.3 | 15.0±0.3 | 14.9±0.2 | 14.8±0.3 | 14.8±0.4 | 15.1±0.3 |
| Hmatocrit (%) | 43.2±0.7 | 42.8±0.7 | 43.2±0.7 | 43.7±0.9 | 44.2±0.6 | 42.1±0.9 | 42.0±0.7 | 41.5±0.1 | 41.5±1.0 | 42.6±0.8 |
| MCV (fl) | 46.8±0.6 | 46.6±0.5 | 46.9±0.2 | 46.9±0.3 | 46.8±1.0 | 50.0±0.2 | 50.1±0.4 | 50.2±0.4 | 50.0±0.3 | 50.2±0.4 |
| MCH (pg) | 16.3±0.2 | 16.2±0.2 | 16.3±0.1 | 16.3±0.2 | 16.2±0.5 | 17.8±0.1 | 17.8±0.2 | 17.8±0.2 | 17.8±0.2 | 17.7±0.3 |
| MCHC (g/dl) | 34.8±0.1 | 34.7±0.1 | 34.8±0.2 | 34.7±0.2 | 34.6±0.3 | 35.5±0.1 | 35.4±0.3 | 35.6±0.5 | 35.6±0.2 | 35.3±0.4 |
| Platelet (10<sup>3</sup>/μl) | 545±29 | 633±27 | 553±60 | 589±74 | 610±66 | 539±39 | 599±50 | 605±16 | 558±84 | 631±33 |
| Reticulocyte (%) | 2.7±0.1 | 2.7±0.2 | 2.6±0.1 | 2.6±0.1 | 2.5±0.1 | 2.6±0.4 | 2.7±0.2 | 2.8±0.1 | 2.9±0.4 | 2.8±0.3 |
| WBC (10<sup>3</sup>/μl) | 4.56±0.77 | 4.65±0.40 | 3.71±0.58 | 3.87±0.07 | 3.82±0.25 | 2.93±0.44 | 2.90±0.56 | 2.74±0.24 | 2.73±0.50 | 2.90±0.31 |
| Differential WBC (%) |  |  |  |  |  |  |  |  |  |  |
| Neutrophil | 24.1±5.7 | 23.3±1.7 | 23.1±2.5 | 27.4±2.6 | 31.5±3.9* | 18.7±7.5 | 20.6±4.7 | 21.4±4.5 | 20.9±2.8 | 25.8±3.7 |
| Lymphocyte | 71.7±6.1 | 72.9±1.5 | 73.2±2.9 | 68.7±2.2 | 64.2±3.4* | 77.8±8.1 | 76.0±5.2 | 75.3±4.1 | 75.6±2.8 | 70.2±4.2 |
| Monocyte | 2.6±0.3 | 2.7±0.3 | 2.4±0.3 | 2.6±0.4 | 3.0±0.5 | 2.4±0.7 | 2.2±0.2 | 2.2±0.1 | 2.4±0.3 | 2.8±0.3 |
| Eosinophil | 1.5±0.5 | 1.1±0.1 | 1.3±0.3 | 1.3±0.2 | 1.2±0.1 | 1.0±0.2 | 1.1±0.7 | 1.0±0.4 | 1.1±0.3 | 1.1±0.1 |
| Basophil | 0.1±0.1 | 0.1±0.1 | 0.0±0.0 | 0.1±0.2 | 0.1±0.2 | 0.2±0.2 | 0.2±0.2 | 0.1±0.2 | 0.1±0.2 | 0.1±0.2 |

Initial sacrifice

**Additional file 10: Extended file 2.**
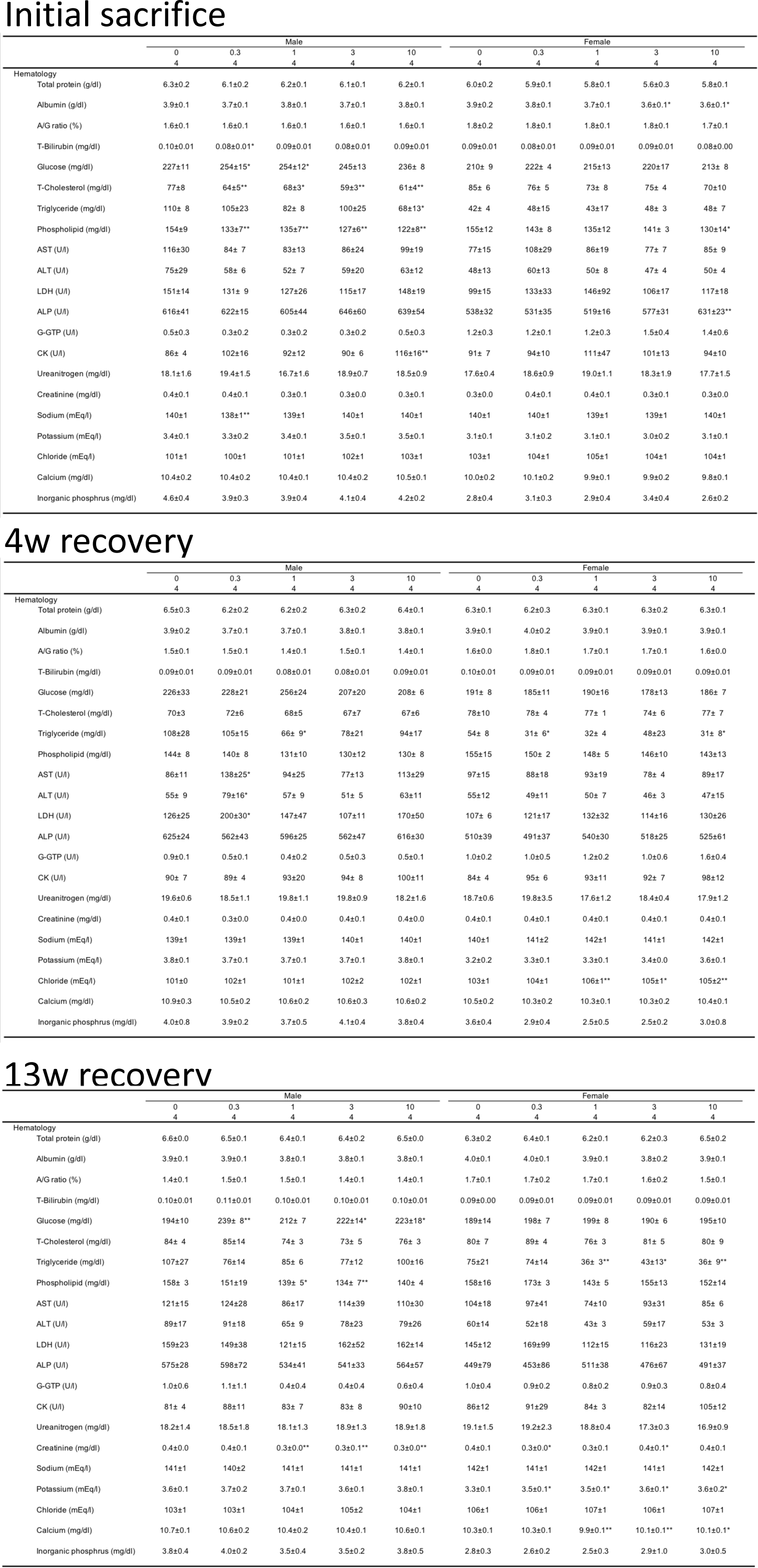
All blood-biochemistry data observed in 13-week inhalation exposure study.

**Additional file 11: Extended file 3.** Absolute/relative organ weights observed in 13-week inhalation exposure study.

Extended file 3-1 Initial sacrifice
|  |  | Male |  |  |  |  | Female |  |  |  |  |
| --- | --- | --- | --- | --- | --- | --- | --- | --- | --- | --- | --- |
|  |  | 0 | 0.3 | 1 | 3 | 10 | 0 | 0.3 | 1 | 3 | 10 |
|  |  | 4 | 4 | 4 | 4 | 4 | 4 | 4 | 4 | 4 | 4 |
| Organ (g) |  |  |  |  |  |  |  |  |  |  |  |
|  | Liver | 9.675±0.607 | 10.201±0.660 | 9.752±0.736 | 9.363±0.336 | 9.487±0.622 | 5.039±0.284 | 5.172±0.317 | 4.942±0.402 | 4.779±0.227 | 4.815±0.346 |
|  | Kidney | 1.884±0.031 | 2.084±0.138 | 1.991±0.127 | 1.964±0.105 | 1.980±0.095 | 1.225±0.035 | 1.243±0.086 | 1.221±0.061 | 1.206±0.045 | 1.186±0.072 |
|  | Spleen | 0.647±0.019 | 0.651±0.032 | 0.658±0.047 | 0.621±0.038 | 0.666±0.027 | 0.414±0.022 | 0.452±0.047 | 0.433±0.030 | 0.424±0.033 | 0.414±0.031 |
|  | Thymus | 0.257±0.047 | 0.254±0.016 | 0.256±0.028 | 0.235±0.021 | 0.256±0.024 | 0.204±0.013 | 0.203±0.020 | 0.193±0.022 | 0.193±0.030 | 0.192±0.011 |
|  | Heart | 0.927±0.054 | 0.962±0.038 | 0.915±0.085 | 0.910±0.069 | 0.970±0.029 | 0.592±0.054 | 0.612±0.040 | 0.581±0.023 | 0.577±0.044 | 0.574±0.033 |
|  | Testis/Ovary | 3.050±0.089 | 3.116±0.114 | 3.069±0.103 | 3.030±0.158 | 3.109±0.158 | 0.106±0.012 | 0.095±0.013 | 0.094±0.007 | 0.084±0.017 | 0.081±0.009 |
|  | Adrenal | 0.051±0.004 | 0.044±0.002 | 0.046±0.003 | 0.047±0.002 | 0.044±0.004 | 0.055±0.001 | 0.052±0.004 | 0.050±0.002 | 0.048±0.006 | 0.047±0.002 |
|  | Brain | 1.918±0.044 | 1.953±0.031 | 1.924±0.083 | 1.906±0.051 | 1.939±0.020 | 1.818±0.008 | 1.787±0.047 | 1.765±0.023 | 1.752±0.035 | 1.782±0.048 |

4w recovery
|  |  | Male |  |  |  |  | Female |  |  |  |  |
| --- | --- | --- | --- | --- | --- | --- | --- | --- | --- | --- | --- |
|  |  | 0 | 0.3 | 1 | 3 | 10 | 0 | 0.3 | 1 | 3 | 10 |
|  |  | 4 | 4 | 4 | 4 | 4 | 4 | 4 | 4 | 4 | 4 |
| Organ (g) |  |  |  |  |  |  |  |  |  |  |  |
|  | Liver | 10.036±0.447 | 9.900±0.511 | 10.218±0.308 | 9.634±0.831 | 9.857±0.484 | 5.282±0.097 | 4.924±0.209 | 4.911±0.319 | 4.833±0.368 | 4.934±0.120 |
|  | Kidney | 2.043±0.145 | 2.113±0.140 | 2.106±0.126 | 2.021±0.129 | 2.012±0.036 | 1.201±0.037 | 1.197±0.082 | 1.144±0.045 | 1.214±0.047 | 1.181±0.022 |
|  | Spleen | 0.677±0.033 | 0.699±0.044 | 0.690±0.037 | 0.670±0.052 | 0.687±0.013 | 0.428±0.020 | 0.442±0.042 | 0.440±0.032 | 0.408±0.029 | 0.411±0.020 |
|  | Thymus | 0.211±0.019 | 0.230±0.023 | 0.233±0.006 | 0.223±0.023 | 0.252±0.022 | 0.147±0.007 | 0.152±0.011 | 0.174±0.005 | 0.157±0.011 | 0.165±0.013 |
|  | Heart | 0.999±0.033 | 0.967±0.061 | 0.998±0.040 | 0.985±0.044 | 0.961±0.061 | 0.580±0.025 | 0.591±0.021 | 0.582±0.039 | 0.598±0.032 | 0.599±0.052 |
|  | Testis/Ovary | 3.127±0.163 | 3.134±0.044 | 3.212±0.031 | 3.122±0.083 | 3.104±0.101 | 0.106±0.012 | 0.108±0.010 | 0.110±0.008 | 0.101±0.006 | 0.106±0.007 |
|  | Adrenal | 0.051±0.001 | 0.046±0.003 | 0.052±0.008 | 0.047±0.005 | 0.043±0.001 | 0.052±0.002 | 0.051±0.003 | 0.048±0.003 | 0.047±0.005 | 0.045±0.003 |
|  | Brain | 1.946±0.007 | 1.952±0.061 | 1.976±0.035 | 1.950±0.044 | 1.945±0.021 | 1.789±0.021 | 1.792±0.036 | 1.760±0.056 | 1.809±0.030 | 1.794±0.035 |

13w recovery
|  |  | Male |  |  |  |  | Female |  |  |  |  |
| --- | --- | --- | --- | --- | --- | --- | --- | --- | --- | --- | --- |
|  |  | 0 | 0.3 | 1 | 3 | 10 | 0 | 0.3 | 1 | 3 | 10 |
|  |  | 4 | 4 | 4 | 4 | 4 | 4 | 4 | 4 | 4 | 4 |
| Organ (g) |  |  |  |  |  |  |  |  |  |  |  |
|  | Liver | 10.823±0.703 | 10.578±0.634 | 10.446±0.305 | 10.350±0.678 | 10.804±0.768 | 5.750±0.456 | 6.041±0.454 | 5.302±0.119 | 5.564±0.352 | 5.342±0.561 |
|  | Kidney | 2.188±0.091 | 2.307±0.123 | 2.288±0.071 | 2.215±0.147 | 2.251±0.097 | 1.273±0.086 | 1.419±0.069 | 1.319±0.050 | 1.364±0.069 | 1.299±0.088 |
|  | Spleen | 0.690±0.054 | 0.698±0.037 | 0.681±0.028 | 0.681±0.016 | 0.703±0.018 | 0.453±0.057 | 0.481±0.015 | 0.451±0.018 | 0.483±0.021 | 0.459±0.042 |
|  | Thymus | 0.224±0.017 | 0.230±0.018 | 0.220±0.012 | 0.215±0.024 | 0.229±0.034 | 0.137±0.012 | 0.159±0.010 | 0.145±0.005 | 0.154±0.021 | 0.157±0.020 |
|  | Heart | 1.027±0.057 | 1.088±0.023 | 1.075±0.061 | 1.010±0.059 | 1.076±0.065 | 0.627±0.078 | 0.716±0.063 | 0.624±0.026 | 0.641±0.047 | 0.641±0.052 |
|  | Testis/Ovary | 3.213±0.078 | 3.222±0.126 | 3.254±0.091 | 3.149±0.061 | 3.322±0.092 | 0.124±0.021 | 0.117±0.009 | 0.104±0.011 | 0.105±0.012 | 0.113±0.009 |
|  | Adrenal | 0.050±0.004 | 0.047±0.005 | 0.048±0.005 | 0.047±0.003 | 0.048±0.008 | 0.052±0.004 | 0.053±0.002 | 0.053±0.006 | 0.052±0.003 | 0.056±0.003 |
|  | Brain | 2.007±0.014 | 2.021±0.062 | 2.026±0.014 | 1.990±0.058 | 2.000±0.029 | 1.846±0.044 | 1.836±0.019 | 1.810±0.028 | 1.837±0.051 | 1.827±0.035 |

Initial sacrifice
| Organ (%) | Male |  |  |  |  | Female |  |  |  |  |
| --- | --- | --- | --- | --- | --- | --- | --- | --- | --- | --- |
|  | 0 | 0.3 | 1 | 3 | 10 | 0 | 0.3 | 1 | 3 | 10 |
|  | 4 | 4 | 4 | 4 | 4 | 4 | 4 | 4 | 4 | 4 |
| Liver | 3.080±0.093 | 3.123±0.086 | 3.072±0.082 | 3.019±0.087 | 2.930±0.019 | 2.750±0.165 | 2.782±0.059 | 2.802±0.202 | 2.787±0.103 | 2.785±0.094 |
| Kidney | 0.601±0.032 | 0.638±0.015 | 0.628±0.020 | 0.633±0.013 | 0.612±0.013 | 0.669±0.016 | 0.668±0.011 | 0.692±0.012 | 0.718±0.040 | 0.686±0.011 |
| Spleen | 0.207±0.014 | 0.199±0.003 | 0.207±0.008 | 0.200±0.005 | 0.206±0.011 | 0.226±0.013 | 0.243±0.008 | 0.245±0.009 | 0.247±0.015 | 0.240±0.009 |
| Thymus | 0.082±0.014 | 0.078±0.002 | 0.081±0.003 | 0.076±0.005 | 0.079±0.007 | 0.111±0.007 | 0.110±0.010 | 0.109±0.012 | 0.112±0.013 | 0.111±0.007 |
| Heart | 0.296±0.014 | 0.295±0.009 | 0.288±0.011 | 0.293±0.013 | 0.300±0.010 | 0.323±0.028 | 0.329±0.011 | 0.329±0.009 | 0.337±0.027 | 0.332±0.019 |
| Testis/Ovary | 0.972±0.027 | 0.956±0.058 | 0.969±0.046 | 0.977±0.047 | 0.962±0.060 | 0.058±0.007 | 0.051±0.006 | 0.053±0.006 | 0.049±0.008 | 0.047±0.005 |
| Adrenal | 0.016±0.001 | 0.014±0.001 | 0.015±0.001 | 0.015±0.001 | 0.014±0.001 | 0.030±0.001 | 0.028±0.002 | 0.028±0.001 | 0.028±0.004 | 0.027±0.002 |
| Brain | 0.611±0.021 | 0.599±0.026 | 0.608±0.035 | 0.615±0.026 | 0.600±0.034 | 0.992±0.006 | 0.963±0.048 | 1.003±0.057 | 1.024±0.082 | 1.033±0.046 |

4w recovery
| Organ (%) | Male |  |  |  |  | Female |  |  |  |  |
| --- | --- | --- | --- | --- | --- | --- | --- | --- | --- | --- |
|  | 0 | 0.3 | 1 | 3 | 10 | 0 | 0.3 | 1 | 3 | 10 |
|  | 4 | 4 | 4 | 4 | 4 | 4 | 4 | 4 | 4 | 4 |
| Liver | 2.962±0.056 | 2.860±0.087 | 2.964±0.028 | 2.878±0.105 | 2.860±0.114 | 2.926±0.016 | 2.740±0.069 | 2.732±0.108 | 2.744±0.149 | 2.730±0.056 |
| Kidney | 0.603±0.032 | 0.610±0.018 | 0.611±0.024 | 0.604±0.016 | 0.584±0.010 | 0.666±0.031 | 0.666±0.049 | 0.637±0.034 | 0.690±0.033 | 0.653±0.007 |
| Spleen | 0.200±0.005 | 0.202±0.005 | 0.200±0.006 | 0.200±0.010 | 0.200±0.009 | 0.237±0.012 | 0.245±0.019 | 0.245±0.007 | 0.232±0.013 | 0.228±0.006 |
| Thymus | 0.062±0.006 | 0.066±0.008 | 0.068±0.002 | 0.067±0.004 | 0.073±0.005 | 0.081±0.005 | 0.085±0.006 | 0.097±0.004 | 0.089±0.007 | 0.091±0.006 |
| Heart | 0.295±0.005 | 0.279±0.013 | 0.290±0.007 | 0.295±0.012 | 0.279±0.014 | 0.322±0.011 | 0.329±0.010 | 0.324±0.017 | 0.339±0.010 | 0.331±0.022 |
| Testis/Ovary | 0.923±0.033 | 0.907±0.043 | 0.932±0.016 | 0.936±0.055 | 0.901±0.042 | 0.059±0.008 | 0.060±0.006 | 0.061±0.003 | 0.058±0.002 | 0.059±0.004 |
| Adrenal | 0.015±0.001 | 0.014±0.001 | 0.015±0.003 | 0.014±0.002 | 0.013±0.001 | 0.029±0.001 | 0.028±0.002 | 0.027±0.002 | 0.027±0.002 | 0.025±0.001 |
| Brain | 0.575±0.015 | 0.564±0.011 | 0.573±0.016 | 0.585±0.044 | 0.565±0.014 | 0.992±0.025 | 0.998±0.028 | 0.981±0.050 | 1.029±0.035 | 0.993±0.008 |

13w recovery
| Organ (%) | Male |  |  |  |  | Female |  |  |  |  |
| --- | --- | --- | --- | --- | --- | --- | --- | --- | --- | --- |
|  | 0 | 0.3 | 1 | 3 | 10 | 0 | 0.3 | 1 | 3 | 10 |
|  | 4 | 4 | 4 | 4 | 4 | 4 | 4 | 4 | 4 | 4 |
| Liver | 2.894±0.101 | 2.805±0.092 | 2.871±0.070 | 2.853±0.106 | 2.874±0.062 | 2.968±0.040 | 2.982±0.168 | 2.804±0.134 | 2.817±0.029 | 2.745±0.195 |
| Kidney | 0.586±0.026 | 0.612±0.034 | 0.629±0.016 | 0.611±0.015 | 0.600±0.018 | 0.658±0.016 | 0.701±0.027 | 0.697±0.032 | 0.691±0.027 | 0.668±0.011 |
| Spleen | 0.184±0.010 | 0.185±0.005 | 0.188±0.008 | 0.188±0.010 | 0.188±0.015 | 0.233±0.011 | 0.238±0.014 | 0.239±0.009 | 0.245±0.010 | 0.236±0.006 |
| Thymus | 0.060±0.004 | 0.061±0.003 | 0.061±0.002 | 0.059±0.004 | 0.061±0.007 | 0.071±0.006 | 0.078±0.006 | 0.077±0.004 | 0.078±0.008 | 0.081±0.008 |
| Heart | 0.275±0.007 | 0.289±0.010 | 0.295±0.010 | 0.279±0.004 | 0.287±0.016 | 0.323±0.021 | 0.353±0.021 | 0.330±0.018 | 0.324±0.011 | 0.330±0.006 |
| Testis/Ovary | 0.860±0.020 | 0.855±0.039 | 0.894±0.027 | 0.870±0.051 | 0.887±0.064 | 0.064±0.007 | 0.058±0.006 | 0.055±0.006 | 0.054±0.005 | 0.058±0.007 |
| Adrenal | 0.014±0.001 | 0.013±0.002 | 0.013±0.001 | 0.013±0.001 | 0.013±0.003 | 0.027±0.002 | 0.026±0.001 | 0.028±0.004 | 0.026±0.001 | 0.029±0.002 |
| Brain | 0.538±0.016 | 0.536±0.013 | 0.557±0.019 | 0.550±0.030 | 0.534±0.036 | 0.956±0.054 | 0.908±0.034 | 0.957±0.040 | 0.932±0.049 | 0.942±0.061 |

Initial sacrifice

**Additional file 12: Extended file 4.**
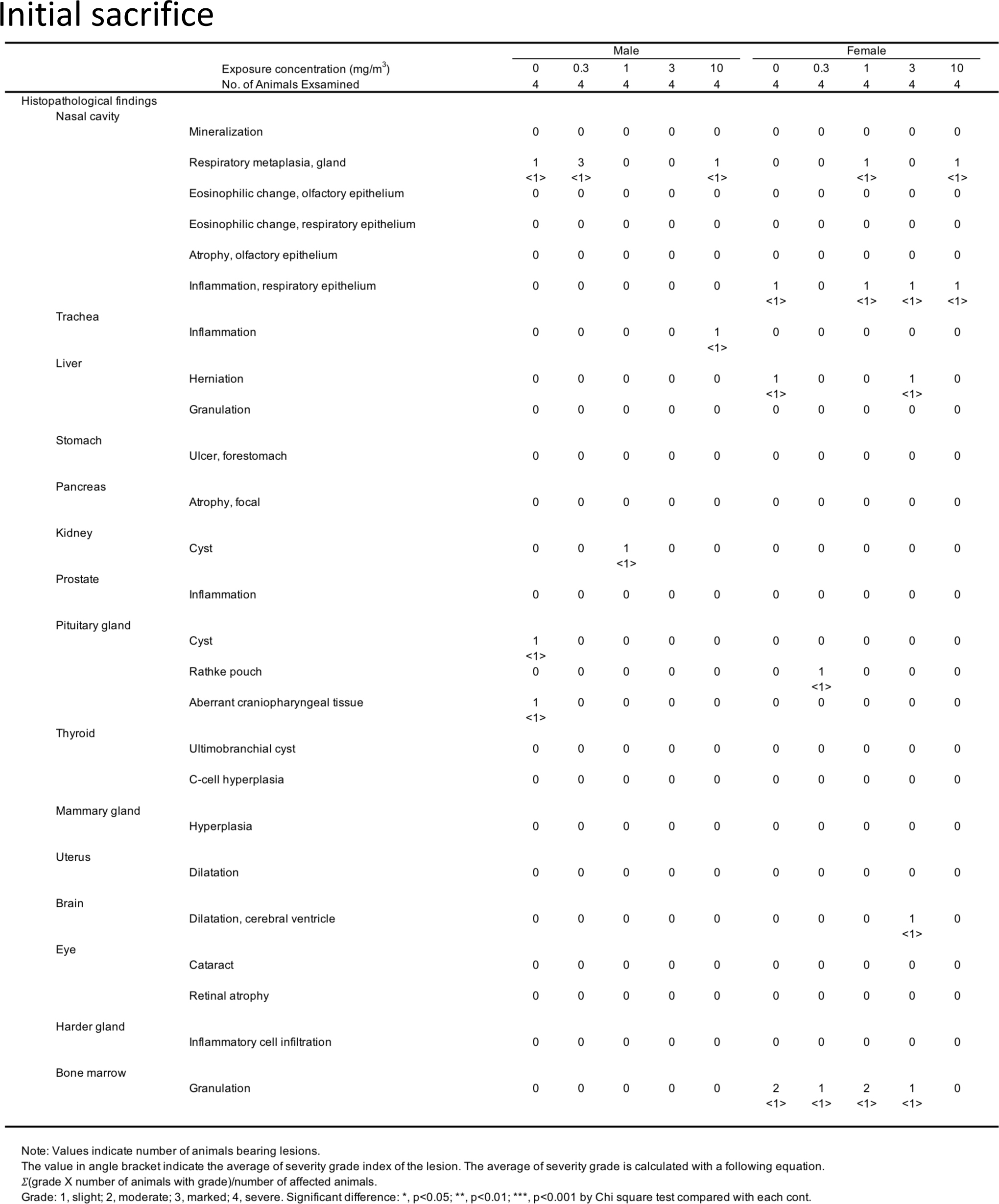
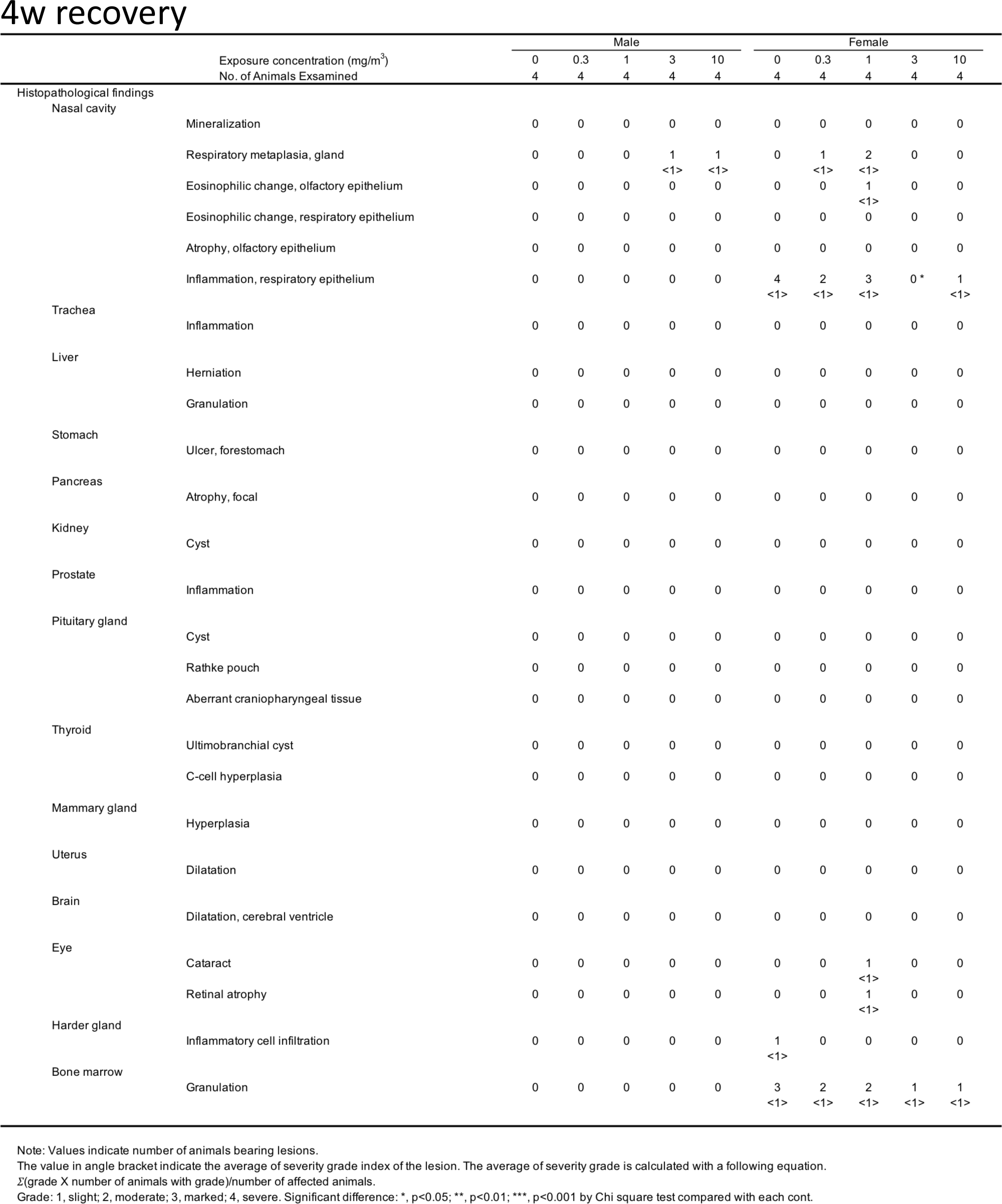
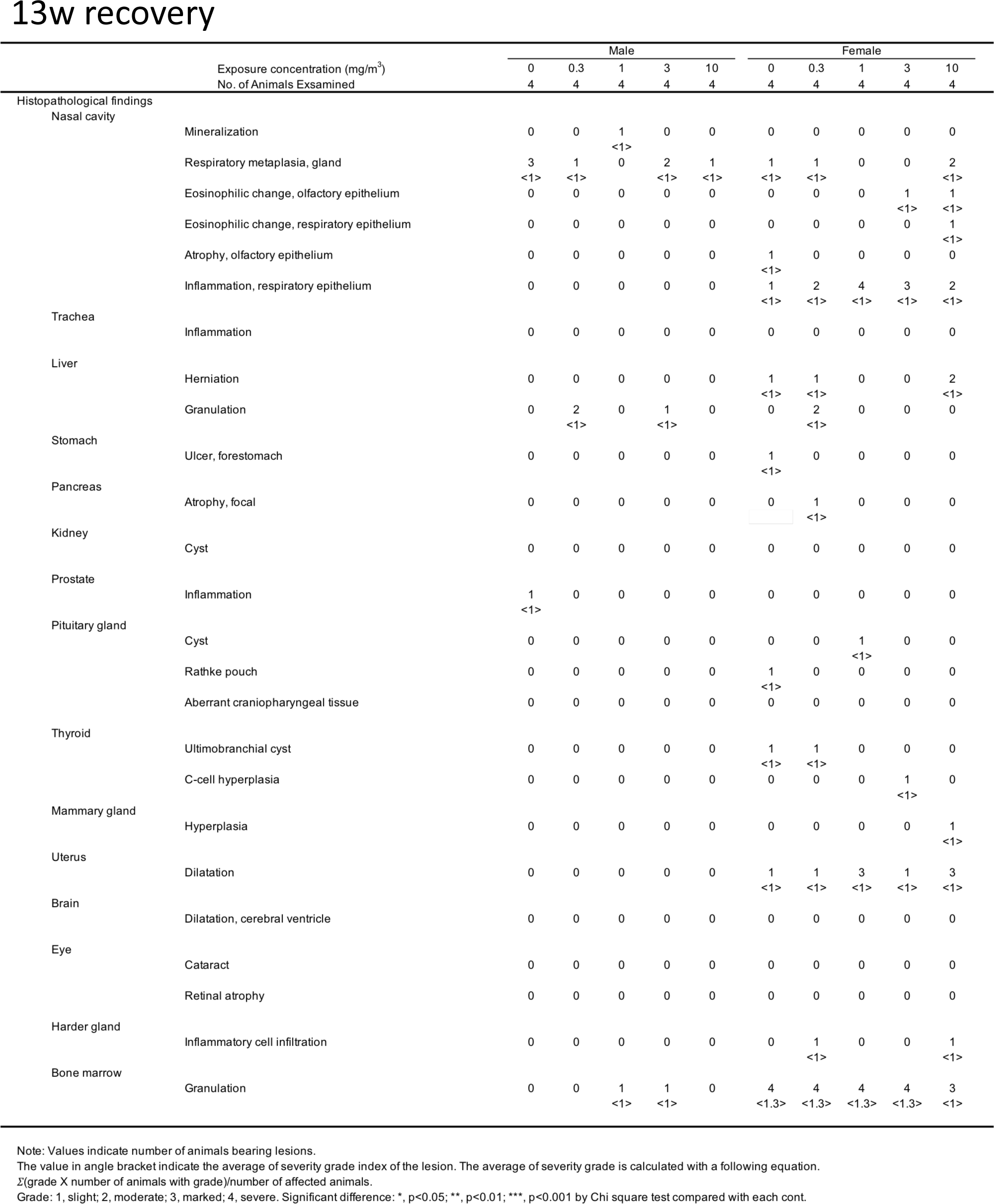
Histopathological findings excluding lung and mediastinal lymph node observed in 13-week inhalation exposure study.

**Additional file 13: Extended file 5.**
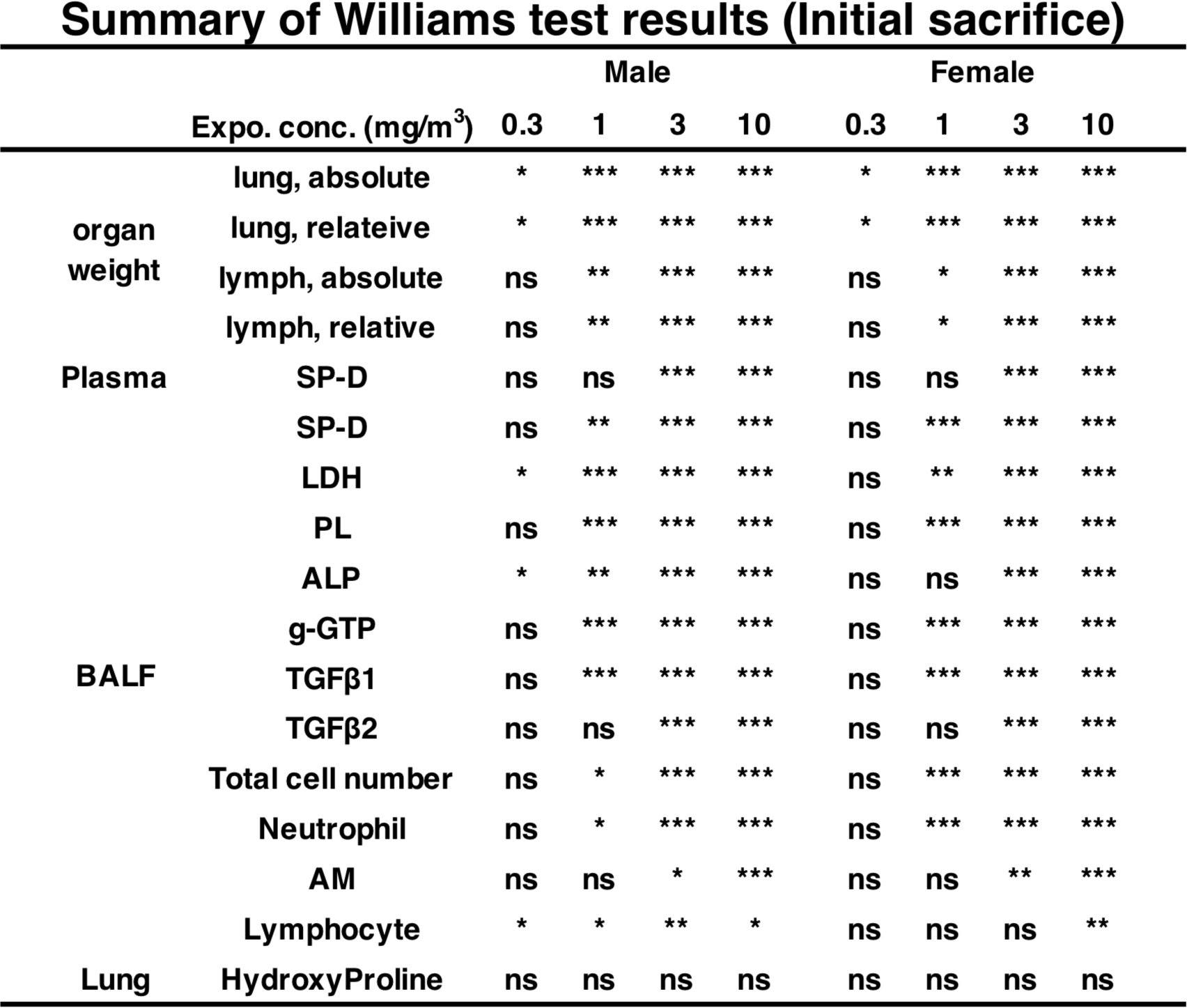
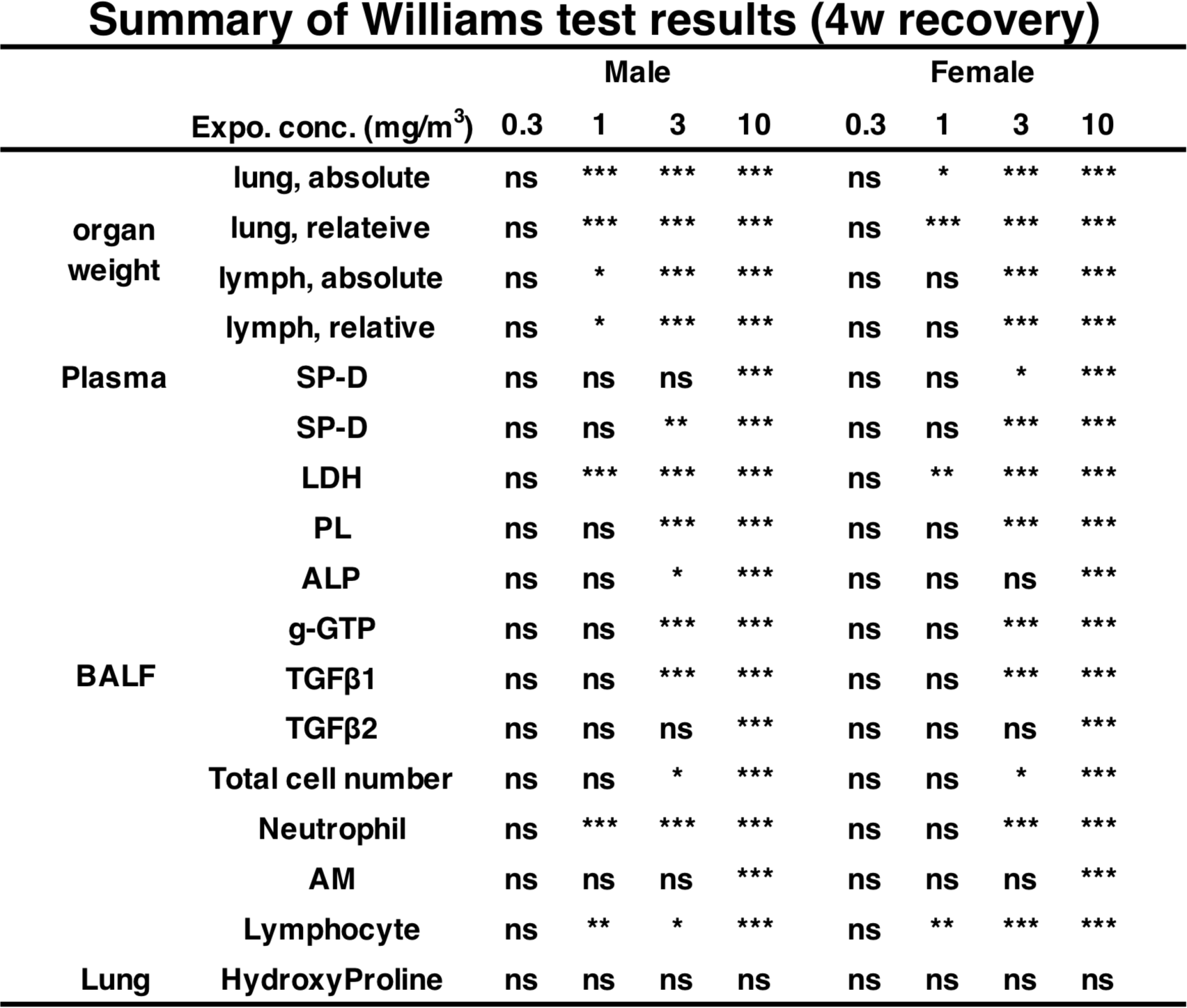
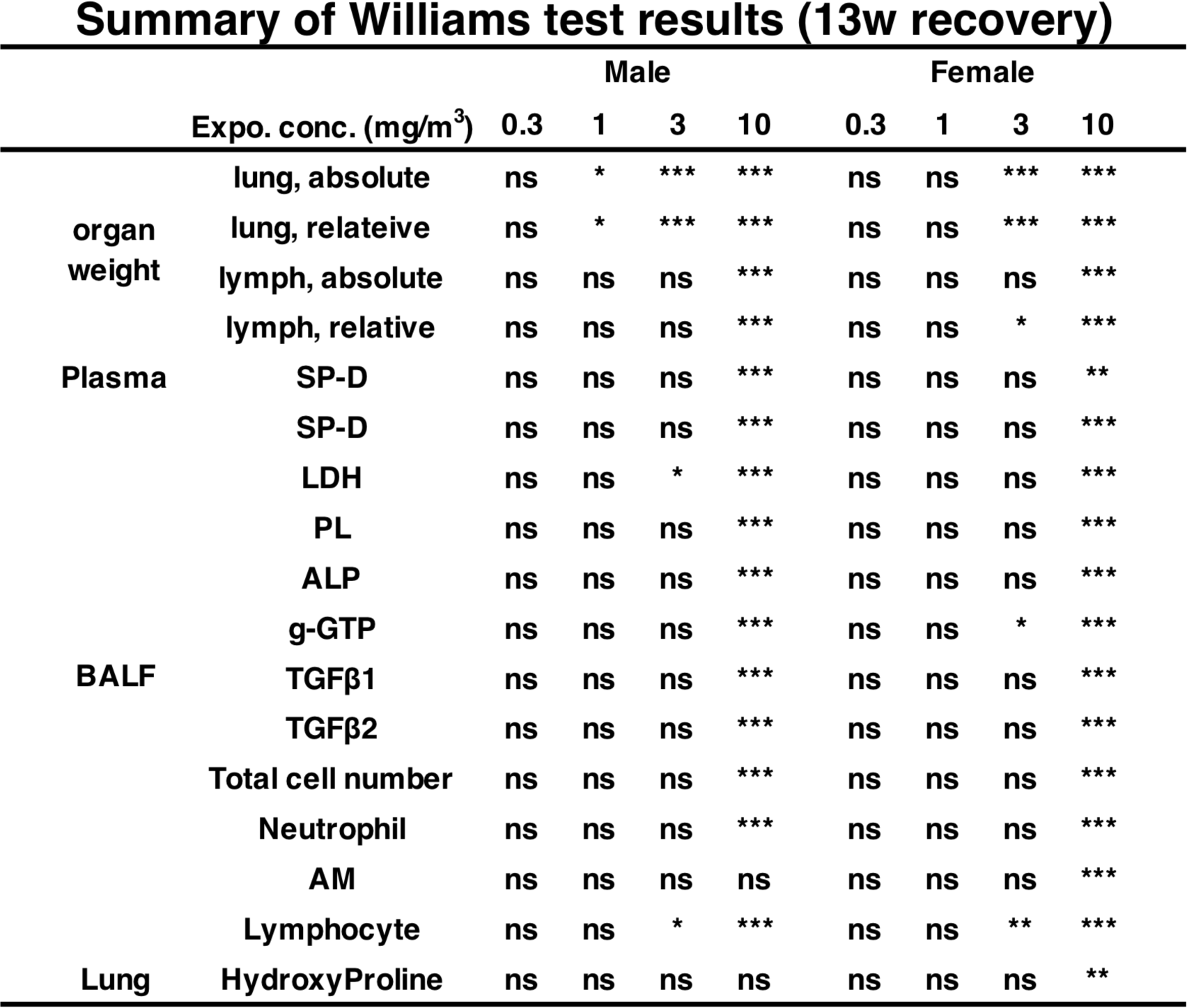
All summary data of Williams test results observed in 13-week inhalation exposure

## References

1. Alert notice about incidence of lung diseases among workers handling cross-linked water-soluble acrylic acid polymer in a chemical factory (published on April 29, 2017). [Internet]. Ministry of Health, Labour, and Welfare.; Available from: https://www.mhlw.go.jp/stf/houdou/0000163568.html.

2. Suka M, Kido T, Yoshioka W, Hachisuka E, Okoshi H, Yamauchi T, et al. Single intratracheal administration of cross-linked water-soluble acrylic acid polymer causes acute alveolo-interstitial inflammation and the subsequent fibrotic formation possibly via the TGF-β1 pathway in the lung of rats. Toxicology. 2021;448:152647.

3. Nishida C, Tomonaga T, Izumi H, Wang K-Y, Ishidao T, Takeshita J, et al. Inflammogenic Effect of Polyacrylic Acid in Rat Lung Following Intratracheal Instillation [Internet]. 2021 [cited 2021 Aug 7]. Available from: https://www.researchsquare.com/article/rs-147025/v1

4. Committee report on lung diseases among workers handling cross-linked water-soluble acrylic acid polymer in a chemical factory (published on April 19, 2019). [Internet]. Ministry of Health, Labour, and Welfare.; Available from: https://www.mhlw.go.jp/stf/shingi/other-roudou_128880_00004.html.

5. Yamano S, Takeda T, Goto Y, Hirai S, Furukawa Y, Kikuchi Y, et al. Pathological characteristics of pulmonary toxicity in F344 rats exposed by inhalation to cross-linked water-soluble acrylic acid polymers [Internet]. 2021 Nov p. 2021.11.13.468475. Available from: https://www.biorxiv.org/content/10.1101/2021.11.13.468475v1

6. Test No. 413: Subchronic Inhalation Toxicity: 90-day Study [Internet]. [cited 2021 Jul 17]. Available from: https://www.oecd-ilibrary.org/environment/test-no-413-subchronic-inhalation-toxicity-90-day-study_9789264070806-en

7. Naji A, Muzembo BA, Yagyu K-I, Baba N, Deschaseaux F, Sensebé L, et al. Endocytosis of indium-tin-oxide nanoparticles by macrophages provokes pyroptosis requiring NLRP3-ASC-Caspase1 axis that can be prevented by mesenchymal stem cells. Sci Rep. 2016;6:26162.

8. van Rooijen N, van Nieuwmegen R. Elimination of phagocytic cells in the spleen after intravenous injection of liposome-encapsulated dichloromethylene diphosphonate. An enzyme-histochemical study. Cell Tissue Res. 1984;238:355–8.

9. Sheridan BC, McIntyre RC, Agrafojo J, Meldrum DR, Meng X, Fullerton DA. Neutrophil depletion attenuates endotoxin-induced dysfunction of cGMP-mediated pulmonary vasorelaxation. Am J Physiol. 1996;271:L820–828.

10. Bustamante-Marin XM, Ostrowski LE. Cilia and Mucociliary Clearance. Cold Spring Harb Perspect Biol. 2017;9:a028241.

11. Wong BW, Zecchin A, García-Caballero M, Carmeliet P. Emerging Concepts in Organ-Specific Lymphatic Vessels and Metabolic Regulation of Lymphatic Development. Dev Cell. 2018;45:289–301.

12. Petrova TV, Koh GY. Biological functions of lymphatic vessels. Science. 2020;369.

13. Jung E, Gardner D, Choi D, Park E, Jin Seong Y, Yang S, et al. Development and Characterization of A Novel Prox1-EGFP Lymphatic and Schlemm’s Canal Reporter Rat. Sci Rep. 2017;7:5577.

14. Saito A, Horie M, Nagase T. TGF-β Signaling in Lung Health and Disease. Int J Mol Sci. 2018;19.

15. Meng X, Nikolic-Paterson DJ, Lan HY. TGF-β: the master regulator of fibrosis. Nat Rev Nephrol. 2016;12:325–38.

16. Takumi Kishimoto, Kenzo Okamoto, Shigeki Koda, Mariko Ono, Yumi Umeda, Shotaro Yamano, et al. Respiratory disease in workers handling cross-linked water-soluble acrylic acid polymer. J Occup Environ Med. [submitted]

17. Halappanavar S, van den Brule S, Nymark P, Gaté L, Seidel C, Valentino S, et al. Adverse outcome pathways as a tool for the design of testing strategies to support the safety assessment of emerging advanced materials at the nanoscale. Particle and Fibre Toxicology. 2020;17:16.

18. Maestrelli P, Boschetto P, Fabbri LM, Mapp CE. Mechanisms of occupational asthma. Journal of Allergy and Clinical Immunology. Elsevier; 2009;123:531–42.

19. Patel PH, Yarrarapu SNS, Anjum F. Byssinosis. StatPearls [Internet]. Treasure Island (FL): StatPearls Publishing; 2021 [cited 2021 Jul 25]. Available from: http://www.ncbi.nlm.nih.gov/books/NBK519549/

20. Zergham AS, Heller D. Farmers Lung. StatPearls [Internet]. Treasure Island (FL): StatPearls Publishing; 2021 [cited 2021 Jul 25]. Available from: http://www.ncbi.nlm.nih.gov/books/NBK557580/

21. Piirilä P, Kanerva L, Keskinen H, Estlander T, Hytönen M, Tuppurainen M, et al. Occupational respiratory hypersensitivity caused by preparations containing acrylates in dental personnel. Clin Exp Allergy. 1998;28:1404–11.

22. Savonius B, Keskinen H, Tuppurainen M, Kanerva L. Occupational respiratory disease caused by acrylates. Clin Exp Allergy. 1993;23:416–24.

23. Reig Rincón de Arellano I, Cimarra Alvarez-Lovell M, Robledo Echarren T, Fernández-Nieto M, Quirce Gancedo S, Seaone Plata C, et al. Occupational asthma due to acrylates in a graphic arts worker. Allergol Immunopathol (Madr). 2006;34:32–6.

24. Bochner BS, Undem BJ, Lichtenstein LM. Immunological Aspects of Allergic Asthma. Annu Rev Immunol. Annual Reviews; 1994;12:295–335.

25. Green RH, Brightling CE, McKenna S, Hargadon B, Parker D, Bradding P, et al. Asthma exacerbations and sputum eosinophil counts: a randomised controlled trial. Lancet. 2002;360:1715–21.

26. Price DB, Rigazio A, Campbell JD, Bleecker ER, Corrigan CJ, Thomas M, et al. Blood eosinophil count and prospective annual asthma disease burden: a UK cohort study. The Lancet Respiratory Medicine. Elsevier; 2015;3:849–58.

27. Nagano K, Nishizawa T, Eitaki Y, Ohnishi M, Noguchi T, Arito H, et al. Pulmonary toxicity in mice by 2- and 13-week inhalation exposures to indium-tin oxide and indium oxide aerosols. J Occup Health. 2011;53:234–9.

28. Nagano K, Gotoh K, Kasai T, Aiso S, Nishizawa T, Ohnishi M, et al. Two- and 13-week inhalation toxicities of indium-tin oxide and indium oxide in rats. J Occup Health. 2011;53:51–63.

29. Nagano K, Nishizawa T, Umeda Y, Kasai T, Noguchi T, Gotoh K, et al. Inhalation carcinogenicity and chronic toxicity of indium-tin oxide in rats and mice. J Occup Health. 2011;53:175–87.

30. Kasai T, Umeda Y, Ohnishi M, Kondo H, Takeuchi T, Aiso S, et al. Thirteen-week study of toxicity of fiber-like multi-walled carbon nanotubes with whole-body inhalation exposure in rats. Nanotoxicology. 2015;9:413–22.

31. Kasai T, Umeda Y, Ohnishi M, Mine T, Kondo H, Takeuchi T, et al. Lung carcinogenicity of inhaled multi-walled carbon nanotube in rats. Part Fibre Toxicol. 2016;13:53.

32. Chong S, Lee KS, Chung MJ, Han J, Kwon OJ, Kim TS. Pneumoconiosis: Comparison of Imaging and Pathologic Findings. RadioGraphics. Radiological Society of North America; 2006;26:59–77.

33. Adegunsoye A, Oldham JM, Bonham C, Hrusch C, Nolan P, Klejch W, et al. Prognosticating Outcomes in Interstitial Lung Disease by Mediastinal Lymph Node Assessment. An Observational Cohort Study with Independent Validation. Am J Respir Crit Care Med. 2019;199:747–59.

34. Baluk P, Naikawadi RP, Kim S, Rodriguez F, Choi D, Hong Y-K, et al. Lymphatic Proliferation Ameliorates Pulmonary Fibrosis after Lung Injury. Am J Pathol. 2020;190:2355–75.

35. Sorensen GL. Surfactant Protein D in Respiratory and Non-Respiratory Diseases. Front Med (Lausanne). 2018;5:18.

36. Beck-Schimmer B, Schwendener R, Pasch T, Reyes L, Booy C, Schimmer RC. Alveolar macrophages regulate neutrophil recruitment in endotoxin-induced lung injury. Respir Res. 2005;6:61.

37. Neupane AS. Understanding the function and behaviour of alveolar macrophages. Cumming School of Medicine; 2021 [cited 2021 Dec 8]; Available from: https://prism.ucalgary.ca/handle/1880/113378

38. Handa O, Naito Y, Yoshikawa T. Rat Cytokine-Induced Neutrophil Chemoattractant-1 (CINC-1) in Inflammation. Journal of Clinical Biochemistry and Nutrition. 2006;38:51–8.

39. Blackwell TS, Holden EP, Blackwell TR, DeLarco JE, Christman JW. Cytokine-induced neutrophil chemoattractant mediates neutrophilic alveolitis in rats: association with nuclear factor kappa B activation. Am J Respir Cell Mol Biol. 1994;11:464–72.

40. Mitsuhashi H, Hata J, Asano S, Kishimoto T. Appearance of cytokine-induced neutrophil chemoattractant isoforms and immunolocalization of them in lipopolysaccharide-induced acute lung inflammation in rats. Inflamm Res. 1999;48:588–93.

41. Yamasawa H, Ishii Y, Kitamura S. Cytokine-induced neutrophil chemoattractant in a rat model of lipopolysaccharide-induced acute lung injury. Inflammation. 1999;23:263–74.

42. Sumagin R. Emerging neutrophil plasticity: Terminally differentiated cells no more. J Leukoc Biol. 2021;109:473–5.

43. Boivin G, Faget J, Ancey P-B, Gkasti A, Mussard J, Engblom C, et al. Durable and controlled depletion of neutrophils in mice. Nat Commun. 2020;11:2762.

44. Bermudez E, Mangum JB, Wong BA, Asgharian B, Hext PM, Warheit DB, et al. Pulmonary Responses of Mice, Rats, and Hamsters to Subchronic Inhalation of Ultrafine Titanium Dioxide Particles. Toxicological Sciences. 2004;77:347–57.

45. Kasai T, Gotoh K, Nishizawa T, Sasaki T, Katagiri T, Umeda Y, et al. Development of a new multi-walled carbon nanotube (MWCNT) aerosol generation and exposure system and confirmation of suitability for conducting a single-exposure inhalation study of MWCNT in rats. Nanotoxicology. 2014;8:169–78.

46. 6-month subchronic inhalation toxicity study of W1009.03 in the rat. Procter and Gamble Company, Miami Valley Laboratories; 1987. Report No.: NO690-5900.

47. Renne R, Brix A, Harkema J, Herbert R, Kittel B, Lewis D, et al. Proliferative and nonproliferative lesions of the rat and mouse respiratory tract. Toxicol Pathol. 2009;37:5S–73S.

